# Whole genome analysis sheds light on the genetic origin of Huns, Avars and conquering Hungarians

**DOI:** 10.1101/2022.01.19.476915

**Authors:** Zoltán Maróti, Endre Neparáczki, Oszkár Schütz, Kitti Maár, Gergely I. B. Varga, Bence Kovács, Tibor Kalmár, Emil Nyerki, István Nagy, Dóra Latinovics, Balázs Tihanyi, Antónia Marcsik, György Pálfi, Zsolt Bernert, Zsolt Gallina, Ciprián Horváth, Sándor Varga, László Költő, István Raskó, Péter L. Nagy, Csilla Balogh, Albert Zink, Frank Maixner, Anders Götherström, Robert George, Csaba Szalontai, Gergely Szenthe, Erwin Gáll, Attila P. Kiss, Zsófia Rácz, Bence Gulyás, Bernadett Ny. Kovacsóczy, Szilárd Sándor Gaál, Péter Tomka, Tibor Török

## Abstract

Huns, Avars and conquering Hungarians were Migration Period nomadic groups which arrived in three successive waves in the Carpathian Basin between the 5^th^ and 9^th^ centuries. Based on historical data each of these groups are thought to have arrived from Asia, although their exact origin and relation to other ancient and modern populations has been debated. In this study we have sequenced 9 Hun, 143 Avar and 113 Hungarian conquest period samples, and identified three core populations, representing immigrants from each period, with no recent European ancestry. Our results suggest that this “immigrant core” of both Huns and Avars originated in present day Mongolia, and their origin can be traced back to Xiongnus. On the other hand, the “immigrant core” of the conquering Hungarians derived from an earlier admixture of Mansis, early Sarmatians and descendants of late Xiongnus. In addition, we detected shared Hun-related ancestry in numerous Avar and Hungarian conquest period genetic outliers indicating a genetic link between these successive nomadic groups. Aside from the immigrant core groups we identified that the majority of the individuals from each period were local residents, harboring “native European” ancestry.

## Background

Successive waves of population migrations associated with the Huns, Avars and Hungarians or Magyars from Asia to Europe had enduring impact on the population of the Carpathian Basin. This is most conspicuous in the unique language and ethno-cultural traditions of the Hungarians, whose closest parallels are found in populations east of the Urals. According to present scientific consensus these eastern links are solely attributed to the last migrating wave of conquering Hungarians (hence shortened as Conquerors), who arrived in the Carpathian Basin at the end of the 9^th^ century CE. On the other hand medieval Hungarian chronicles, foreign written sources and Hungarian folk traditions maintain that the origin of Hungarians can be traced back to the European Huns, with subsequent waves of Avars and Conquerors considered as kinfolks of the Huns^1,2^.

Both Huns and Avars founded a multiethnic empire in Eastern Europe, centered on the Carpathian Basin. The appearance of Huns in European written sources ca 370 CE was preceded by the disappearance of Xiongnus (Asian Huns) from Chinese sources. Likewise, the appearance of Avars in Europe in the sixth century, broadly correlates with the collapse of the Rouran Empire. However the possible relations between Xiongnus and Huns as well as Rourans and Avars remains largely controversial due to the scarcity of sources^3^.

From the 19th century onward, linguists reached a consensus that the Hungarian language is a member of the Uralic language family, belonging to the Ugric branch with its closest relatives, the Mansi and Khanty languages^4,5^. On this linguistic basis, the Hungarian prehistory was rewritten, and the Conquerors were regarded as descendants of a hypothetical Proto-Ugric people. At the same time, the formerly accepted Hun-Hungarian relations were called into question by source criticism of the medieval chronicles^6^.

Due to the scarceness of bridging literary evidence and the complex archaeological record, an archaeogenetic approach is best suited to provide insights into the origin and relationship of ancient populations. To this end, we performed whole genome analysis of European Hun, Avar and Conqueror period individuals from the Carpathian Basin, in order to shed light on the long debated origin of the European Huns, Avars and Conquerors. The majority of our 271 ancient samples (Supplementary Table 1a) were collected from the Great Hungarian Plain (Alföld), the westernmost extension of the Eurasian steppe, which provided favorable environment for the arriving waves of nomadic groups. The overview of archaeological sites and time periods of the studied samples is shown in Fig. 1, and a detailed archaeological description of the periods, cemeteries and individual samples is given in Supplementary Information. From the studied samples we report 73 direct AMS radiocarbon dates, of which 50 are first reported in this paper (Supplementary Table 2).

**Fig.1.**
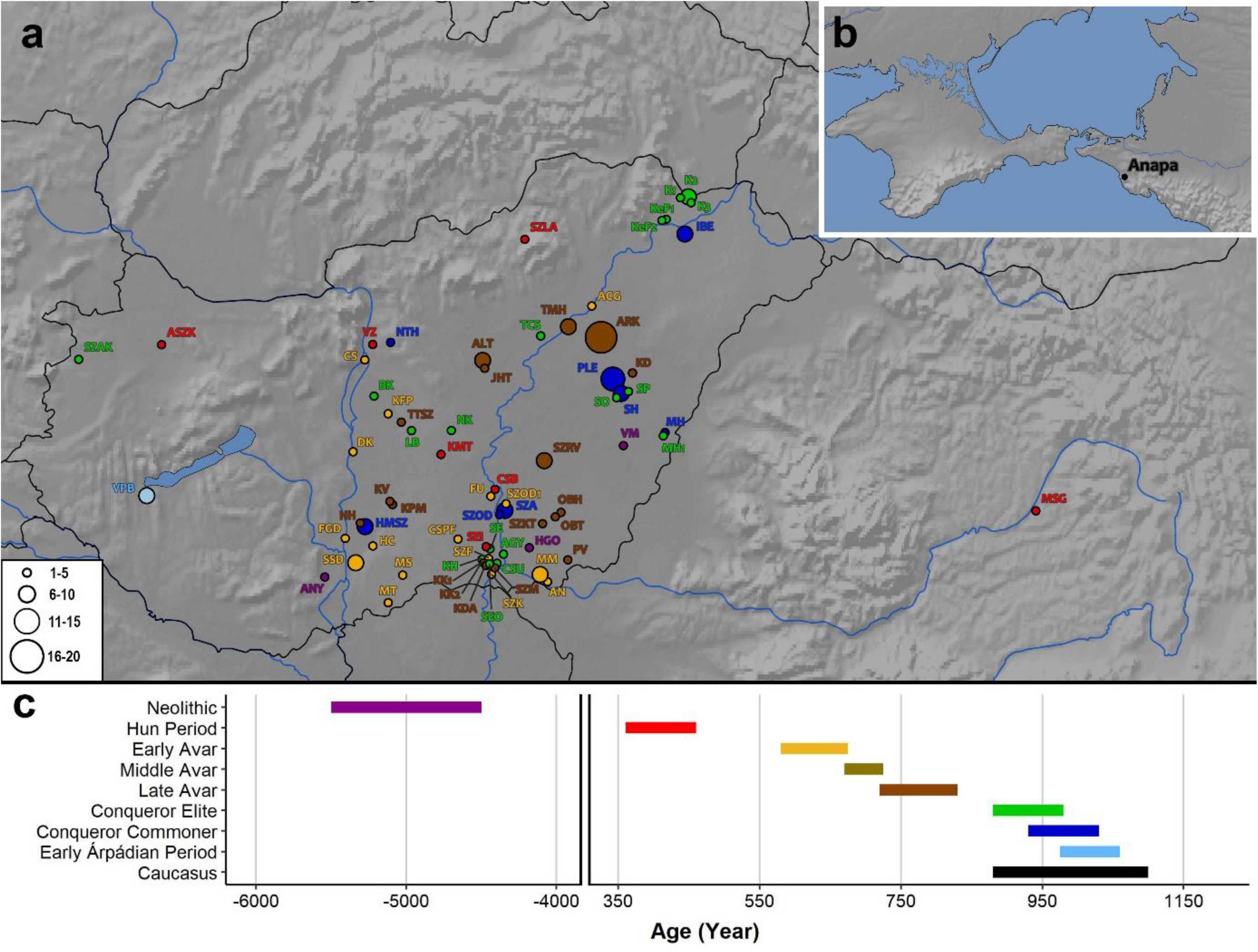
Archaeological sites and time periods of the studied samples. **a,** Distribution of sites with their associated culture and time period indicated by color. Color coding on panel **a,** corresponds to the time periods labelled with the same color in panel c, circle size is proportional to sample size, as indicated. **b,** Inset map of the Caucasus region, from where 3 samples were studied. **c,** Timeline of historical periods and corresponding archaeological cultures.

### Most individuals had local European ancestry

We performed principal component analysis (PCA), by projecting our ancient genomes onto the axes computed from modern Eurasian individuals (Fig. 2a and Extended Data Fig.1). On Fig. 2a most samples from each period project on with modern European populations, moreover these samples form a South-North cline along the P2 axes, which we termed the EU-cline. In order to group the most similar genomes, we clustered our samples together with all published ancient Eurasian genomes, according to their pairwise genetic distances obtained from the first 50 PCA dimensions (PC50 clustering, see Methods). As such, we identified five genetic clusters within the EU-cline (Fig. 2 and Supplementary Table 3), well sequestered along the P2 axes, which were named EU_Core1 to EU_Core5 respectively.

**Fig. 2.**
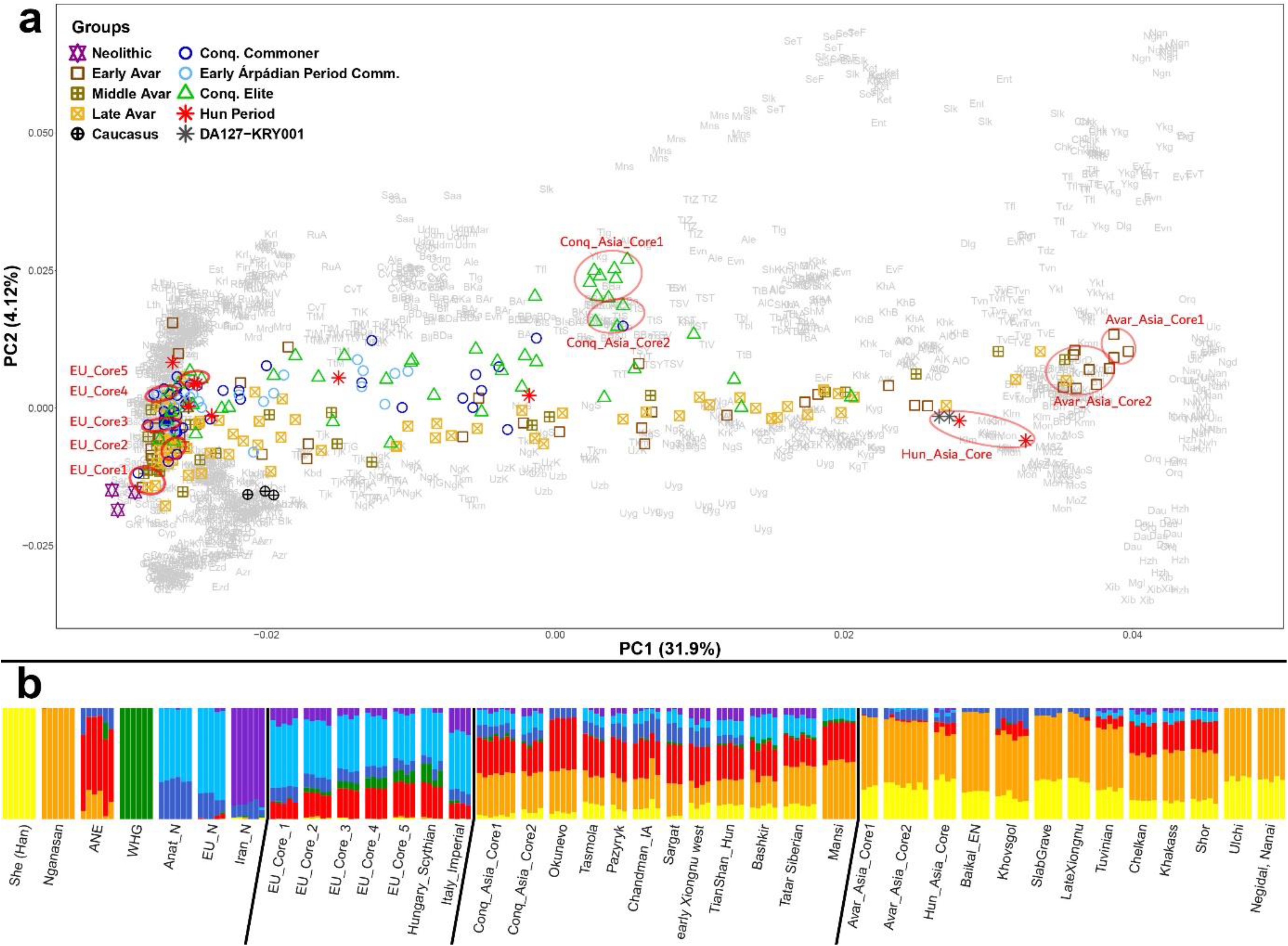
PCA and ADMIXTURE analysis. **a,** PCA of 271 ancient individuals projected onto contemporary Eurasians (gray latter codes, defined in Supplementary Table 8). Conquest and Avar period samples form two separable genetic clines. Genetically homogenous groups are encircled with red. **b,** Unsupervised ADMIXTURE (K=7) results of the red circled core groups, and the populations with most similar ADMIXTURE composition to these. The 7 populations maximizing each ADMIXTURE component are shown at the left

EU_Core1 clusters with Langobards from Hungary^7^, Iron Age, Imperial and Medieval individuals from Italy^8^, Minoans and Mycenaeans from Greece^9^ (Supplementary Table 3). EU_Core2, 3 and 4 cluster among others with Langobards^7^ and Bronze Age samples from Hungary^10,11^, the Czech Republic and Germany^11^, while EU_Core5 clusters with Hungarian Scythians^12^.

Unsupervised ADMIXTURE analysis revealed a gradient-like shift of genomic components along the EU-cline (Fig. 2b) with increasing Ancient North Eurasian (ANE) and Western Hunter-Gatherer (WHG) and decreasing early Iranian farmer (Iran_N) and early European farmer (EU_N) ancestries from South to North. It is also apparent that EU-cline samples contain negligible Asian (Nganasan and Han) components. ADMIXTURE also confirms that similar genomes had been present in Europe and the Carpathian Basin before the Migration Period, as EU_Core1 and 5 have comparable patterns to Imperial Period individuals from Italy and Iron Age Scythians from Hungary.

The diversity of the medieval Hungarian population in the EU-cline is conspicuous. We considered these groups local residents, although similar populations could hypothetically habe been present on the Medieval Pontic Steppe too.

### The European Huns had Xiongnu ancestry

Despite the paucity of Hun period samples, we can discern a “Hun-cline” along the PC1 axes (Fig 2a). Two individuals, MSG-1 and VZ-12673 (the same sample as HUN001^13^, resequenced with higher coverage) site at the extreme eastern pole of the cline, close to modern Kalmyks and Mongols. On PC50 clustering they tightly cluster together with two other Hun period samples; Kurayly_Hun_380CE (KRY001)^13^ and a Tian Shan Hun outlier (DA127)^12^ (Supplementary Table 3). As latter samples also form a genetic clade with VZ-12673 (see below) we grouped these four samples under the name of Hun_Asia_Core (Fig. 2), though analyzed the new samples separately. The Hun_Asia_Core also clusters with numerous Xiongnu, Medieval Mongol, Turk^14^ and Xianbei^13^ genomes from Mongolia as well as several Avar samples from this study. ADMIXTURE confirmed the similarity of Hun_Asia_Core individuals, and showed prevailing east Eurasian Nganasan and Han components with no traces of WHG (Fig. 2b and Supplementary Table 4), implying that these individuals represent immigrants with no European background.

Outgroup f3-statistics indicated common ancestry of MSG-1 and VZ-12673 (Extended Data Fig. 2), as both individuals shared highest drift with Neolithic farmers from the Wuzhuangguoliang site in northern China^15^, earlyXiongnu_rest, Ulaanzukh and SlabGrave samples from Mongolia^14^, pointing to a likely Mongolian origin and early Xiongnu affinity of these individuals.

Distal qpAdm modeling from pre-Iron Age sources indicated major (70-94%) Wuzhuangguoliang and minor (6-30%) Mongolian Bronze Age ancestries in MSG-1 and VZ-12673, while proximal modeling from post-Bronze Age sources gave two types of alternative models representing two different time periods (Supplementary Table 5a and 5b). The best P-value models showed major late Xiongnu (with Han admixture) and minor Scytho-Siberian/Xianbei ancestries, while alternative models indicated 78-100% Kazakhstan_OutTianShanHun or Kurayly_Hun_380CE and 0-12% Xiongnu/Xianbei/Han ancestries. In latter models VZ-12673 formed a clade with both published Hun_Asia_Core samples. In conclusion, our Hun_Asia_Core individuals could be equally modelled from earlier Xiongnu and later Hun age genomes.

The two other Hun period samples KMT-2785 and ASZK-1 were located in the middle of our PCA clines (Fig. 2 and Extended Data Fig 1a), and accordingly they could be modelled from European and Asian ancestors. The best passing models for KMT-2785 predicted 76% Late Xiongnu and 24% local EU_Core, while alternative model showed 86% Sarmatian^12^ and 14% Xiongnu ancestries (Supplementary Table 5c). Both models implicate Sarmatians as in the Late Xiongnus of the first model 46-52% Sarmatian and 48-54% Ulaanzuukh_SlabGrave components had been predicted^14^. The ASZK-1 genome formed a clade with Sarmatians in nearly all models. The rest of the Hun period samples map to the northern half of the EU cline, nevertheless two of these (SEI-1 and SEI-5) could be modelled from ~70% EU_Core and 30% Sarmatian components. The prevalent Sarmatian ancestry in 4 Hun period samples, implies significant Sarmatian influence on European Huns.

CSB-3 was modelled as ~80% EU_Core and 20% Scytho-Siberian, while SEI-6 formed a clade with the Ukraine_Chernyakhiv^16^ (Eastern Germanic/Goth) genomes. The SZLA-646 outlier individual at the top of the EU-cline formed a clade with Lithuania_Late_Antiquity^12^ and England_Saxon^17^ individuals. The last two individuals presumably also belonged to Germanic groups allied with the Huns.

Out of the 6 individuals in the Hun-cline (including DA127 and KRY001) four carried the R1a1a1b2 (R1a-Z93) Y-chromosomal haplogroup (Y-Hg), and one carried Q (Supplementary Table 1a), indicating that these Hg-s could be common among the European Huns, most likely inherited from Xiongnus^18^. Considering all published post-Xiongnu Hun era genomes^12,13^, we counted 10/23 R1a-Z93 and 9/23 Q Hgs, supporting this observation.

### Huns and Avars had related ancestry

Our Avar period samples also form a characteristic PCA “Avar-cline” on Fig. 2, extending from Europe to Asia. PC50 clustering identified a single genetic cluster at the Asian extreme of the cline with 12 samples, derived from 8 different cemeteries, which we termed Avar_Asia_Core (Fig. 2, Supplementary Table 3). 10/12 samples of Avar_Asia_Core were assigned to the early Avar period, 4 of them belonging to the elite, and 9/12 were males.

Avar_Asia_Core clusters together with Shamanka_Eneolithic and Lokomotiv_Eneolithic^19^ samples from the Baikal region, as well as with Mongolia_N_East, Mongolia_N_North^15^, Fofonovo_EN, Ulaanzuukh_SlabGrave and Xiongnu^14^ from Mongolia (Supplementary Table 3). This result is recapitulated in ADMIXTURE (Fig. 2b), which also shows that Nganasan and Han components predominate in Avar_Asia_Core with traces of Anat_N and ANE, while Iranian and WHG ingredients are entirely missing. It follows, that Avar_Asia_Core was derived from East Asia, most likely from present day Mongolia.

We performed two-dimensional f4-statistics to detect minor genetic differences within the Avar_Asia_Core group. Avar_Asia_Core individuals could be separated along a Bactria-Margiana Archaeological Complex (BMAC)-Steppe Middle-Late Bronze Age (Steppe_MLBA) cline (Extended Data Fig 3), with 3 individuals bearing negligible proportion of these ancestries. The Steppe_MLBA-ANE f4-statistics gave similar results. As the 3 individuals with the smallest Iranian, Steppe and ANE ancestries also visibly separated on PCA, we set apart these under the name of Avar_Asia_Core1, while the other 9 samples were regrouped as Avar_Asia_Core2 (Fig. 2).

According to outgroup f3-statistics both Avar_Asia_Core groups had highest shared drift with genomes having predominantly Ancient North-East Asian (ANA) ancestry (Extended Data Fig. 2), like earlyXiongnu_rest, Ulaanzuukh, and Slab Grave^14^. It is notable that from the populations with top 50 f3 values, 41 are shared with Hun_Asia_Core, moreover Avar_Asia_Core1 is the 16^th^ in the top list of VZ-12673 and 35^th^ in that of MSG-1, signifying common deep ancestry of European Huns and Avars.

According to distal qpAdm models Avar_Asia_Core formed a clade with the Fofonovo_EN and centralMongolia_preBA genomes (Supplementary Table 6a), both of which had been modelled from 83%–87% ANA and 12%–17% ANE^14^. All data consistently show that Avar_Asia_Core preserved very ancient Mongolian pre-Bronze Age genomes, with ~90% ANA ancestry.

Most proximodistal qpAdm models (defined in Methods) retained distal sources, as Avar_Asia_Core1 was modelled from 95% UstBelaya_N^20^ plus 5% Steppe Iron Age (Steppe_IA) and Avar_Asia_Core2 from 80-92% UstBelaya_N plus 8-20% Steppe_IA (Supplementary Table 6b). The exceptional proximal model for Avar_Asia_Core1 indicated 58% Yana_Medieval^20^ plus 42% Ulaanzukh, while for Avar_Asia_Core2 69% Xianbei_Hun_Berel^13^ plus 31% Kazakhstan_Nomad_Hun_Sarmatian^12^ ancestries. The latter model also points to shared ancestries between Huns and Avars.

From the 76 samples in the Avar-cline, 26 could be modelled as a simple 2-way admixture of Avar_Asia_Core and EU_Core (Supplementary Table 6c) indicating that these were admixed descendants of locals and immigrants, while further 9 samples required additional Hun and/or Iranian related sources. In the remaining 40 models Hun_Asia_Core and/or Xiongnu sources replaced Avar_Asia_Core (Supplementary Table 6d, summarized in Supplementary Table 1b). Scythian-related sources with significant Iranian ancestries, like Alan, Tian Shan Hun, Tian Shan Saka^12^, or Anapa (this study), were ubiquitous in the Avar-cline, but given their low proportion, qpAdm was unable to identify the exact source.

Xiongnu/Hun-related ancestries were more common in certain cemeteries, for example it was detected in most samples from Hortobágy-Árkus (ARK), Szegvár-Oromdűlő (SZOD), Makó-Mikócsa-halom (MM) and Szarvas-Grexa (SZRV). Y-chromosomal data seem to corroborate this conclusion, as 8/10 males from ARK carried Y-Hg Q, while 2/10 R1a-Z94, 3/3 males from SZRV carried R1a-Z94 and 2/2 males from MM carried Hg Q (Supplementary Table 1a).

### The Conquerors have Ugric, Sarmatian and Hun ancestry

The Conquest period samples also form a characteristic genetic “Conq-cline” on PCA (Fig. 2). It is positioned north of the he Avar-cline, whilst only reaching the midpoint of the P1 axis. PC50 clustering identified a single genetic cluster at the Asian extreme of the cline (Supplementary Table 3) with 12 samples, derived from 9 different cemeteries, which we termed Conq_Asia_Core. This genetic group consists of 6 males and 6 females and 11 of the 12 individuals belonged to the Conqueror elite according to archaeological evaluation.

The PCA position of Conq_Asia_Core corresponds to modern Bashkirs and Volga Tatars (Fig. 2a) and they cluster together with a wide range of eastern Scythians, western Xiongnus and Tian Shan Huns^12^, which is also supported by ADMIXTURE (Fig. 2b).

Two-dimensional f4-statistics detected slight genetic differences between Conq_Asia_Core individuals (Extended Data Fig 4), obtained via multiple gene flow, as they had different proportion of ancestry related to Miao (a modern Chinese group) and Ulaanzuukh_SlabGrave (ANA)^14^. Besides, individuals were arranged linearly along the Miao-ANA cline, suggesting that these ancestries covary in the Conqueror group, thus could have arrived together, most likely from present day Mongolia. As four individuals with highest Miao and ANA ancestries also had shifted PCA locations, we set these apart under the name of Conq_Asia_Core2, while the rest were regrouped as Conq_Asia_Core1 (Fig. 2).

Admixture f3-statistics indicated that the main admixture sources of Conq_Asia_Core1 were Steppe_MLBA populations and ancestors of modern Nganasans (Extended Data Fig. 5). Outgroup f3-statistics revealed that Conq_Asia_Core1 shared highest drift with modern Siberian populations speaking Uralic languages; Nganasan (Samoyedic), Mansi (Ugric), Selkup (Samoyedic) and Enets (Samoyedic) (Extended Data Fig. 5), implicating that Conq_Asia_Core shared evolutionary past with language relatives of modern Hungarians. For this reason we co-analyzed Mansis, the closest language relatives of Hungarians with Conq_Asia_Core.

From pre-Iron Age sources Mansis could be qpAdm modelled from 48% Mezhovskaya^10^, 44% Nganasan and 8% Botai^19^, while Conq_Asia_Core1 from 52% Mezhovskaya, 13% Nganasan, 20% Altai_MLBA_o^13^ and 15% Mongolia_LBA_CenterWest_4D^15^ (Supplementary Table 7a and 7b) confirming shared late Bronze Age ancestries of these groups, but also signifying that the Nganasan-like ancestry was largely replaced in Conq_Asia_Core by a Scytho-Siberian-like ancestry including BMAC^13,15^ derived from the Altai-Mongolia region.

From proximal sources Conq_Asia_Core1 could be consistently modelled from 50% Mansi, 35% Early/Late Sarmatian and 15% Scytho-Siberian-outlier/Xiongnu/Hun ancestries, and Conq_Asia_Core2 had comparable models with shifted proportions (Supplementary Table 7c). As the source populations in these models defined inconsistent time periods, we performed DATES analysis^21^ to clarify admixture time.

DATES revealed that the Mansi-Sarmatian admixture happened around 643-431 BCE, apparently corresponding to the early Sarmatian period, while the Mansi-Scythian/Hun admixture was dated around 217-315 CE, consistent with the post-Xiongnu, early Hun period rather than the Iron Age (Extended Data Fig 6).

Most individuals of the Conqueror cline proved to be admixed descendants of the immigrants and locals, as 31 samples could be modelled as two-way admixtures of Conq_Asia_Core and EU_Core (Supplementary Table 7d, summarized in Supplementary Table 1b).

The remaining samples mostly belonged to the elite, many projecting with the Avar-cline (Fig. 1), of which 5 could be modelled from Conq_Asia_Core with Hun and Iranian associated additional sources. 17 outlier individuals lacked Conq_Asia_Core ancestry, which was replaced with Avar_Asia_Core or Xiongnu/Hun-related sources, accompanied by Iranian associated 3^rd^ sources (Supplementary Table 7e). This result was again in line with Y-Hg data, as nearly all Conquest period males with R1a-Z94 or Q Hgs belonged to the last category (Supplementary Table 1a).

## Discussion

The genomic history of Huns Avars and Conquerors revealed in this study reconciles with historical, archaeological and linguistic sources (summarized in Fig. 3). Our data shows that the leader strata of both European Huns and Avars originated from the area of the former Xiongnu Empire, from present day Mongolia, and both groups can be traced back to early Xiongnu ancestors. Northern Xiongnus were expelled from Mongolia in the second century CE, and during their westward migration Sarmatians were one of the largest groups they confronted. Sergey Botalov presumed the formation of a Hun-Sarmatian mixed culture in the Ural region before the appearance of Huns in Europe^22^, which fits the significant Sarmatian ancestry detected in our Hun samples, though this ancestry had been present in late Xiongnus as well^14^. Thus our data are in accordance with the Xiongnu ancestry of European Huns, claimed by several historians^23,24^. We also detected Goth- or other German-type genomes among our Hun period samples, again consistent with historical sources^23^.

**Fig. 3.**
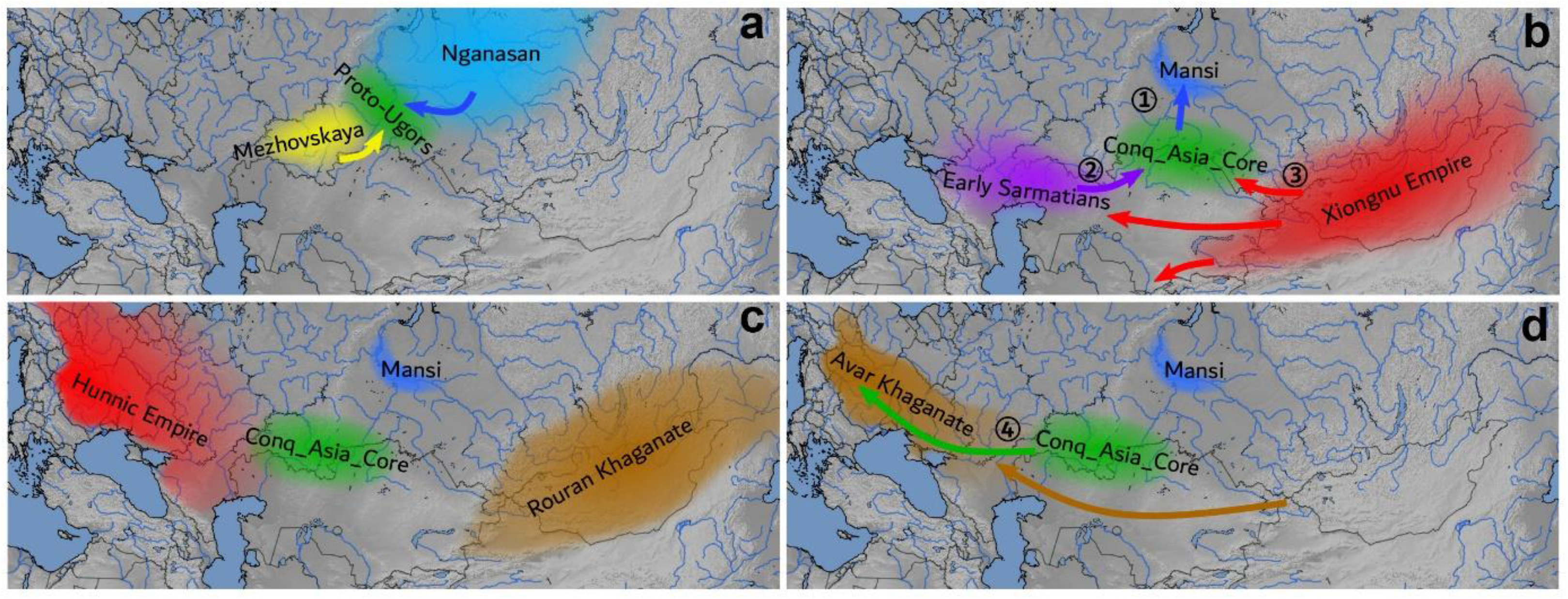
Summary map. **a,** Proto-Ugric peoples emerged from the admixture of Mezhovskaya and Nganasan populations in the late Bronze. **b,** 1. During the Iron Age Mansis separated. 2. proto-Conquerors admixed with Early Sarmatians 643-431 BCE and 3. with early Huns 217-315 CE. **c,** By the 5^th^ century the Xiongnu descent Hun Empire occupied Eastern Europe incorporating its population, and the Rouran Khaganate emerged on the former Xiongnu territory. **d,** By the middle 6^th^ century the Avar Khaganate occupied the territory of the former Hun Empire incorporating its populations. 4. By the 10^th^ century Conquerors associated with the remnants of both empires during their migration and within the Carpathian Basin.

Our data are compatible with the Rouran origin of Avar elite^25^, though the single low coverage Rouran genome^26^ provided a poor fit in the qpAm models (Supplementary Table 6b). The elite preserved very ancient east Asian genomes with undisputable origin, as had been also inferred from Y-Hg data^27,28^, however just half of the Avar-cline individuals had Avar_Asia_Core ancestry, implicating diverse origin of the Avar population. Our models indicate that the Avars incorporated groups with Xiongnu/Hun_Asia_Core and Iranian ancestries, presumably the remnants of the European Huns and Alans or other Iranian peoples on the Pontic Steppe, as suggested by Kim 2013^23^. People with different origin were seemingly distinguished, as samples with Hun-related genomes were buried in separate cemeteries.

The Conquerors, who arrived in the Carpathian Basin after the Avars, had distinct genomic background with elevated levels of western Eurasian admixture. They carried very similar genomes to modern Bashkirs and Tatars, in agreement with our previous results from uniparental markers^28,29^. Their genomes were shaped by several admixture events, of which the most fundamental was the Mezhovskaya-Nganasan admixture around the late Bronze Age, leading to the formation of a “proto-Ugric” gene pool. This was part of a general demographic process, when most Steppe_MLBA populations received an eastern Khovsgol related Siberian influx together with a BMAC influx^13^, and ANA related admixture became ubiquitous on the eastern Steppe^21^ establishing the Scytho-Siberian gene pool. Consequently proto-Ugric groups could be part of the early Scytho-Siberian societies of the late Bronze Age-early Iron Age steppe-forest zone in the northern Kazakhstan region, in the proximity of the Mezhovskaya territory.

Our data support linguistic models, which predicted that Conquerors and Mansis had a common early history^4,30^. Then Mansis migrated northward, probably during the Iron Age, and in isolation they preserved their Bronze-Age genomes. In contrast the Conquerors stayed at the steppe-forest zone and admixed with Iranian speaking early Sarmatians, also attested by the presence of Iranian loanwords in the Hungarian language^30^. This admixture likely happened when Sarmatians rose to power and started to integrate their neighboring tribes before they occupied the Pontic-Caspian Steppe.

All analysis congruently indicated, that the ancestors of Conquerors further admixed with a group from Mongolia, carrying Han-ANA related ancestry, which could be identified with early European Huns, compelling reconsideration of written historical sources about the Hun-Hungarian relations. It is to be examined, how this genetic link is related to reports in medieval Hungarian chronicles about the Hun ancestry of the Conqueror elite, which according to the current state of historiography is not sufficiently supported^31^. This admixture could happen before the Huns arrived to the Volga region and integrated local tribes east of the Urals, including Sarmatians and the ancestors of Conquerors. These data are compatible with a Conqueror homeland around the Ural region, in the vicinity of early Sarmatians, along the migration route of the Huns, as had been surmised from the phylogenetic connections between the Conquerors and individuals of the Kushnarenkovo-Karayakupovo culture in the Trans-Uralic Uyelgi cemetery^32^. Recently a Nganasan-like shared Siberian genetic ancestry was detected in all Uralic-speaking populations, Hungarians being an exception^33^. Our data fills this gap, as Conq_Asia_Core has high Nganasan ancestry, notwithstanding this is negligible in modern Hungarians, partly because of the substantially smaller number of immigrants compared to the local population.

The large number of genetic outliers with Hun_Asia_Core ancestry in both Avars and Conquerors testify that these successive nomadic groups were indeed assembled from overlaping populations.

## Materials and Methods

### Ethics statement

The human bone material used for ancient DNA analysis in this study were obtained from anthropological collections or museums, with the permission of the custodians in each case. In addition, we also contacted the archaeologists who excavated and described the samples, as well as the anthropologists who published anthropological details. In most cases these experts became co-authors of the paper, who provided the archaeological background in the Supplementary Information.

### Accelerator mass spectrometry radiocarbon dating

Here we report 73 radiocarbon dates, of which 50 are first reported in this paper. The sampled bone fragments were measured by accelerator mass spectrometry (AMS) in the AMS laboratory of the Institute for Nuclear Research, Hungarian Academy of Sciences, Debrecen, Hungary. Technical details concerning the sample preparation and measurement are given in^34^. Several radiocarbon measurements were done in the Radiocarbon AMS facility of the Center for Applied Isotope Studies, University of Georgia (n = 6;), technical details concerning the sample preparation and measurement are available here: https://cais.uga.edu/facilities/radiocarbon-ams-facility/). The conventional radiocarbon data were calibrated with the OxCal 4.4 software (https://c14.arch.ox.ac.uk/oxcal/OxCal.html, date of calibration: 4th of August 2021) with IntCal 20 settings^35^. Besides, we collected all previously published radiocarbon data related to the samples of our study.

### Ancient DNA laboratory work

All pre-PCR steps were carried out in the dedicated ancient DNA facilities of the Department of Genetics, University of Szeged and Department of Archaeogenetics, Institute of Hungarian Research, Hungary. Mitogenome or Y-chromosome data had been published from many of the samples used in this study^29,36^, and we sequenced whole genomes from the same libraries, whose preparations had been described in the above papers. For the rest of the samples we used the following modified protocol. DNA was extracted from bone powder collected from petrous bone or tooth cementum. 100 mg bone powder was predigested in 3 ml 0,5 M EDTA 100 μg/ml Proteinase K for 30 minutes at 48 °C, to increase the proportion of endogenous DNA. After pelleting, the powder was solubilized for 72 hours at 48 °C, in extraction buffer containing 0.45 M EDTA, 250 μg/ml Proteinase K and 1% Triton X-100. Then 12 ml binding buffer was added to the extract, containing 5 M GuHCl, 90 mM NaOAc, 40% isopropanol and 0,05% Tween-20, and DNA was purified on Qiagen MinElute columns.

Partial UDG treated libraries were prepared as described in^29^, but the preamplification step was omitted, and libraries were directly double indexed in one PCR-step after the adapter fill with Accuprime Pfx Supermix, containing 10mg/ml BSA and 200nM indexing P5 and P7 primers, in the following cycles: 95°C 5 minutes, 12 times 95°C 15 sec, 60°C 30 sec and 68°C 3 sec, followed by 5 minute extension at 68°C. The indexed libraries were purified on MinElute columns and eluted in 20μL EB buffer (Qiagen). Quantity measurements of the DNA extracts and libraries were performed with the Qubit fluorometric quantification system. The library fragment distribution was checked on TapeStation 2200 (Agilent). We estimated the endogenous human DNA content of each library with low coverage shotgun sequencing generated on iSeq 100 (Illumina) platform. Whole genome sequencing was performed on HiSeqX or NovaSeq 6000 Systems (Illumina) using paired-end sequencing method (2×150bp) following the manufacturer’s recommendations.

### Bioinformatics processing

Sequencing adapters were trimmed with the Cutadapt software [DOI:10.14806/ej.17.1.200] and sequences shorter than 25 nucleotides were removed. The raw reads were aligned to the GRCh37 (hs37d5) reference genome by Burrow-Wheels-Aligner (v 0.7.17)^37^, using the MEM command with reseeding disabled. To remove exogenous DNA only the primary alignments with >= 90% identity to reference were considered in all downstream analysis. In case of paired-end sequencing data we only kept properly paired primary alignments. Sequences from different lanes with their unique read groups were merged by samtools^38^. PICARD tools^39^ were used to mark duplicates. In case of paired-end reads we used the ATLAS software package^40^ mergeReads task with the options “updateQuality mergingMethod=keepRandomRead” to randomly exclude overlapping portions of paired-end reads, to mitigate potential random pseudo haploidization bias.

### Quality assessment of archaic sequences

Ancient DNA damage patterns were assessed using MapDamage 2.0^41^, and read quality scores were modified with the Rescale option to account for post-mortem damage. Mitochondrial contamination was estimated with the Schmutzi software package^42^. Contamination for the male samples was also assessed by the ANGSD X chromosome contamination method^43^, with the *“-r X:5000000-154900000 -doCounts 1 -iCounts 1 – minMapQ 30 -minQ 20 -setMinDepth 2*” options.

### Uniparental haplogroup assignment

Mitochondrial haplogroup determination was performed with the HaploGrep 2 (version 2.1.25) software^44^, using the consensus endogen fasta files resulting from the Schmutzi Bayesian algorithm. The Y haplogroup assessment was performed with the Yleaf software tool^45^, updated with the ISOGG2020 Y tree data set. Random pseudo haploid calling of whole genome samples were performed by the ANGSD software package (version: 0.931-10-g09a0fc5)^43^, using the „-*doHaploCall 1 -doCounts 1 -sites*” options and the Human Origins site coordinates, as well as the 1240K site coordinates of the Reich laboratory data sets.

### Genetic sex determination

Biological sex was assessed with the method described in^46^. Fragment length of paired-end data and average genome coverages (all, X, Y, mitochondrial) was assessed by the ATLAS software package^40^ using the BAMDiagnostics task. Detailed coverage distribution of autosomal, X, Y, mitochondrial chromosomes was calculated by the mosdepth software^47^.

### Estimation of genetic relatedness

Presence of close relatives in the dataset interferes with unsupervised ADMIXTURE and population genetic analysis, therefore we identified close kins and just one of them was left in the dataset. We performed kinship analysis using the 1240K data set and the PCAangsd software (version 0.931)^48^ from the ANGSD package with the “-inbreed 1 -kinship” options. We used the R (version 4.1.2); the RcppCNPy R package (version 0.2.10) to import the Numpy output files of PCAangsd.

### Population genetic analysis

The newly sequenced genomes were merged and co-analyzed with 2367 ancient (Supplementary Table 3) and 1397 modern Eurasian genomes (Supplementary Table 8), most of which were downloaded from the Allen Ancient DNA Resource (Version v42.4)^49^. We also downloaded the Human Origins dataset (HO, 6,2K SNP-s) and/or the 1240K data sets published in^13–15^. As the HO SNPs are fully contained in the larger 1240K set we filtered out the HO data when only the 1240K data set was published. In case the Reich data set contained preprint data of the same individual published later, we always used the published genotypes.

Since some data set contained diploid, and mixed call variants we performed random pseudo haploidization of all data prior to downstream analysis. Most of the analysis was done with the HO dataset, as most modern genomes are confined to this dataset, however, we run some of the f-statistics with the 1240K data if these were available

### Principal Component Analysis (PCA)

We used the modern Eurasian genome data published in Jeong et al. 2019^50^, confined to the HO dataset, to draw a modern PCA background on which ancient samples could be projected. However, in order to obtain the best separation of our samples in the PC1-PC2 dimensions, South-East Asian and Near Eastern populations were left out, and generally just 10 individuals were selected from each of the remaining populations, leaving 1397 modern individuals from 179 modern populations in the analysis (Supplementary Table 8). PCA Eigen vectors were calculated from 1397 pseudo-haploidized modern genomes with smartpca (EIGENSOFT version 7.2.1)^51^. Before projecting pseudo-haploidized ancient genomes, we excluded all relatives, and used the individuals with best genome coverage. All archaic genomes were projected on the modern background with the “lsqproject: YES” option. Since the archaic samples were projected, we used a more relaxed genotypization threshold (>50k genotyped markers) to exclude samples only where the results could be questionable due to the low coverage.

### Unsupervised Admixture

We carried out unsupervised admixture with 3277 genomes including 1010 modern and 2027 ancient published genomes plus 240 ones from present study, excluding all published relatives from each dataset. For this analysis we used the autosomal variants of the HO data set as many relevant modern populations are missing from the 1240K set. We set strict criteria for the selection of individual samples to minimize bias and maximize the information content of our data set. We excluded all samples with QUESTIONABLE flag or Ignore tag based on the annotation file of the data set to remove possibly contaminated samples and population outliers. To compose a balanced high quality data set furthermore we restricted the selection to maximum 10 individuals per populations and excluded all poorly genotyped samples (<150K genotyped markers).

To prepare the final marker set for ADMIXTURE analysis we removed variants with very low frequency (MAF <0.005) leaving 471625 autosomal variants. We pruned 116237 variants in linkage disequilibrium using PLINK with the options “--*indep-pairwise 200 10 0.25*” leaving a final 355388 markers for the 3277 individuals. The total genotyping rate of this high quality data set was 0.811831. We performed the unsupervised ADMIXTURE analysis for K=3-12 in 30 parallel runs with the ADMIXTURE software (version 1.3.0)^52^ and selected the lowest cross validation error model (K=7) for visualization.

### Hierarchical Ward clustering

Similar PCA position and Admixture composition may indicate population relations derived from shared genetic ancestry, but we further verified the close relation of similar looking genomes using Hierarchical Clustering (ward.D2)^53^ implemented in R 3.6.3^54^. Though the first two PCA Eigenvalues capture the strongest levels of variation in the data, in our analysis subsequent Eigenvalues had comparable magnitude to the second one, indicating that lower Eigenvectors sill harbored significant additional genetic information. Genetically most similar genomes are expected to occupy similar positions along multiple PCA dimensions, thus we clustered individuals according to their genetic distances obtained from the first 50 PCA dimensions (PC50 clustering). As hierarchical clustering is an ideal tool to arrange multidimensional data according to similarity, we clustered our 271 genomes according to the pairwise weighed Euclidean distances of their first 50 PCA Eigenvalues and Eigenvectors (PC50 distances), where distances were calculated as follows: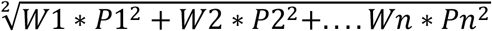 where *W*=Eigenvalue and *P*=Eigenvector. The hierarchical clusters based on the first PC50 distances are shown in (Supplementary Table 3).

In order to obtain genetically homogenous source populations for qpAdm, we also regrouped relevant published samples with the same method. For this end we projected 2367 relevant published Eurasian ancient samples on the same modern PCA background described above and performed the same Hierarchical Ward clustering based on the pairwise PC50 distances. This way we regrouped Late Bronze Age, Iron Age and Medieval samples published in ^10,12–16,55,56^ (Supplementary Table 3). If samples from published genetic groups fell into different PC50 clusters, we subdivided the original group according to the clusters. In some cases we distinguished subgroups even within the same PC50 clusters, if they were separated by relatively large PC1 or PC2 distances. We selected the best coverage samples as representatives for each group.

### Admixture modeling using qpAdm

We used qpAdm^57^ from the ADMIXTOOLS software package^58^ for modeling our genomes as admixtures of 2 or 3 source populations and estimating ancestry proportions. We set the details:YES parameter, to evaluate Z-scores for the goodness of fit of the model (estimated with a Block Jackknife).

As we tested a large number of source populations, testing every possible combination of sources (Left populations) and outgroups (Right populations) was impossible. Instead we run the analysis just with source combinations of 2 and 3 (rank 2 and 3). As qpWave is integrated in qpAdm, the nested-P values in the log files indicate the optimal rank of the model, that is if P-value for the nested model is above 0.05, the Rank-1 model should be considered^57^.

For revealing past population history of the Test populations from different time periods, we run two separate qpAdm analysis. In the so called distal analysis pre-Bronze Age and Bronze Age populations were included as sources. Next we run a so-called proximodistal analysis, in which just the most relevant distal sources were included in the Left population list, supplemented with a large number of post-Bronze Age populations. In latter runs potentially more relevant proximal sources competed with distant Bronze Age sources, and plausible models with distal sources indicated the lack of relevant proximal sources. In some cases, we used modern populations as sources, because the more relevant ancient sources were seemingly unavailable.

We optimized the Right populations for each run. After several initial runs with a diverse set of Right populations, we collected the models with at least one significant Z score, from the detailed “gendstat” lines of the log files of all qpAdm models. We also counted how many models we would not reject if we excluded the F4 statistics with significant Z scores of a given Right population. Based on this information we could test if all the right populations were needed to reject the models. Then we repeated the qpAdm analysis with the optimally reduced Right populations until most Right populations were needed to reject models. As an important exception, we always kept Right populations that measured the main genetic components of our test population.

Since all of our Test populations were Eurasian samples we used a suitable outgroup, (Ethiopia_4500BP_published.Sg) as Right Base throughout our analyses. In order to further exclude suboptimal models, finally we applied the “model-competition” approach described in^21^. As each model-competition run gave a different P-value with different standard deviations, we deemed it more informative to provide the maximum, minimum and average P-values for the best final models, instead of the P-values and standard deviations of the original models.

Many of our samples were part of genetic clines between East and West Eurasia. In order to reveal the genetic ancestry of individual samples within genetic clines the identified genetic groups at the eastern and western extremes were also added to the Left-populations as sources. Many of the samples within clines could be modelled as simple two-way admixture of these two populations, or three way admixtures with a third source. The remaining individuals were considered genetic outliers, which were modelled from different sources.

### f3-statistics

Outgroup f3-statistics is suitable to measure shared drift between two test populations after their divergence from an outgroup^59^ thus providing a similarity measure between populations. We measured the shared drift between the identified homogeneous new genetic groups in our sample set and all published modern and ancient populations, to identify populations with shared evolutionary past. As an outgroup we used African Mbuti genomes and applied ADMIXTOOLS^58^ to calculate f3 statistics. Admixture f3-statistics in the form f3(Test; X, Y) can be used to identify potential admixture sources of the Test population^58^, and most negative f3 values indicate the major admixture sources. We used the qp3Pop program of ADMIXTOOLS with the *inbreed: YES* parameter.

### Two dimensional f4-statistics

To measure different levels of bi-directional gene flow into populations with shared genomic history Narasimhan et al. 2019^21^ applied a so called “Pre-Copper Age affinity f4-statistics”, with a 2-dimensional representation of the f4 values from two related statistics. This way populations or individuals with significantly different proportions of ancestry related to the two sources can be visualized. We applied this 2-dimensional f4 statistics to measure gene flow from multiple sources into members of the same population. This way we could explore the fine sub-structure of populations and identify potential sources of the gene flow.To calculate f4-statistics we used the qpF4ratio from ADMIXTOOLS^58^.

### Dating admixture time with DATES

The DATES algorithm^21^ was developed to infer the date of admixture, and this software was optimized to work with ancient DNA and single genomes. As qpAdm often revealed that a two- or three-way admixture well explains the genome history of the studied population, we used DATES to determine admixture time.

## Data availability

The aligned sequences are available through the European Nucleotide Archive (http://www.ebi.ac.uk/ena) under accession number PRJEB49971. The previously published data co-analysed with our newly reported data can be obtained as described in the original publications, referenced in Supplementary Table 3.

## Acknowledgements

We are grateful to our archaeologist colleagues Gabriella M. Lezsák and Andrej Novicsihin for providing us the Anapa samples, and Gábor Lőrinczy for his help regarding the Avar material. We are thankful to all the museum curators and archaeologists who provided bone material for this study; Herman Ottó Museum Miskolc, Laczkó Dezső Museum Veszprém, Budapest History Museum, Ferenczy Museum Szentendre, Dobó István Castle Museum Eger, Jósa András Museum Nyíregyháza, Katona József Museum Kecskemét, Janus Pannonius Museum Pécs.

## Funding

This research was funded by grants from the National Research, Development and Innovation Office (K-124350 to T.T. and TUDFO/5157-1/2019-ITM; TKP2020-NKA-23 to E.N.), The House of Árpád Programme (2018–2023) Scientific Subproject: V.1. Anthropological-Genetic portrayal of Hungarians in the Árpadian Age to T.T. and No. VI/1878/2020. certificate number grants to E.N.

## Conflicts of Interest

P.L.N. from Praxis Genomics LLC, I.N. and D.L. from SeqOmics Biotechnology Ltd. and Zs.G. from Ásatárs Ltd. were not directly involved in the design and execution of the experiments or in the writing of the manuscript. This affiliation does not alter our adherence to Nature’ policies on sharing data and materials.

## Additional Information

### Extended Data Figures

**Extended Fig. 1:**
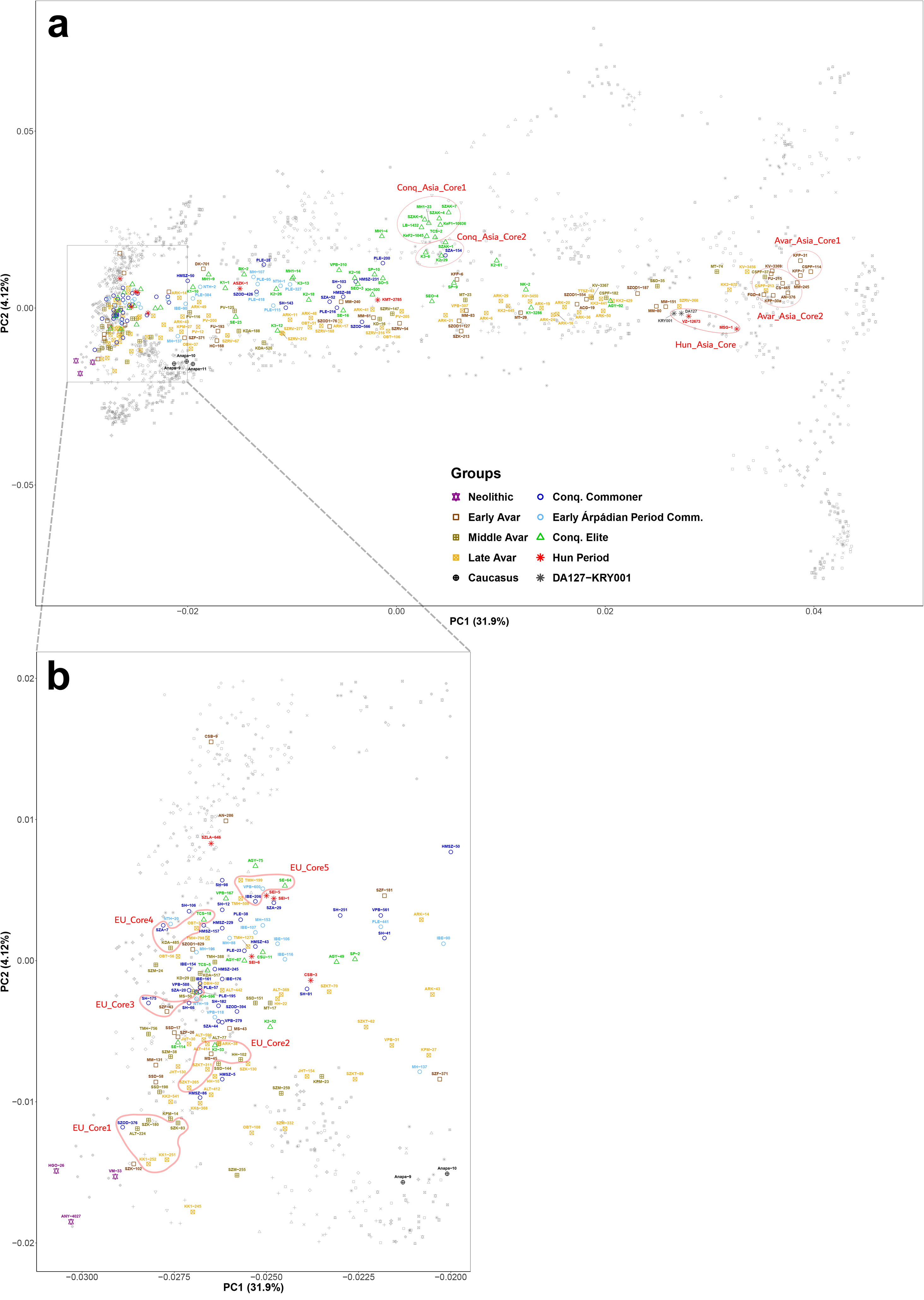
PCA of ancient samples. **a,** PCA of the Conqeror and Avar clines with sample names. **b,** Enlarged PCA of the EU-cline with sample names.

**Extended Fig. 2:**
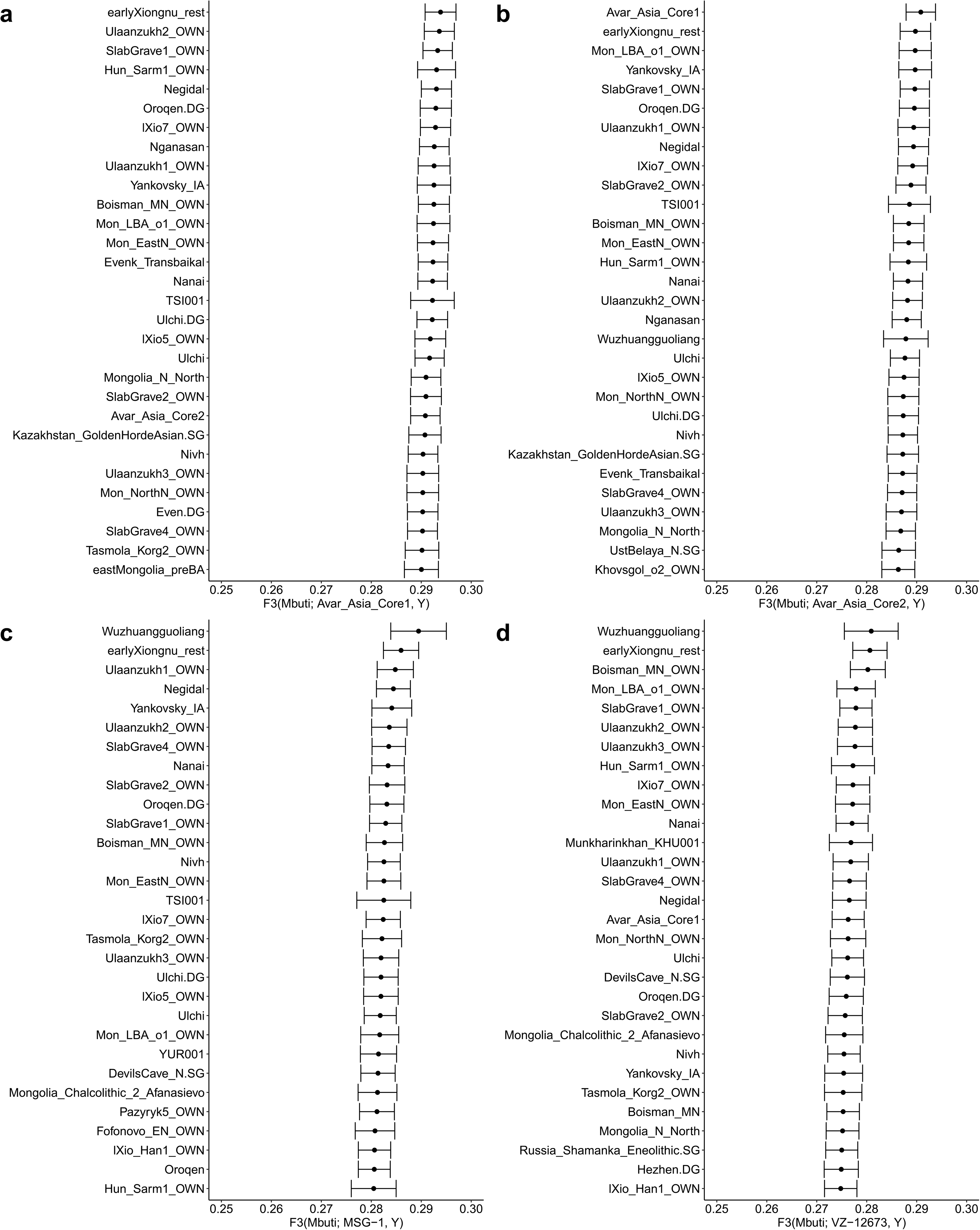
Outgroup f3-statistics for Avar_Asia_Core and Hun_Asia_Core individuals. **a,** Outgroup f3-statistics for Avar_Asia_Core1, **b,** Avar_Asia_Core2, **c,** MSG−1 and **d,** VZ–12673 Hun_Asia_Core individuals, in the form of F3(Mbuti; Test, Y). Populations with the top 30 f3 values are shown.

**Extended Fig. 3:**
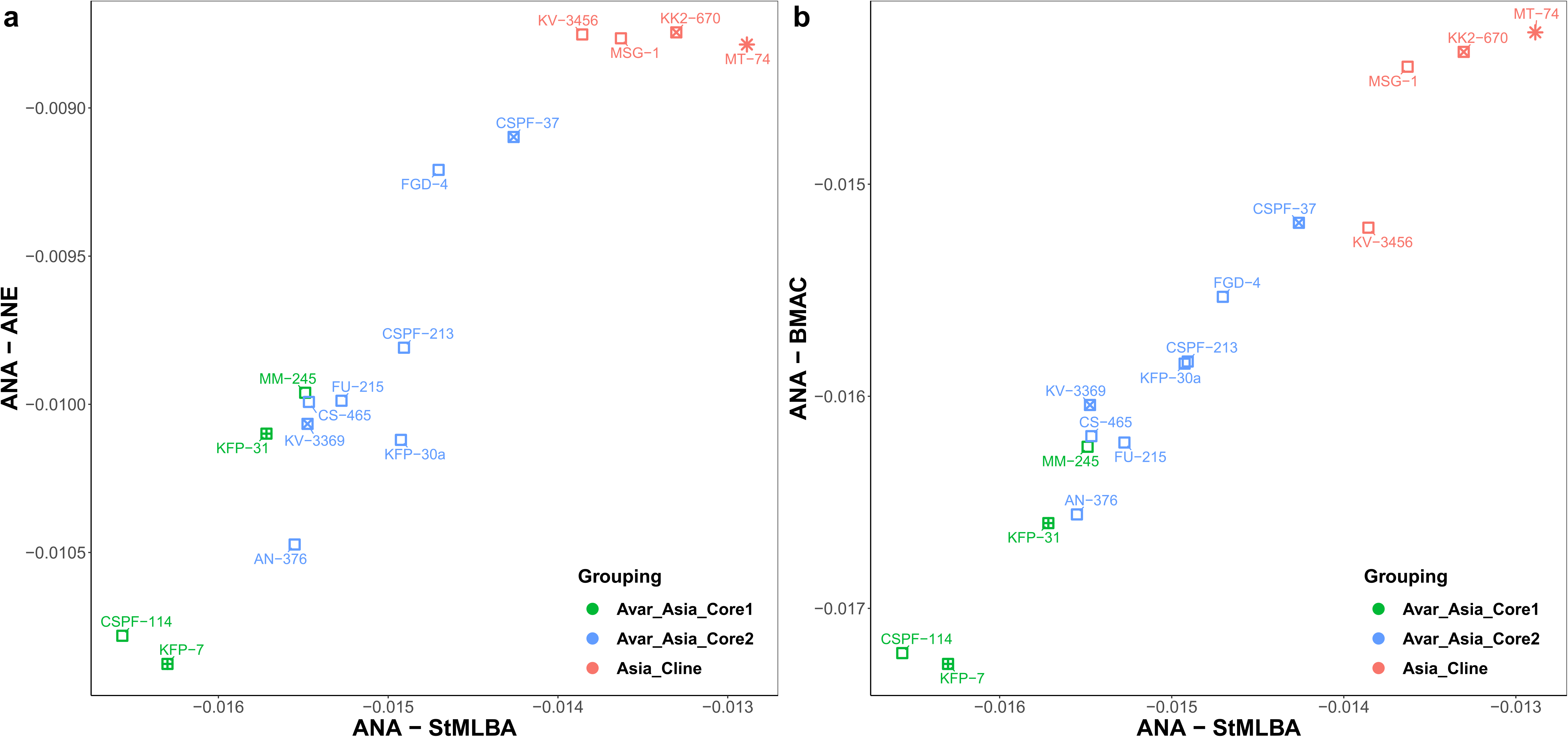
2-dimensional f4-statistics for Avar_Asia_Core individuals. **a,** 2-dimensional f4-statistics in the form of *f4(Ethiopia_4500BP.SG, Test; Ulaanzuukh_SlabGrave, Russia_MLBA_Sintashta)* versus *f4(Ethiopia_4500BP.SG, Test; Ulaanzuukh_SlabGrave, Uzbekistan_BA_Bustan)* measuring the proportion of Steppe_MLBA versus BMAC ancestries in comparison to ANA, and **b,***f4(Ethiopia_4500BP.SG, Test; Ulaanzuukh_SlabGrave, Russia_MLBA_Sintashta)* versus *f4(Ethiopia_4500BP.SG, Test; Ulaanzuukh_SlabGrave, Kazakhstan_Eneolithic_Botai.SG*) measuring the proportion of Steppe_MLBA versus ANE ancestries in comparison to ANA.

**Extended Fig. 4:**
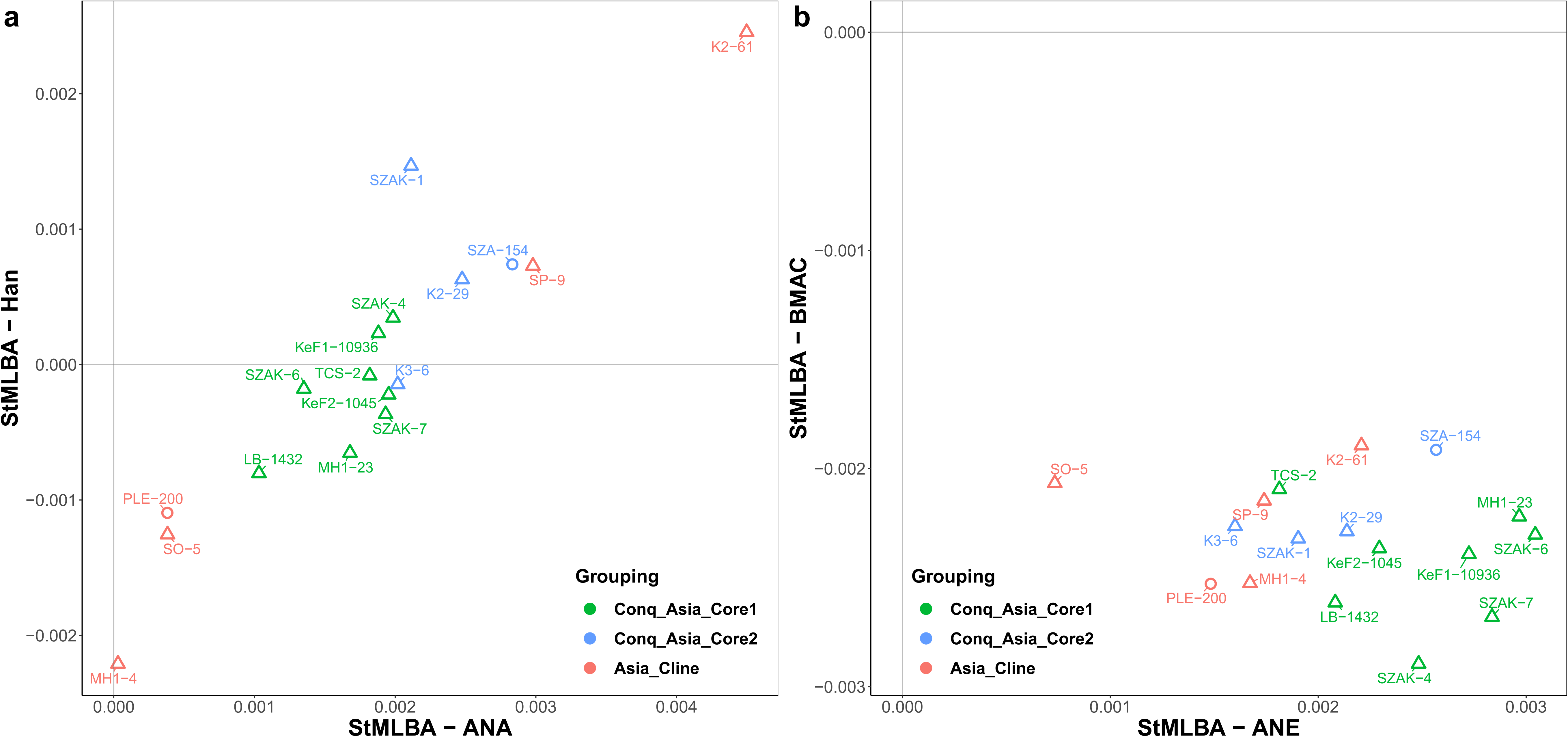
2-dimensional f4-statistics for Conq_Asia_Core individuals. **a,** 2-dimensional f4-statistics in the form of *f4(Ethiopia_4500BP, Test; Sintashta, Ulaanzuukh_SlabGrave)* versus *f4(Ethiopia_4500BP, Test; Sintashta, Miao_modern)* measuring the proportion of ANA and Han ancestries in comparison to Steppe_MLBA, and **b,***f4(Ethiopia_4500BP, Test; Sintashta, Kazakhstan_Eneolithic_Botai.SG)* versus *f4(Ethiopia_4500BP, Test; Sintashta, Uzbekistan_BA_Bustan)* measuring the proportion of ANE and BMAC ancestries in comparison to Steppe_MLBA.

**Extended Fig. 5:**
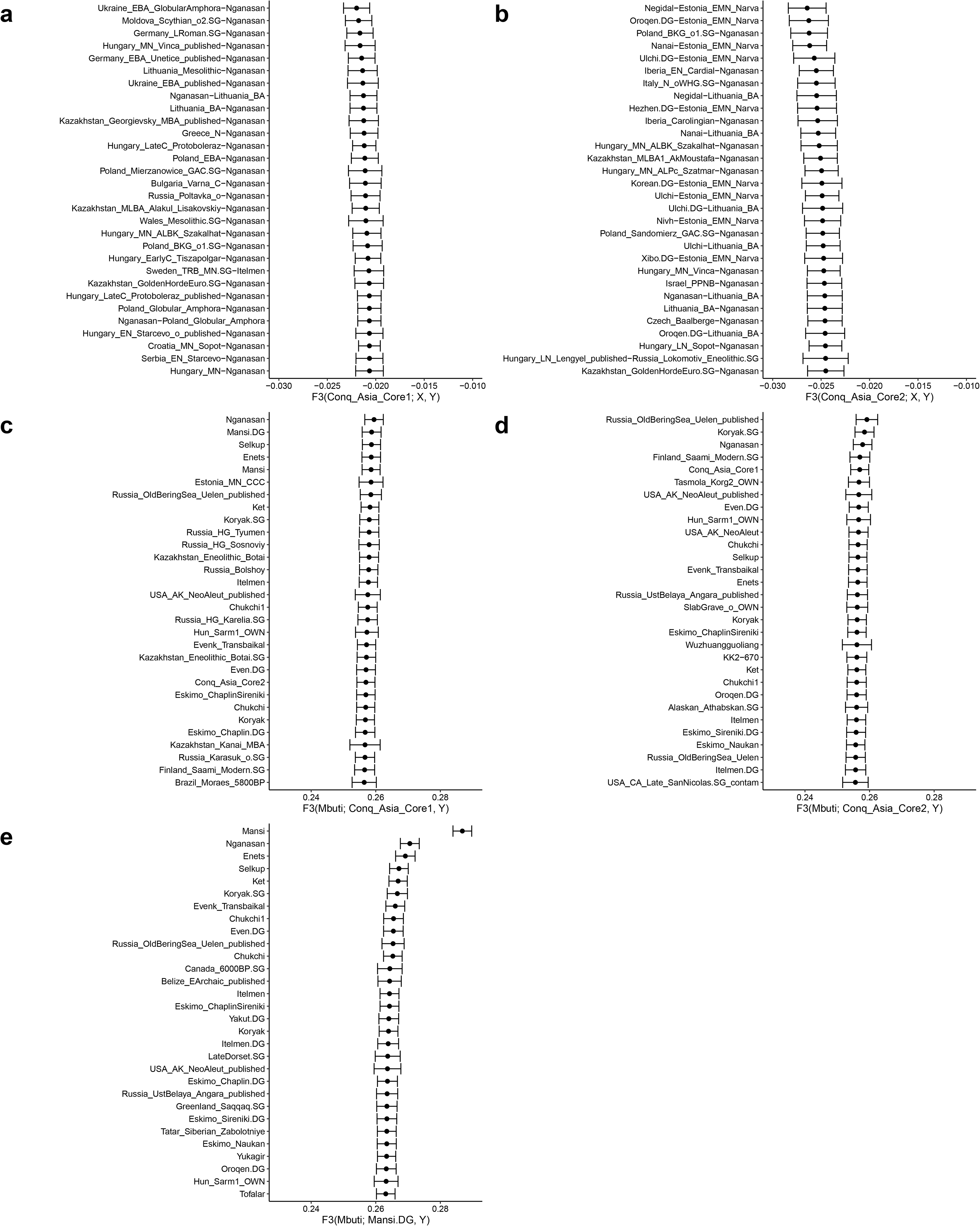
Admixture and outgroup f3-statistics for Conq_Asia_Core and modern Mansis. **a,** The top 30 population pairs giving most negative f3 values in the statistics of f3(Conq_Asia_Core1; X, Y). **b,** The top 30 population pairs giving most negative f3 values in the statistics of f3(Conq_Asia_Core2; X, Y). **c,** Outgroup f3-statistics for Conq_Asia_Core1, populations with top 30 f3 values are shown from the statistics f3(Mbuti; Conq_Asia_Core1, Y). **d,** Outgroup f3-statistics for Conq_Asia_Core2, populations with top 30 f3 values are shown from the statistics f3(Mbuti; Conq_Asia_Core2, Y). **e,** Outgroup f3-statistics for modern Mansis, top 30 f3 values are shown from the statistics f3(Mbuti; Mansi.DG, Y).

**Extended Fig. 6:**
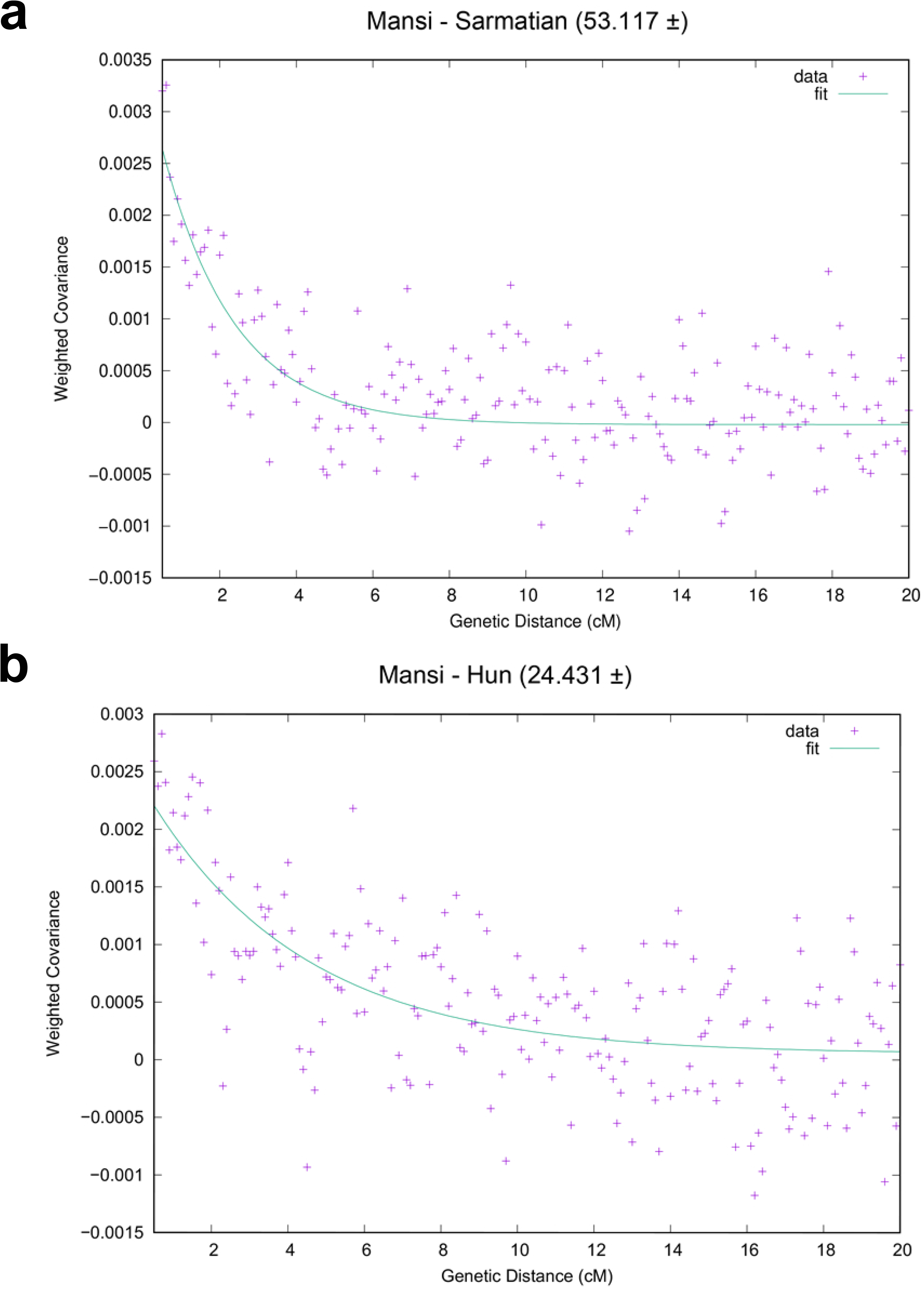
DATES analysis. **a,** DATES analysis for estimating Mansi-Sarmatian and **b,**Mansi-Hun admixture time in the Conq_Asia_Core target. Figures show the weighted ancestry covariant decays for the indicated 2-way admixture sources. Curves show the fitted exponential functions, from which the number of generations since admixture are calculated by the program.

## Supplementary Information

This files contains Archaeological background, Supplementary Methods (Sampling strategy, Population genetic analysis strategy and qpAdm analysis strategy) and Supplementary Results (detailed description of each analysis).

**Supplementary Table 1: Summary of Sequencing and qpAdm modeling results**

**Table S1a:** Summary of sequencing results for each genome, including read numbers, coverage details, estimated contamination, genetic sex determination results, and the determined mitogenome and Y-chromosomal haplogroups. Here we also provide the graveyard, archaeological culture and anthropological information for each sample.

**Table S1b:** Summary of the qpAdm models for each sample extracted from Supplementary Tables 5-7, compared to their position on the PCA map, grouped according to model types.

**Supplementary Table 2:** Radiocarbon data

**Table S2a:** Radiocarbon data for 50 samples from this study, first published here, and

**Table S2b:** Radiocarbon data for for 23 samples, which had been published in other studies.

**Supplementary Table 3:** Hierarchical Ward clustering

**Table S3a:** Hierarchical Ward clustering of 2635 ancient genomes (including our 271 genomes) according to the pairwise weighed Euclidean distances of their first 50 PCA Eigenvalues and Eigenvectors. Here we list each ancient genome used in the present study, together with their ID-s and references. We also indicate published genomes, which were regrouped in our analysis, together with the short forms, which appear in our qpAdm tables (Supplementary Tables 5-7).

**Table S3b:** Reviewable list of the genomes, which were regrouped in our analysis, together with the short forms, extracted from Table S3a

**Supplementary Table 4: Unsupervised ADMIXTURE results** (K=7)

**Table S4a:** Unsupervised ADMIXTURE results for 3277 genomes including 1010 modern and 2027 ancient published genomes, plus 240 ones from present study. Samples are arranged according to genome similarity of individuals.

**Table S4b:** The same results as in Table S4a, but samples are grouped into populations and arranged according to genome similarity of populations.

**Supplementary Table 5: Distal and proximodistal qpAdm results for Hun period individuals and Anapa samples from the Caucasus.**

**Table S5a:** qpAdm distal 2-source modeling of MSG-1 and VZ-12673 Hun_Asia_Core samples.

**Table S5b:** qpAdm proximodistal 2-source modeling of MSG-1 and VZ-12673 Hun_Asia_Core samples.

**Table S5c:** qpAdm proximal 2-source modeling of KMT-2785 and ASZK-1 Hun-cline samples.

**Table S5d:** qpAdm proximal 2-source modeling of EU-cline Hun period individuals.

**Table S5e:** qpAdm proximal 2-source modeling of SZLA-646.

**Table S5f:** qpAdm proximodistal 2-source modeling of Anapa from Iranian sources.

**Table S5g:** qpAdm proximodistal 2-source modeling of Anapa from Steppe sources.

**Supplementary Table 6: Distal and proximodistal qpAdm results for Avar_Asia_Core and each member of the Avar-cline**.

**Table S6a:** qpAdm distal 2-source modeling of Avar_Asia_Core1 and 2.

**Table S6b:** qpAdm proximodistal 2-source modeling of Avar_Asia_Core1 and 2.

**Table S6c:** qpAdm 2-source modeling of Avar_cline individuals.

**Table S6d:** qpAdm 3-source modeling of Avar_cline individuals.

**Table S6e:** qpAdm modeling of within cemetery clines from the Avar period.

**Table S6f:** qpAdm 2-source modeling of Avar period individuals in the EU-cline.

**Supplementary Table 7: Distal and proximodistal qpAdm results for Conq_Asia_Core, Mansis, and each member of the Conqueror-cline**.

**Table S7a:** Distal qpAdm 3-source modeling of Conq_Asia_Core1, Conq_Asia_Core2 and Mansi.

**Table S7b:** qpAdm distal 4-source modeling of Conq_Asia_Core1.

**Table S7c:** qpAdm 3-source proximodistal modeling of Conq_Asia_Core1 and 2.

**Table S7d:** qpAdm 2-source modeling of Conq-cline individuals.

**Table S7e:** qpAdm 3-source modeling of Conq-cline individuals

**Table S7f:** qpAdm 2-source modeling of Conquest period EU-cline individuals.

**Supplementary Table 8: List of modern Eurasian individuals used as PCA background** in Fig. 2, together with their ID-s and references.

**Supplementary Table 9: List of genetic relatives identified in this study** using the 1240K data set and the PCAangsd software. We indicate the kinship level, and also provide mtDNA and Y-chromosome data for each individual. We also indicate the individuals, which were left in the population genetic analysis.

## Contributions

T.T and E.N. initiated and led the study. P.L.N., I.N., D.L., A.G. and R.G., produced sequence data. Z.M., T.K. and E.Ny., curated sequence data. Z.M., O.S. and T.T., analysed the data. T.T., Z.M. and O.S., interpreted results with considerable input from I.R. Zs.G., C.H., S.V., L.K., Cs.B., Cs.Sz., G.Sz., E.G., A.P.K., Zs.R., B.G., B.Ny.K., Sz.S.G., P.T. and B.T. excavated skeletons and provided archaeological data. K.M., G.I.B.V., B.K., E.N. and O.Sz. sampled skeletons and made the DNA work. B.T., A.M., Gy.P. and Zs.B., described/analysed skeletons and provided bone material. A.Z. and F.M. provided additional DNA material. T.T. wrote the manuscript with contributions from all authors. All authors contributed to final interpretation of data.

## Notes

### Competing Interest Statement

P.L.N. from Praxis Genomics LLC, I.N. and D.L. from SeqOmics Biotechnology Ltd. and Zs.G. from Asatars Ltd. were not directly involved in the design and execution of the experiments or in the writing of the manuscript. This affiliation does not alter our adherence to journals policies on sharing data and materials.

