## Supplementary Information for "Whole genome analysis sheds light on the genetic origin of Huns, Avars and conquering Hungarians"

#### Table of Content

**S1. Arcahaeological Background**________________________________________________4
S1.1 General archaeological description of the Hun period in the Carpathian Basin__________________________________________________4
S1.2 Description of the studied Hun Period cemeteries and graves___________________5
S1.2.1 Árpás-Dombiföld (ASZK)______________________________________5
S1.2.2 Budapest-Vezér utca (VZ)_____________________________________6
S1.2.3 Csongrád-Berzsenyi utca (CSB)_________________________________6
S1.2.4 Kecskemét-Mindszenti-dűlő (KMT)______________________________7
S1.2.5 Marosszentgyörgy-Kerekdomb (MSG)____________________________7
S1.2.6 Sándorfalva-Eperjes, Ivótavak (SEI)_____________________________8
S1.2.7 Szilvásvárad-Lovaspálya (SZLA)________________________________9
S1.3 General archaeological description of the Avar period in the Carpathian Basin__________________________________________________9
S1.4 Description of the studied Avar Period cemeteries and graves________________11
S1.4.1 Ároktő-Csík Gát (ACG)_______________________________________11
S1.4.2 Alattyán-Tulát (ALT) ________________________________________11
S1.4.3 Apátfalva-Nagyútdűlő (AN)____________________________________12
S1.4.4 Hortobágy-Árkus-Homokbánya (ARK)___________________________13
S1.4.5 Budapest–Csepel-Kavicsbánya (CS) _____________________________15
S1.4.6 Csólyospálos-Felsőpálos (CSPF)_________________________________15
S1.4.7 Dunavecse-Kovacsos dűlő (DK)_________________________________16
S1.4.8 Fajsz-Garadomb (FGD)_______________________________________16
S1.4.9 Felgyő-Ürmös tanya (FU)_____________________________________17
S1.4.10 Hajós-Cifrahegy (HC)_______________________________________17
S1.4.11 Homokmégy-Halom (HH)_____________________________________17
S1.4.12 Jánoshida-Tótkérpuszta (JHT)________________________________18
S1.4.13 Kaba-Dögös (KD)__________________________________________19
S1.4.14 Kiskundorozsma-Daruhalom-dűlő (KDA)_________________________19
S1.4.15 Kiskundorozsma-Kettőshatár I (KK1)____________________________20
S1.4.16 Kiskundorozsma-Kettőshatár II (KK2)___________________________21
S1.4.17 Kunpeszér-Felsőpeszéri út, Homokbánya (KFP)____________________21
S1.4.18 Kiskőrös-Pohibuj Mackó dűlő (KPM)____________________________22
S1.4.19 Kiskőrös-Vágóhídi dűlő (KV)__________________________________23

S1.4.20 Makó-Mikócsa-halom (MM)____23
S1.4.21 Mélykút-Sáncdűlő (MS)____24
S1.4.22 Madaras-Téglavető (MT)____25
S1.4.23 Orosháza-Béke tsz., Homokbánya (OBH)____26
S1.4.24 Orosháza-Bónum Téglagyár (OBT)____26
S1.4.25 Pitvaros-Víztározó (PV)____27
S1.4.26 Sükösd-Ságod (SSD)____28
S1.4.27 Szeged-Fehértó-A (SZF)____29
S1.4.28 Szeged-Kundomb (SZK)____30
S1.4.29 Székkutas-Kápolnadűlő (SZKT)____31
S1.4.30 Szeged-Makkoserdő (SZM)____32
S1.4.31 Szegvár-Oromdűlő (SZOD1)____33
S1.4.32 Szarvas-Grexa téglagyár (SZRV)____33
S1.4.33 Tiszafüred-Majoros-halom (TMH)____34
S1.4.34 Tatárszentgyörgy-Szabadrétpuszta (TTSZ)____35
S1.5 General archaeological description of the Hungarian Conquest Period____36
S1.6 Descriptions of the studied 10-11th-century cemeteries and graves____37
S1.6.1 Algyő-258. kútkörzet (AGY)____37
S1.6.2 Bugyi-Kisványpuszta (BK)____38
S1.6.3 Szeged-Csongrádi út (CSU)____38
S1.6.4 Homokmégy-Székes (HMSZ)____39
S1.6.5 Ibrány-Esbóhalom (IBE)____40
S1.6.6 Karos-Eperjesszög I, II, III (K1, K2, K3)____41
S1.6.7 Kenézlő-Fazekaszug I (KeF1)____44
S1.6.8 Kenézlő-Fazekaszug II (KeF2)____44
S1.6.9 Szeged-Kiskundorozsma-Hosszúhát (KH)____45
S1.6.10 Ladánybene-Benepuszta (LB)____45
S1.6.11 Magyarhomorog-Kónyadomb (MH)____46
S1.6.12 Nagykőrös-Fekete dűlő (NK)____47
S1.6.13 Nagytarcsa-Homokbánya (NTH)____48
S1.6.14 Püspökladány-Eperjesvölgy (PLE)____48
S1.6.15 Szeged-Öthalom (SEO)____50
S1.6.16 Sándorfalva-Eperjes (SE)____50
S1.6.17 Sárrétudvari-Hízóföld (SH)____51
S1.6.18 Sárrétudvari-Őrhalom (SO)____53
S1.6.19 Sárrétudvari-Poroshalom (SP)____53
S1.6.20 Szakony-Kavicsbánya (SZAK)____54
S1.6.21 Szegvár-Szőlőkalja (SZA)____55
S1.6.22 Szegvár-Oromdűlő (SZOD)____56
S1.6.23 Tiszanána-Cseh-tanya (TCS)____57
S1.7 Cemeteries with multiple periods____57
S1.7.1 Vörs-Papkert B (VPB)____57
S1.8 Neolithic cemeteries____59
S1.8.1 Alsónyék-Bátaszék, Mérnökségi telep (ANY)____59
S1.8.2 Hódmezővásárhely-Gorzsa (HGO)____59
S1.8.3 Vésztő-Mágori-halom (VM)____60
S1.9 Samples from the Caucasus region____60
S1.9.1 Anapa–Andreyevskaya Shhel (Anapa)____60
S1.8 Archaeological references____61

**S2 Supplementary Methods**______70
S2.1 Sampling strategy______70
S2.2 Population genetic analysis strategy and methods______70
S2.3 qpAdm analysis strategy______71
S2.3.1 Optimizing the Right populations______72
S2.3.2 Excluding suboptimal models with model-competition______72
S2.3.3 Modeling members of genetic clines______73
**S3 Supplementary Results**______73
S3.1 Kinship relations______73
S3.2 Defining EU_Core groups______74
S3.3 The Conqueror cline______75
S3.4 Defining Conqueror_Asia_Core______76
S3.4.1 Hierarchical clustering______76
S3.4.2 PCA and unsupervised ADMIXTURE ______76
S3.4.3 f3-statistics______76
S3.4.4 Two-dimensional f4-statistics______77
S3.4.5 distal qpAdm ______77
S3.4.6 proximodistal qpAdm______79
S3.4.7 Dating admixture time with DATES ______80
S3.5 qpAdm modeling of Conqueror cline members______82
S3.6 The Avar cline ______83
S3.7 Defining Avar_Asia_Core______83
S3.7.1 Hierarchical clustering and ADMIXTURE ______83
S3.7.2 f3-statistics______84
S3.7.3 Two-dimensional f4-statistics ______84
S3.7.4 distal qpAdm ______84
S3.7.5 proximodistal qpAdm ______85
S3.8 qpAdm modeling of Avar cline members ______86
S3.9 Hun period samples______87
S3.9.1 PCA and unsupervised ADMIXTURE ______87
S3.9.2 f3-statistics______87
S3.9.3 distal qpAdm modeling of Hun_Asia_Core______88
S3.9.4 proximodistal qpAdm modeling of Hun_Asia_Core______88
S3.9.5 qpAdm modeling of Hun cline members______90
S3.9.6 qpAdm modeling Hun period EU cline members______91
S3.10 samples from the Caucasus______93
S3.10.1 PCA and ADMIXTURE______93
S3.10.2 qpAdm analysis ______93
S3.11 Neolithic samples ______94
S3.12 Y-chromosome and mtDNA results ______95
**S4 Genetic References**______96

### S1. Archaeological Background

#### S 1.1 General archaeological description of the Hun period in the Carpathian Basin:

The Huns, who appeared on the eastern borders of Europe in the 4th century CE, fundamentally changed the lives of barbarian groups living outside the Roman Empire, altering and dismantling the socio-economical structures that had been built up over centuries. Archaeological research has divided the period following the emergence of the Hun Empire in Europe and its fall into three major periods and a transitional period: D1 (last third of the 4th century CE - the first decade of the 5th century CE); D2 (the second third of the 5th century CE), D2/D3 (the classical Hun period, the Attila period), D3 (the half-century which following the fall of the empire/ the second half of the 5th century CE, the period after the disintegration of the Hunnic empire). In many cases, the boundaries between the epochs are difficult to draw, as some processes and archaeological sites may remain in use across two periods [1].

The "panic" caused by the sudden surge of the Huns at the end of the 4th century CE led to the migration of large and small groups throughout Central and Eastern Europe. In the Carpathian Basin these processes are very well observed, as the mixing of several former imperial cultures, which also existed during the Imperial era, can be visible within a single site. This phenomenon can probably be explained by the migration of small groups. In the eastern part of the Carpathian Basin from this period, the material of a single archaeological site often shows the influence of Chernyakhov, Przeworsk and local Sarmatian cultures. This phenomenon is an excellent illustration of the arrival of several small communities in the area. The influence of the late phase of the Eastern European Chernyakhov culture is particularly marked. The end of the period probably coincides with the mass migration of barbarian groups to the west and the final closure of the latest dateable sites of the Chernyakhov culture (the Post-Chernyakhov period) in the first decade of the 5th century CE [2,3].

After the first decades of the 5th century CE, the population in the eastern part of the Carpathian Basin may have decreased significantly, as is shown by the changes in the settlement network. The local population and the population arriving in the late 4th and early 5th centuries CE increasingly buried their dead in solitary burials, in small pairs of cemeteries, or cemeteries with a few graves (5–30). The unifying material culture in the Danube region is characterized by the predominance of elements of eastern Germanic culture alongside surviving local populations and eastern nomadic representation. Naturally, both in the Transdanubian region and in the eastern part of the Carpathian Basin, we might expect a higher proportion of pre-Hun populations to persist [4]. However, their material cultivation was also significantly influenced by the previous styles.

In the second phase of the Early Migration Period, the main features of the monumental amount of material in the area (Unternsiebenbrunn horizon) are the object types (plate brooches, buckles) that developed in the final phase of the Chernyakhov culture, local late Roman influences (pottery, chip carving belt/carved buckles) and new impulses from the East. In addition to the changing material culture, however, the study of the archaeological sites has shown that many imperial traditions survived unchanged into the 5th century CE. In the eastern part of the Carpathian Basin, the highest number of graves (70-140 graves) has been found in the Csongrád (Csongrád-Csanád County) and Ártánd (Hajdú-Bihar County) areas. These sites show both the cultivation of the earlier Sarmatian period and the new cultural influences of the Hun period [5–7].

The question of when the leadership and elite of the Hun Empire moved their headquarters to the area is still undecided. The first half of the 5th century CE and the 2nd and 3rd decades of the 5th century CE are possible, based on written sources, but no archaeological evidence can be found about this migration.

In the second third of the 5th century CE, more marked changes in the archeological material from the region can be discovered again. This can be attributed mainly to the emergence of a unified cultural identity of the Hun Empire, which also appeared in the material culture. After the shift of the centre of power, the features of the unifying material culture, which had existed together in this form for only a very short time, gradually emerged. The cultural image of the newly emerging elite, which was not linked to ethnic groups but to the nomadic state itself, was formed by the use of objects in the polychrome style (horse equipment, costume elements, weapons and their ornaments), elements of oriental-style weaponry and the wearing of large silver plate brooches. Their burials may have embodied an emerging imperial identity, since we know of very similar material from regions far from the Carpathian Basin such as Juszkowo, Jakuszowicze, Altlußheim [8–11]. For this very reason, the ethnopolitical groups that appear in the written sources of the period cannot be identified in the material culture. To fit in, the various Germanic and Sarmatian groups presumably also began to adopt to the customs of the imperial elite. It was precisely through the concentration of power along the Danube that the steppe state, governed by a small nomadic leadership, achieved its success. It succeeded in bringing together different warrior groups, mainly Germanic, and exploiting their rivalry managed to create a serious military challenge for the two Roman imperial powers [12–14]. The continuous campaigns of the nomadic polity providing steady income could have attracted small, mobile armed communities from more distant regions. The unification of material culture and the fashions set by the Hun elite are reflected in the contemporary account of Priscus of Panium. According to the source's report, a person who had risen from being a former Eastern Roman prisoner of war to the barbarian society of the free was identified by the historian as a Hun based on his clothes [15].

Local populations and newcomers often formed new communities and burial sites within a very short time. The local population quickly adapted to the culture of the immigrant groups, retaining some of their customs and funerary rites. This process, which was previously only evident in the archaeological record, have recently been confirmed by modern scientific methods (e.g., concerning the site Mőzs-Icsei dűlő) [16].

In the decades following the fall of the Hun Empire, a sharp discontinuity is not always discernible in the Carpathian Basin archaeological sites. In the first decades, certain elements of representation (silver plate brooches for women, weapons) and certain customs show continuity with the period of nomadic rule [17,18]. The reason for this, in addition to the continuity of the population, is that the material of these burials can be largely attributed to the local elite who gained power in the Hun period. In the second half of the 5th century CE, in addition to continuity, technical innovations (chip carving technique on buckles and brooches) appear in the finds. The often mixed-origin population of the 5th century CE is integrated into its own culture by the emergence of the Germanic kingdoms at the end of the century, which can be seen as the eastern branch of the western Merovingian culture (row-grave cemeteries culture) [19].

#### S1.2 Description of the studied Hun Period cemeteries and graves

##### S1.2.1 Árpás-Dombiföld, Szérűskert-1 (ASZK; Győr-Moson-Sopron County) [20]

**ASZK-1**

The solitary grave of a young adult male was discovered near the present-day Árpás village. The body was laid on its back, in a shallow grave with North-South orientation, among the ruins of the Roman Mursella municipium. The burial was found inside a former building, next to the wall. The building was presumably no longer occupied at the time the grave was dug as rubble and building stones were found in the fill of the burial (the complete destruction of the building may have taken place later). Grave goods included a golden belt buckle, golden trouser and boot-buckles, sabretache with iron knife and forceps, needle-like bronze stick, and a bronze fragment. Further findings were a jar, a glass cup, a large bronze vessel (cauldron or kettle), animal bones (cattle leg bone and sheep sacrum), and a golden coating of a probably wooden animal sculpture. The jar and glass cup have Roman parallels, but the funerary custom, costume accessories and animal sculpture have Hun Age eastern steppe analogies. The finding was dated to the 2nd third and middle of the 5th century CE (D2 or D2/D3 period) when the Huns ruled Pannonia.

##### S1.2.2 Budapest-Vezér utca (VZ; Pest County) [21]

**VZ-12673:**

In December 1961, construction workers found metal objects and horse bones during a construction project in Vezér Street in Budapest. Workers excavated the foot end of the former burial, as well as the horse skull and vertebrae. The complete excavation of the human skeleton and the grave was carried out by Tibor Nagy following the discovery. The site was located on the banks of the Rákos stream and possibly dug at a prominent point. The finds and the burial were forgotten until the 5th-century grave was processed by Margit Nagy.

The burial was oriented in the NE–SW directions. The grave pit was 170 cm long and 90 cm wide, and the human remains were found at a depth of 112 cm. The grave contained the remains of a young adult male with Europid and Mongoloid characteristics on the skull, together with a partial horse burial (only the skull and vertebrae remained), presumably placed at the feet, in front of the body of the deceased. The workers could only have disturbed the legs and feet before the excavation. The skull was slightly depressed, the hands resting on the pelvis. The following objects were recovered from beside the skeleton: next to the human skeleton were the B-shaped iron buckle (on the sacrum), seven gold mounts with garnet incrustations (between the thighs), and a small, oval iron buckle (on the inner side of the left thigh). Unfortunately, some of the objects that were included in the documentation are no longer to be found (a vessel fragment behind the skull; a 12 cm–long, single-edged knife; and a disc-shaped 'button' next to the lower ribs on the right side of the skeleton).

The construction workers found several other artefacts in this part of the grave and in the soil they had removed from the pit: an iron bell, a bronze bell, a horse bit, the fragment of an iron bar covered with gold foil, and a diamond-shaped harness ornament.

The grave and its artefacts are among the few archaeological materials of the 5th century CE that can be traced back to the nomadic traditions of Eastern Europe, of which only a few good parallels are known from the Carpathian Basin. Apart from the objects, the burial rites (partial horse burial above the dead, lack of weapon accessories) are closely linked to the Eastern European region, from which we know good analogies. The finds can be dated roughly to the middle third to the middle of the 5th century CE (D2 or D2/D3 period).

##### S1.2.3 Csongrád-Berzsenyi utca (CSB; Csongrád-Csanád County) [7]

A Hun pot and possibly grave finds from the site were rescued in the 1930s by László Tary, a local polyhistor doctor. In the early 1950s, the grave of the first individual with an artificially deformed skull was discovered in Berzesenyi Street during the laying of a sewer. According to residents, several small mounds were visible in the area before the main earthworks started. Based on this information, Mihály Párducz began a systematic excavation in 1959. He succeeded in excavating a total of 9 burials. The majority of the burials show the continuation of the customs of the earlier Sarmatian period since they are predominantly south-north oriented. Although in his publication Párducz had classified graves 8 and 9 as belonging to the Hun period, it is now clear based on the finds that they were burials dating to the Early Avar period. The gender distribution of the graves in the cemetery was as follows: three females, three males, two children, and one individual with an artificially deformed skull of undetermined sex. However, the cemetery is not fully excavated. The graves also yielded Csongrád-Zieling L-type shield boss, spearheads, iron brooches with a returned foot, buckle with animal heads turned towards each other, and a ribbed/ cup. The finds excavated here were dated from the turn of the 4th and 5th centuries CE, the typical D1 period. In the absence of a complete excavation, we cannot determine the upper chronological limits of the use of the site. In addition to the local Sarmatian features (orientation, pottery), several new features appear in the grave finds, and the number of weapon attachments increases compared to the earlier period. The cemetery at Csongrád-Berzsenyi Street clearly shows features characteristic of sites in the Csongrád area dating to the 4th-5th century CE. In addition to many ritual elements of the Sarmatian population, new types of objects and customs appear in the region under study, so that it is possible that the local autochthonous population, who were presumably joined by newcomers, was continued to live here in the Hun period.

**CSB-3: Grave No. 3:**
The grave of an adult male was found at a depth of 157 cm, with a grave pit 89 cm wide and 237 cm long, while the length of the human skeleton measured in the grave was 186 cm. The burial was oriented in the SSW-NNE directions. The grave has been disturbed/robbed, and only the skullcap and the right-side bones remain in their original position. Near the original location of the pelvis a round iron buckle, fragments of an iron returned foot brooch, an iron nitt, and a flint stone were found. While next to the left leg an iron spearhead was unearthed with its edge turned upwards. The grave was dated to the end of the 4th century CE and the beginning of the 5th century CE.

**CSB-9: Grave No. 9 (Early Avar period):**
The remains of an adult female were found at a depth of 118 cm. The E-W oriented grave pit was 59-47 cm wide and 211 cm long. Both hands of the deceased rested on the pelvis. An iron buckle was found here, while earrings with a spherical pendant were found around the temporal bone. A fragment of a dished grey ceramic vessel was also recovered from the grave. The burial can be dated to the Early Avar period based on the earrings and the orientation.

##### S1.2.4 Kecskemét-Mindszenti-dűlő-RL 11/2785 (KMT; Bács-Kiskun County) [22]

**KMT-2785**
During a preventive excavation of the expansion related to the Mercedes factory on the outskirts of Kecskemét town (Bács-Kiskun County, Hungary) in 2017, near a late Sarmatian-Hunnic age settlement, a solitary niche grave was found, containing a wealthy young adult male with artificially strongly deformed Mongoloid skull. The grave was oriented in the N-S directions, and the grave goods included a short sword, crescent-shaped gold hair ring, back-bended gold plate with a ribbed surface (which was probably the decoration of the knife's handle), a kidney-shaped silver buckle with a slightly bent side covered with gold foil, another buckle that suspended the sword, two shoe buckles (both buckles were made with a similar technique), a worked bone object, a Roman bronze coin, a small lithic flake and a Murga-type jug. The buckles made of silver and plated with gold, are showing Hunnic age analogies. Based on the burial rite and the finds, the tomb can be dated to the 2$^{nd}$ quarter of the 5th century CE. The burial customs and the findings differ from the traditions of the local Sarmatian period graves.

##### S1.2.5 Marosszentgyörgy-Kerekdomb/ Singeorgiu de Mures-Kerekdomb (MSG; Mureş County, Romania) [23]

During the construction of the bypass route between Corunca-Ernei (Mureş County, Romania), in the course of preventive excavations, the Archaeological Department of the Mureş County Museum discovered a Roman village and a necropolis dating from the Migration Period (according to the funerary offerings and the funeral rites), near Sângeorgiu de Mureş. The excavations brought to light nineteen houses, fifteen pits and three single graves. According to the archaeological material, the settlement can be dated to the second half of the 3rd century CE and the beginning of the 4th century CE, and the grave goods dated the cemetery to the end of 4th century CE and the beginning of 5th century CE. The graves were discovered on the upper part of the second terrace of the Terebici River, situated in line. Two graves were located inside the village. One burial (grave no. 3 Cx. no 58) was completely robbed, and only the mold of the human skeleton in a secondary position and remains of wood construction on the bottom of the grave could be observed. We involved one sample in our analysis.

**MSG-1: Grave No. 1 (Cx. No. 41):**
Grave No. 1 was located inside the settlement, trenched in a Roman period house. It contained a partial horse burial oriented North-South, typical for the Migration Period. Unfortunately, the grave was robbed, and the right side of the human skeleton and a significant part of the horse skeleton was destroyed. The preservation of human remains was relatively good, and they belonged to a young adult male with an artificially deformed Mongoloid skull. The grave goods included a grey jug, a stemmed goblet made of dark yellowish-green glass, a one-sided antler comb with a case decorated with concentric circles and bronze plate, four silver rivets, an iron knife, and an inlaid golden mount (of a dagger gripe) decorated with cloisonné technique. Although the grave goods of the deceased and the funerary rite show strong Steppe influences, the latest date of the burial, based on the objects found in the grave, suggests that it took place in the second half and last third of the 5th century CE.

**S1.2.6 Sándorfalva-Eperjes, Ivótavak (SEI; Csongrád-Csanád County)** [24]
Five circular ditches (ditched greaves) and eleven graves were found in 1981, and the site is considered fully excavated (Vörös 1985). From the eleven graves, seven burials were surrounded by these circular ditches, which is a burial custom characteristic of the Sarmatian Period burials. Concerning the burial customs, traces of wooden coffins were also registered in the graves (e.g., metal parts, layers of organic material remains above and under the skeleton). The graves formed lines in the cemetery. Although most of the burials were robbed, the grave finds were considered rather wealthy. The grave goods are composed of weapons (spearheads, swords), belt accessories (buckles and metal plates), jewelry (e.g., a string of beads, bracelets), brooches, different types of vessels made of clay or glass, and implements (knives, bone needle case). In addition, a Roman coin from the 1st century was found, but it was not considered to have a dating value. Based on the archaeological material, the cemetery was dated to the 4th and 5th centuries CE, i.e., the Hun period. The evaluation of the finds, burial customs and ethnic background of the population is, however, much more complex since the archaeological material show strong analogies with Sarmatian period cemeteries and other series connected to Germanic populations. The distribution of the burials, the composition of the grave goods suggests that a small population with inner family relationships were buried in this cemetery. The results of the anthropological investigation led by Antónia Marcsik is not published yet, but two sub-adults, five adult males and three adult females were identified in the series. We involved three samples in our analysis.

**SEI-1: Grave No. 1:**
A robbed burial surrounded by a circular ditch. The skeletal remains of an adult male individual were found in the southern end of the grave. The grave goods were a sword, a knife, fragments of a non-identifiable iron tool, a bronze T-shaped brooch and another two-piece bronze brooch with a foot leaning to the side. Besides, accessories of belts (bronze and iron buckles, lead belt mounts), glass fragments of a bowl, and a Roman coin emitted in the 1st century were found. The grave was dated to the end of the 4th and the 5th centuries CE.

**SEI-5: Grave No. 5:**
The skeletal remains of an adult male were found in the burial surrounded by a circular ditch. Traces of a wooden coffin were registered in the grave, but the region of the chest was robbed. The grave finds were a sword, belt accessories (bronze belt buckle, lead belt mounts), a knife, and a glass bowl. The burial was dated to the end of the 4th and the 5th centuries CE.

**SEI-6 Grave No. 6:**

It was the burial of an adult female individual. Traces of coffin (layers of organic materials above and under the skeleton) were registered in the grave pit. The burial contained a bronze brooch with a foot leaning to the side, fragments of possibly another bronze brooch, a string of different types of beads (e.g., beads made of glass, amber, and carnelian) with miniatures of toiletries in the middle (a spoon and a needle with coiled end), a knife, bronze bracelets, a bone needle case with an iron needle inside, and a clay vessel. The grave was dated to the end of the 4th and the 5th centuries CE.

**Szilvásvárad-Lovaspálya (SZLA; Heves County)** [25]

**SZLA-646: Object 646:**

In 2016, the Dobó István Castle Museum in Eger, Hungary, carried out a preventive archaeological excavation in the southeastern part of Szilvásvárad, in the valley section of the Szalajka stream, for the construction of the Lipica Equestrian and Event Centre. Naturally, the area excavated has yielded objects from several periods (Neolithic, La Tene period, Roman Imperial age, Hun and Árpád periods). A Roman age Barbarian settlement dating back to the Imperial period was continuously used until the 4th and 5th centuries CE. On the edge of this site, 160 m from the settlement objects, a burial dated to the 5th century CE was found during the excavation.

The SW-NE oriented grave contained a middle adult female, buried in a supine position. The grave pit was 236 cm long and 80–65 cm wide, and the human remains were found at a depth of 59 cm. The grave pit was rectangular shaped with curved corners.

The following finds were recovered from the tomb: a large glass bead, a pair of tweezers, fragments of a strongly corroded iron object, an iron knife, a bronze cicada brooch, a D-shaped cast bronze buckle (at the ends of the bar are two animal heads facing each other), a bell-backed comb, and a small, single-handled jug.

Most of the finds recovered from the burials form an integral part of the material culture of the Carpathian Basin in the 5th century CE, but some objects suggest strong western connections. The pottery vessel can be considered as completely atypical within the local material, as confirmed by petrography studies. The jar presumably follows the workshop traditions of the so-called Mayen-Kolbenz region. The best analogues of the buckle are also known mainly from the Roman Empire's western provinces (Gaul). These suggest that the lady buried at Silvásvárad and her relatives may have obtained the above-mentioned objects through personal contacts, which also illustrates the mobility and interpersonal contacts of people and groups in the 5th century CE, several of our written sources recall that the Huns placed many mercenaries at the disposal of the Western Roman Empire for its battles against the barbarians (Western Goths, Burgunds) and provincial rebels (Bagaudae) in the western provinces. Based on the finds, the burial was dated in the middle third of the 5th century CE.

#### S1.3 General archaeological description of the Avar period in the Carpathian Basin:

According to Byzantine sources, the Avars arrived in Europe in the 550s CE. After establishing themselves in the Carpathian Basin in 567–568 CE, the Avar Khaganate survived for some two and a half centuries. The first phase of the Avar period can be marked with the wars against Byzantium, which seems to have ended with the aborted Sasanian-Avar siege of Constantinople (626 CE). Around this time, the Avars disappeared from the written sources. From the 630s CE, only a few scattered references imply that the Avars still lorded the Carpathian Basin before the Carolingian occupation around 800 CE.

The deconstruction of the nomadic power structures occurred simultaneously with the complex crisis of the Byzantine Mediterranean, leading to the disappearance of the booty and gifts that had sustained the system under the khagan’s rule. In this sense, the early Avar polity can be considered as a ‘secondary’ or ‘shadow’ empire [26,27], whose cohesion was maintained as a response to the strength of the Byzantine Empire. The decline of Byzantine power also meant the end of the Early Avar period from a structural point of view.

As reflected in the archaeological record, the transformation in the seventh century, which followed the disintegration of the early Avar khaganate, was on a scale that justifies the distinguishing of a new era, the 'Late Avar period' (ca. 650–820 CE). For the chronological phases of the Avar period see [28]. Chronologically, the time frame corresponds to the Dark Ages or the Transition period [29] of the Mediterranean, lasting from roughly the later seventh to the onset of the ninth century, a period that is generally described as 'the long eighth century [30].

The social and administrative structures of the Avar Khaganate were doubtless determined by its nomadic traditions [31]. In the burial context, the prominence of horses [32], certain types of weaponry [33], and the outstanding importance of the ornate belts indicate the dominance of the symbols of nomadic prestige. In the late seventh century, uniform material and – as far as this can be established from the funerary rituals – spiritual culture extending to virtually all aspects of life with minimal regional divergences emerged across the entire Carpathian Basin [34]. Behind the cultural phenomena, a transformation can be identified in the "deep structures" of the khaganate, in the economy, communication/traffic, the production of strategic resources (iron, probably copper and salt) and low-value handicraft products (copper alloys and bone objects). In the 'Middle Avar period', the most important element in the transformation was the transition to an extensive settlement system based on rural settlements [35]. These phenomena suggest that the khaganate had lost its nomadic character and integrated into the structures of the surrounding world.

However, the Avar elite could not compete with the efficient centralising ideology of Christianity and the centralised structures of the ecclesiastic hierarchies used purposfully by the Carolingians. The Avar elite and political formation kept disintegrating until the 830s CE when Avars were last mentioned in the written sources.

For an archaeological and historic introduction of a genetic paper, it is essential to pay attention to the questions of population processes. The population groups called 'Avar' were heterogeneous concerning their origin. The archaeological record testifies that the Avar age Carpathian Basin was inhabited relatively densely. The groups arriving from the steppe were of heterogeneous descent, although groups of central-Asian origin were identified among them. These people settled among the Gepids in the Great Plain and Transylvania [36] and the remnants of the Langobard and the Late Antique population in Transdanubia [37,38]. The blending between different peoples began quite quickly, as reflected by the cultural heterogeneity of the earliest grave groups of newly opened row-grave cemeteries in the 7th century CE, especially in the large cemeteries of Transdanubia [39]. This cultural heterogeneity remained strong until the late seventh century, also due to archaeologically identifiable (e.g., the cemeteries of Alattyán–Tulát [40] and Szarvas–Grexa-téglagyár [41]) internal migrations and the possibly continuous infiltration of Eastern European groups.

In Avar archaeology, there is a long-lasting debate on the impact of the migrant groups on the population density and the culture of the Avar khaganate. Traditional research preferred a rather static picture of large immigrant – or conquering – population groups that arrived to the macroregion in 567/568 CE (the Avars of Baian) and the 670s CE (the Onogur-Bulgars of Kuber). Today it seems much more plausible that the Carpathian Basin was a permanent destination of continuous but (mostly) small-scale immigrations/infiltrations from the second half of the 6th century to the late 7th, or probably the 8th century CE. However, it is hard to answer what kind of role the immigrants had plaid in the population history of the region.

The height of Early Avar military activity – a disastrous period for Byzantium – can be roughly correlated with the Late Antique Little Ice Age. At the end of the LALIA in the mid-seventh century began a milder period to the Carpathian Basin [42]. It must have contributed significantly to the population expansion, which can be detected in the expansion of the Avar settlement area [35] from the mid-7th century CE onwards. Therefore, the widening of the settlement territory to previously uninhabited territories, mostly hilly regions, besides other factors (e.g., changes in agricultural technology and the spread of back-yard animal breeding), must go back at least partially to a radical growth of the indigenous population. However, the archaeological record eludes the attempts to determine population size. Besides internal growth, the role of repeating immigrations during the 7th century CE seems to be very probably among its causes, although migrations can be documented by archaeological arguments only in few cases (e.g., [43]).

The 20th-century Central European research generally insisted on the theory that the ‘Avars’ – used as a general term for the entire population in the region – were exterminated during the Frankish Wars. However, as early medieval ethnicum was assigned based on ruling elite, the loss of the Avars meant only the defeat and more or less complete demolition of the privileged groups. However, east of the Tisza River, like in Hortobágy-Árkus (see in the present paper), the descendants of the Avar elites could have survived until the Hungarian conquest. Among ‘common people’ who dwelled in the dense settlement network, the loss of population due to the wars and uncertainty following the fall of the khaganate could have been dramatic, but certainly not complete.

In summary, global variables like climate, changes in the political milieu in the surrounding world and internal factors such as immigration, internal migrations and the advanced state of the shift to sedentism among the nomads in the Carpathian Basin had their impact on the population. After one and a half-century of renewed prosperity, the 9th century CE may have been characterized by a decrease in population numbers, although integrated with Slavic immigrants to an unknown extent, the population of the Avar khaganate survived until the Hungarian conquest period.

#### S1.4 Description of the studied Avar Period cemeteries and graves

##### S1.4.1 Ároktő-Csík Gát (ACG; Borsod-Abaúj-Zemplén County) [44]

Two hundred-fifty graves of a multi-period cemetery were excavated by Emese Lovász between 1996 and 2000. The burial ground was used continuously from the 5th to the 9th century CE, however, the archaeological material is not published yet. The skeletal remains of two hundred-forty-nine individuals were available for the anthropological investigation led by Ivett Kővári and László Szathmáry. Based on the archaeological documentation, they divided the burials into four groups. These are the Hun-German period (20 cases), Early Avar period (18 cases), Late Avar period (204 cases), and 9th-century graves (7 cases). Concerning the Early Avar period material, they identified seven sub-adult and eleven adult individuals. Besides, they described five females and seven males. In the Late Avar period group, 76 sub-adults and 100 adults were distinguished, and 61 females and 65 males were registered. Based on craniometric data, partial populational continuity was assumed between the Early and Late Avar period groups. We involved one sample in our analysis.

**ACG-19: Grave No. 19**

It was the burial of a middle-adult female. The grave was categorized as an Early Avar period case. Radiocarbon data published in this paper (643-680; 750-758 CE with 95% probability) confirmed that the individual was, indeed, buried in the Avar period. However, the fine dating of the assemblage, whether it belongs to the Early or Late Avar period of the cemetery, is unclear.

##### S1.4.2 Alattyán-Tulát (ALT; Jász-Nagykun-Szolnok County) [40,45,46]

A large Avar age cemetery with 712 graves was excavated between 1934 and 1938. The archaeological material is composed of various types of artefacts: weapons (e.g., swords, archery equipment, axes, armor fragments), amulets, jewelry (e.g., earring, strings of beads), belts with bronze and silver ornaments, bone needle cases, vessels, however, horse burial and horse-riding equipment was not found in the graves. Based on the grave-good types, Ilona Kovrig described three typological and three chronological groups of the cemetery. The burial ground was dated from the 7th to the 9th centuries CE. The anthropological material was analyzed by Sándor Wenger. He described 19 sub-adults and 225 adults, and he identified 108 female and 115 male individuals in the series. The taxonomical analysis was performed by Pál Lipták. The main part of the series is composed of skulls with Europid characteristics (mainly Brachycran group (40%) and Cromagnoid types (20%). Besides, Europo-Mongoloid features were described in certain cases.

**ALT-77: Grave No. 77**
The skeletal remains of an adult male were found in the grave. The burial contained a loop and a bone tube. The burial belonged to the first chronological group dated to the first half of the 7th century CE.

**ALT-224: Grave No. 224**
It was the burial of an adult male with Europid skull characteristics. The grave goods were a bronze ring, earrings with a pendant and belt ornaments. It was categorized in the second chronological group dated to the end of the 7th and 8th centuries CE.

**ALT-369: Grave No. 369**
The grave contained the skeleton of an adult male with Europid (Mediterranean and Nordic) skull characteristics. Belt ornaments, a loop, a knife and an iron tool with a bone handle were found in the burial. It belonged to the third chronological group dated to the 8th and first half of the 9th century CE.

**ALT-412: Grave No. 412**
The skeleton of an adult female with Europid characteristics on the skull was found in the burial. The grave contained an earring, beads and a spindle whorl. The burial belonged to the third chronological group dated to the 8th and first half of the 9th centuries CE.

**ALT-414: Grave No. 414**
Burial site of an adult female. The grave goods were earrings, melon seed-shaped beads, a spindle whorl, a knife, and a bracelet. The grave belonged to the third chronological group dated to the 8th and first half of the 9th century CE.

**ALT-442: Grave No. 442**
The skeletal remains of an adult female with Europid skull characteristics was found in the grave with a knife and an earring with a pendant. It was categorized in the second chronological group dated to the end of the 7th and 8th century CE.

**ALT-596: Grave No. 596**
An adult female with Europid skull characteristics was buried in the grave with a bronze ring, a melon seed-shaped bead and a spindle whorl. The burial belonged to the third chronological group dated to the 8th and first half of the 9th century CE.

**S1.4.3 Apátfalva-Nagyútdűlő (AN; Csongrád-Csanád County)** 47,48
During the preventive excavation of the M43 motorway on the outskirts of Apátfalva village, seventeen Avar-age graves were found by Tibor Paluch in 2010. There were graves in two burial groups laying within 80 meters from each other, which can be separated into two different chronological horizons based on the burial customs and the material. One of the groups consisted of 9 Early Avar-age burials. The most important characteristics of these are E–W orientation, end-niche grave and partial animal sacrifices. This burial group was dated to 630–650/660 CE. The other burial group consisted of 8 graves dated to the end of the 7th century CE. The skeletal remains of 15 individuals were fit for anthropological analysis. Based on the results, nine adults (seven males and two females) and six sub-adults were described. In addition, six skulls represented Europid characteristics (four of them with

Mongoloid features) and one skull belonged to the Mongoloid type. Concerning the metric and taxonomic data, the series showed similarities with other populations of the Early Avar period at the Southern part of the Great Hungarian Plain (e.g., Makó-Mikócsa-halom). We involved two samples in our analysis.

**AN-286: Grave No. 286 (Obnr 286/Snr 362):**
In the E–W orientation end-niche grave, the skeleton of an adult male individual slid upside down into the niche was found. Between the niche and the pit, the bones of a sheep were found. Besides, two stirrups, iron buckles near the skeleton, an unknown iron plate, and flint were found in the pit. The grave was dated to 630–650/660 CE.

**AN-376: Grave No. 376 (Obnr 376/Snr 461):**
In the E–W orientation end-niche grave, an 8-9-years-old sub-adult was found. The head pushed forward into the niche. The bones of a female sheep, an iron buckle, bronze belt mounts, and a stirrup were found in the pit. The grave was dated to 630–650/660 CE.

**S1.4.4 (Hortobágy-) Árkus-Homokbánya (ARK; Hajdú-Bihar County)** [49]
The wealthiest known late Avar cemetery consisted of 52 documented graves. However, the original grave number was higher (probably did not exceed a hundred) since sand extraction earlier destroyed many burials. The burial ground is considered fully excavated and composed of 23 male, 14 female and 14 sub-adult graves. The cemetery remains a unique phenomenon in the late Avar period because of the high number of burials with horse-riding and weapon-related equipment, the high quality of material culture and the relatively high proportion of precious metal artefacts. Furthermore, the Hortobágy-Árkus series is one of those cemeteries which connects the Avar age and the Hungarian Conquest period since the cemetery was dated between the late 7th and early 10th centuries CE. The cultural aspects of the cemetery are very much different from that of the other rural communities of the late Avar period in the Carpathian Basin. They probably represented an elite group in fact, the only identified elite in the late period of the Avar age. We involved seventeen samples in our study.

**ARK-6: Grave No. 6:**
It was the disturbed grave of a sub-adult individual (infantia I). The burial did not contain any known grave goods, and it was dated to the late Avar period (8th century CE?).

**ARK-11: Grave No. 11:**
The disturbed grave (especially the chest region) contained the skeletal remains of a young adult male. Traces of coffin were registered in the pit, but the burial did not contain any grave finds. It was dated to the 8th-9th centuries CE.

**ARK-14: Grave No. 14b:**
The grave was heavily disturbed, and a ritual cause for the disturbance was identified. At the north wall of the pit, the complete skeleton of a young-adult female (14a) was unearthed. The deceased was in a position upside down with the skull and the arms under the body. Gold foil ornaments of the headdress, two gold band rings, gold earrings and silver armbands were found at the skeleton. Besides, a single skull and some bone fragments of a young adult male (14b) were found on the bottom of the pit. The precious metal artefacts and the position of the remains provide a strong argument that the woman was probably killed and thrown into the secondary opened grave pit. Based on the well-datable objects, the female burial was dated to the first half of the 9th century CE. However, radiocarbon dating of the skeletons (14a: 700–780 CE with a 66% probability; 14b: 760–880 CE with a 66% probability) resulted in different dates. In this case, however, the archaeological dating of the golden artefacts seems

to be more trustworthy since the relative position of the skeletal remains contradicts the relative age suggested by the radiocarbon dating.

**ARK-16: Grave No. 16:**
The skeletal remains of a young adult male were found in the disturbed grave. Traces of a coffin were registered in the pit, but the burial did not contain any grave finds. It was dated to the 8th-9th centuries CE.

**ARK-17: Grave No. 17:**
It was the disturbed grave of a sub-adult (infantia II) individual. The burial did not contain any known grave goods. It was dated to the late Avar period (8th century CE?).

**ARK-19: Grave No. 19:**
The grave contained the skeleton of a juvenile with male characteristics and without any grave finds. The burial was dated to the 8th-9th centuries CE.

**ARK-20: Grave No. 20:**
It was the burial of a sub-adult (infantia I) individual. The grave contained two melon seed-shaped beads at the skull. It was dated to the late Avar period (8th-9th centuries CE).

**ARK-21: Grave No. 21:**
The burial contained the skeleton of a juvenile with female features. The bones were covered by a brownish layer, probably by leather remains. The grave finds were golden earrings. The archaeological material was dated to the end of the late Avar period or early 9th century CE, supported by radiocarbon dating.

**ARK-24: Grave No. 24:**
The skeleton of a young adult male was found in the grave. The disturbed bones were in a layer mixed with burnt charcoal and ash. The high-prestige deceased was buried in a coffin with a horse, a belt with gilded cast brass ornaments and a saber. The burial was dated to the last phase of the late Avar period (supposed absolute date: 770-800 CE). Besides, radiocarbon analysis resulted in a longer, partially overlapping interval concerning the probable dating of the skeleton (680-780 CE with 66% of probability).

**ARK-29: Grave No. 29:**
The partially destroyed skeleton of a juvenile with male features was found in the grave. The high-prestige individual was buried in a coffin with a horse and a belt with gilded cast brass ornaments. The burial was dated to the second half of the late Avar period (i.e., between 760-810 CE). In addition, radiocarbon analysis dated the skeletal remains to a longer, partially overlapping interval (680-780 CE with 66% of probability).

**ARK-36: Grave No. 36:**
The skeleton of a young adult male was found in the destroyed grave, in a wooden chamber. The burial contained the skeleton of a horse but did not contain any known, datable grave goods. The grave was dated to the Avar period based on radiocarbon data (660–770 CE with 66% of probability).

**ARK-38: Grave No. 38:**
It was the grave of a young adult female buried in a coffin. The remains of a wooden vessel, repaired with a strip made of non-ferrous metal (copper alloy). The burial was dated to the 8th-9th centuries CE.

**ARK-41: Grave No. 41:**
The disturbed grave contained the skeleton of a young-adult male with a bronze buckle. The burial was dated to the early phase of the late Avar period (first half of the 8th century CE).

**ARK-43: Grave No. 43:**
The skeletal remains of a young adult female were found in the grave. The upper body parts were disturbed. The burial did not contain any grave finds. It was dated to the 8th-9th centuries CE.

**ARK-48: Grave No. 48:**
It was the strongly disturbed grave of an adult female. Only the lower parts of the legs were intact. In the disturbed area, a fragment of an earring and a bone needle case were found. The burial was dated to the second half of the late Avar period (late 8th century, early 9th century CE).

**ARK-49: Grave No. 49:**
The skeleton of a young adult male was found in the disturbed grave. The burial contained the skeleton of a horse, heart- and shield-shaped gilded-silver belt mounts with plant motifs, bone handle fragments of a knife (?), textile pieces, and fragments of bark or thin wood object. It was dated to the late 9th and first half of the 10th century CE.

**ARK-50: Grave No. 50:**
The strongly disturbed grave contained the skeleton of an adult male and a horse. A bone bow plate and a belt mount were found in a secondary position. The assemblage was dated to the late 9th and first half of the 10th century CE.

**S1.4.5 Budapest–Csepel-Kavicsbánya (CS; Pest County)** [50–52]

**CS-465: Grave No. 465:**
In 1924, the remnants of a lonely rich male burial were found in the Csepel gravel mine. The skull of a middle-adult (45-50-year-old) male taxonomically showed Mongoloid (Tungid) characteristics. Among the grave finds, a cast bronze buckle, a bronze nail, arrowheads, a sword with gold-plated P-shaped suspension loop, and iron coffin clips were placed in the Hungarian National Museum. The analogies of the sword are known from the Kunpeszér grave No. 3. Based on the radiocarbon dating of this grave, the Csepel grave finds can be dated between 630–650/660 CE.

**S1.4.6 Csólyospálos-Felsőpálos (CSPF; Bács-Kiskun County)** [53,54]

In the outskirts of Csólyospálos, 258 Avar graves were partially excavated by Mihály Kőhegyi and Erika Wicker between 1983 and 1987. The cemetery was founded probably by warriors and their families. Animal bones, potteries and eggs were also found in the cemetery. Based on the archaeological material, the burial ground was opened in the 7th century CE (around 650), and the last graves can be dated to the 8th century CE (around 750). Anthropological analysis was carried out on the fragmented skeletal remains of 244 individuals. Concerning the sex and age-at-death data, 156 adults and 88 sub-adults were described, and 73 males and 76 females were identified in the series. According to the results of the morphological analysis, 10% of the skulls showed Mongoloid characteristics. However, the low quality of the bones highly limited the evaluation and interpretation of the data. We involved four samples in our study.

**CSPF-37: Grave No. 37:**
The skeleton of a male individual with Mongoloid characteristics was in the NW–SE orientation grave. The adult male was buried into a coffin and wore a roof-shaped hair clamp and a belt decorated with

round-shaped mounts. Besides, a shield-shaped bronze buckle, an iron buckle and a knife were found. The grave was dated to the 7th century CE (around 670).

**CSPF-114: Grave No. 114:**

An adult male with Mongoloid characteristics was buried in the NW–SE orientation grave. Three-three postholes stood out from the line of the pit wall registered in the large, square-shaped grave pit. The belt was decorated with silver hemispherical fittings, smaller and larger belt ends. A bow with middle-wide arms was placed on the left side of the dead. The quiver contained four arrows located on the right leg bones. A one-edged iron sword was between the left elbow and left knee. In addition, cattle bones were found between the knees. The tomb can be dated to 650–670 CE.

**CSPF-182: Grave No. 182:**

The skeletal remains of an adult male were in the NW–SE orientation grave. The grave goods were an iron buckle, a knife, a bone stick-end, and three three-edged, narrow arrowheads. The burial was dated between 670–700 CE.

**CSPF-213: Grave No. 213:**

The skeletal remains of an adult male were found in the NW–SE orientation, large, square-shaped pit grave. Small bead pendant earrings, two iron buckles and two knives were found in the grave. Besides, the quiver was next to the left leg, and a bone stick-end was next to the right leg. The bow with wide arms was smashed and placed to the left side (near the elbow and knee). The tomb was dated around 700 CE.

**S1.4.7 Dunavecse-Kovacsos dűlő (DK; Bács-Kiskun County)** [55]

During a preventive excavation of the M8 motorway on the outskirts of Dunavecse town, two early Avar period graves (feature 701 and 702) were found by Andrea Lantos 50 m from each other in 2004–2005. We involved grave No. 701 in our analysis.

**DK-701: Grave No. 701:**

The feature was an NW–SE orientation prestigious burial. An adult male was buried with a belt decorated with pressed silver mounts, sword, bow, and quiver. In addition, a Byzantine jug was put near the head of the dead. The horse-riding equipment (stirrups, bit, head and breast collar, and breeching) was put on the chest of the dead, while the saddle decorated with bone mount was imposed on the foot. The grave was dated to 630–650/660 CE.

**S1.4.8 Fajsz-Garadomb (FGD; Bács-Kiskun County)** [56]

Eight early Avar-age graves, arranged in two irregular and loose lines, were excavated on the outskirts of Fajsz village by Mihály Kőhegyi in 1962. The cemetery, mainly composed of armed male burials, was the burial site of a large family or genus belonging to the middle class or leadership of the Avar society. The skeletons showed Mongoloid characteristics, and the dead were buried between 600–650/660 CE. We involved one sample in our analysis.

**FGD-4: Grave No. 4:**

Skeletal remains of a middle-adult male with Mongoloid characteristics were found in the grave. The individual wore silver earrings and a belt decorated with pressed silver pseudo buckle mounts. Furthermore, a bow and a quiver with arrowheads were put in the grave. The burial was dated to the 7th century CE, around 630.

**S1.4.9 Felgyő-Ürmös tanya (FU; Csongrád-Csanád County)** [57,58]
On the outskirts of the Felgyő village, Gyula László excavated 216 Avar graves between 1960 and 1977. The cemetery is not fully excavated. Data on the known graves indicate that the cemetery was established in the 7th century CE (around the 630s). Furthermore, the burial site was used until the end of the Avar period. From the 216 graves, only the skeletal remains of 166 individuals were available for anthropological analysis. Altogether 58 sub-adults and 108 adults were described, and 49 females and 60 males were determined. Concerning the taxonomic features, the series is composed of skulls with Europid characteristics (e.g., Mediterranean, Cromagnoid). Besides, several skulls showed both Europid and Mongoloid features, and in six cases the Mongoloid characteristics were dominant. We involved two samples in our study.

**FU-193: Grave No. 193:**
The skeleton of an adult male was found in the NNW–SSE orientation. The burial did not contain any known grave goods. The burial was dated indirectly based on the surrounding graves dated to 670–700 CE.

**FU-215: Grave No. 215:**
The NW–SE orientation grave contained the skeleton of a young adult male with Mongoloid characteristics. The grave goods were silver earrings, earrings with small spherical pendants, iron buckles, Fönlak-type belt mounts, iron ring, a knife, and a bone clasp. A bow with narrow arms was placed on the dead, and sheep vertebrae were also found in the grave. The grave can be dated between 630–650 CE, based on the Fönlak-type belt fittings.

**S1.4.10 Hajós-Cifrahegy (HC; Bács-Kiskun County)** [59,60]
169 Avar graves were excavated between 1978 and 1983 by Mihály Kőhegyi. The site is situated 1,5 km southwest of Hajós-Pincefalu. The cemetery is not fully excavated, but its boundaries were found in the western, eastern and northern directions. The cemetery was used from 650 to 810/820 CE. The skeletal remains of 135 individuals were distinguished during the anthropological analysis led by Antónia Marcsik. The number of the sub-adults was significantly lower (34) than the number of adult individuals (101). Concerning the sex ratio, a higher proportion of females (59%) compared to males (41%) was registered in the series. Based on the results of metric and taxonomic analysis, the population was relatively homogeneous and mostly composed of people with Europid characteristics. Mongoloid features were described only in two cases. We involved one sample in our analysis.

**HC-168: Grave No. 168:**
The skeletal remains of a juvenile (with female features) were found in the NW–SE orientation grave. The grave goods consisted of beads, iron bracelets, a fragment of a bronze chain, a bone comb, a biconical spindle whorl, and a hand-formed vessel. Based on the finds, the grave was dated to the 7th century CE (around 650–660).

**S1.4.11 Homokmégy-Halom (HH; Bács-Kiskun County)** [61,62]
A total of 167 graves of an Avar age cemetery was saved by Gyula László and Nándor Fettich in 1936. However, the burial ground is considered partially excavated, and according to the estimations, only one-third of the graves were found/ saved. The material was not fully published yet. The most important findings in the material are the belt sets (with plate and cast belt ornaments), silver and bronze earrings, bracelets, different types of beads, pottery (e.g., vessels), and animal bones. The cemetery was dated to the end of the 7th and the 8th centuries CE. The anthropological material, 77 skeletons, was analyzed by Pál Lipták. He identified six sub-adults and seventy-one adults, and he described 37 males and 34 females in the series. The skulls dominantly showed Europid characteristics

(Mediterranean types, Dolichocran and Brachycran groups), and Mongoloid features appeared only in three cases. We involved three samples in our analysis.

**HH-10: Grave No. 10:**
A young adult male with Europid characteristics on the skull was found in the grave. The burial contained cast bronze belt ornaments with griffon and tendril motifs. It was dated to the 8th century CE.

**HH-22: Grave No.22:**
The skeletal remains of a young adult male with Europid characteristics on the skull were found in the grave. The burial did not contain any known grave goods. It was dated to the 7–8th centuries CE.

**HH-102: Grave No. 102:**
The skeleton of a young adult male with Mongoloid skull characteristics was found in the burial. The grave goods were an iron buckle, a knife, plate belt ornaments, an iron ring, sabretache closure ornament, and a bone disentangling hook. It was dated to the last third of the 7th and beginning of the 8th centuries CE.

##### S1.4.12 Jánoshida-Tótkérpuszta (JHT; Jász-Nagykun-Szolnok County) [63,64]

A late Avar cemetery with 259 graves was excavated 4 km far from the Alattyán-Tulát Avar period cemetery. The archaeological material is composed of cast belt fittings (only a few pressed fittings were found), earrings, a string of beads (e.g., melon seed-shaped beads), bronze bracelets, spindle-whorls, needle cases, pottery (e.g., vessels), and animal bones. Weapons were found only in a few cases (swords in four graves and archery equipment in one grave). The cemetery was dated to the end of the 7th and 8th centuries CE. The anthropological material was investigated by Sándor Wenger. Only 46 skulls and 18 postcranial skeletons were available for the analysis, from which he described nine sub-adults and thirty-seven adults, and he identified 15 female and 23 male individuals. Based on the cranial properties, he distinguished three typological groups. Most of the skulls showed Europid characteristics (Nordoid, Mediterranean and Baltic types), but in certain cases, Europo-Mongoloid features were also detected. We involved three samples in our analysis.

**JHT-30: Grave No. 30:**
It was the disturbed grave of a young adult male. The burial contained a set of bronze cast belt ornaments with tendril motifs, the bone grip of an unknown tool and a bone end-ornament of a staff. It was dated to the end of the 7th and beginning of the 8th centuries CE.

**JHT-130: Grave No. 130:**
The skeletal remains of a middle-adult male were found in the grave with animal bones and a knife (lost). The burial was dated to the 8th and 9th centuries CE.

**JHT-154: Grave No. 154:**
The skeleton of a young adult male with an iron buckle (lost) was found in the burial. It was dated to the end of the 7th and 8th centuries CE.

##### S1.4.13 Kaba-Dögös / Kaba-Bitózug (KD; Hajdú-Bihar County) [65]

Near the road between the villages Kaba and Tetétlen, a cemetery dating to the middle and late Avar period was excavated in 1974-1975. The documented 160 graves formed several rows in three groups. The graves were oriented to the N–S directions. Although two burials were unearthed with horse-riding-related equipment, the archaeological material of the cemetery was composed of simple artefacts, and only several graves contained more prestigious inventory (e.g., pressed or cast belt sets or jewelry made from gold or silver). The cemetery was dated to the 8th century CE. The anthropological analysis of the skeletal remains is in progress. We involved two samples in our analysis.

**KD-16: Grave No. 16:**
A middle-adult male individual with Europid (Cromagnoid) characteristics was buried in the grave. A pressed bronze belt set with undecorated and griffin-decorated strap ends was found in the pit. The burial was dated to the 8th century CE.

**KD-29: Grave No. 29:**
It was the burial of a young- or middle-adult male with Europid characteristics. The burial contained a cast bronze belt set, an iron buckle and a silver finger ring. The grave was dated to the 8th century CE.

##### S1.4.14 Kiskundorozsma-Daruhalom-dűlő (KDA; Csongrád-Csanád County) [66,67]

In 2003 and 2004, three Avar age cemeteries were found close together during the preventive excavations of highways M5 and M43. The burial grounds were near the present-day town of Kiskundorozsma, at the western and eastern banks of the Maty Creek. The Kiskundorozsma-Daruhalom-dűlő cemetery is considered fully excavated, and 93 burials were found. The graves formed four parallel lines, and they were oriented mainly in NW-SE directions. Traces of wooden coffins were registered in 62% of the pits, besides remains of other wooden constructions were also documented. Animal bones were found in 57% of the burials. Implements (e.g., knives) and pottery (e.g., vessels) were the most frequent types of grave goods in the series. Female graves contained belt buckles, a string of beads (e.g., melon seed-shaped beads) and different hoop-jewelry (e.g., finger rings, bracelets). Besides, male burials were characterized by similar types of jewelry and sets of ornamented belts. The cemetery was dated to the Middle Avar period, the last third of the 7th century CE, and close relationships were supposed with the population of the Kiskundorozsma-Kettőshatár I and II series. We involved four samples in our analysis.

**KDA-188: Grave No. 188:**
The skeleton of a young adult female with Europid characteristics was in the grave. Traces of a coffin, bronze earrings, a string of beads, and an iron sickle were found in the pit. The burial was dated to the last third of the 7th century CE.

**KDA-485: Grave No. 485:**
It was the niche grave of an old adult male with Europid characteristics on the skull. The traces of a coffin were documented in the pit. Besides, bronze and silver earrings, a vessel, an egg, sheep bones, a simple belt with an iron buckle, a knife, and fire-lighting tools were found. The burial was dated to the last third of the 7th century CE.

**KDA-517: Grave No. 517:**
The skeletal remains of a young adult male were found in the grave with traces of a coffin and an iron buckle. The burial was dated to the last third of the 7th century CE.

**KDA-520: Grave No. 520:**
The burial contained the skeleton of a middle adult female with Europid characteristics on the skull. The grave contained a spindle-whorl and the skeleton of a chicken. It was dated to the last third of the 7th century CE.

**S1.4.15 Kiskundorozsma-Kettőshatár I (KK1; Csongrád-Csanád County)** [66,67]

The cemetery was found near the Daruhalom-dűlő burial ground, at the eastern bank of the Maty Creek. Although the site has been partially excavated, 298 graves and an external ditch were unearthed, and the cemetery is considered one of the largest Avar age cemeteries in the county. Concerning the grave pits, remains of wooden constructs and graves with a niche were documented in some cases. In addition, traces of wooden coffins were found in 34% of the burials and animal bones (food offerings) were found in a high number of cases. Among the different types of grave goods, knives, belt buckles and spindle-whorls were found most frequently in the graves. Besides, the female burials contained beads, bronze or silver earrings, hair rings, and bracelets. Frequent artefacts of male burials were the belt sets with bronze or silver ornaments. The cemetery was dated from the end of the 7th CE to the beginning (first third) of the 9th century CE. Based on archaeological and anthropological data (e.g., high frequency of cases with leprosy in both series), a close relationship is suspected between the populations of the Daruhalom-dűlő and Kettőshatár I cemeteries. We involved five samples in our study.

**KK1-245: Grave No. 245:**

The burial contained the skeleton of a middle adult male with Europid and Mongoloid characteristics (Uralian) on the skull. A simple belt with iron buckle, a knife, and the skeleton of a chicken were found in the grave. It was dated to the end of the 7th and 1st third of the 8th century CE.

**KK1-251: Grave No. 251:**

The skeletal remains of a middle adult male with Mongoloid skull characteristics were found in the grave. The burial contained a belt with iron buckle and hoops, a knife, the skeleton of a chicken, and leg bone of a sheep. It was dated to the end of the 7th and 1st third of the 8th century CE.

**KK1-252: Grave No. 252:**

The disturbed grave contained the skeleton of a middle adult male with Mongoloid skull characteristics. The skeleton of a chicken, partial skeletons of a cattle and a fish, a vessel, fire-lighting tools, and a belt with iron buckle and bronze plate ornaments were found in the burial that was dated to the last third of the 7th century CE.

**KK1-368: Grave No. 368:**

It was the niche grave of a young adult female with Europid characteristics. The burial contained bronze earrings, a string of beads, and the skeleton of a chicken. It was dated to the 1st third of the 8th century CE.

**KK1-541: Grave No. 541:**

The burial of a middle adult male with Europid characteristics contained a cattle bone, bronze earrings, a knife and a belt with bronze ornaments. It was dated to the last third of the 7th century CE.

**S1.4.16 Kiskundorozsma-Kettőshatár II (KK2; Csongrád-Csanád County)** [66,67]

The third Avar age cemetery from Kiskundorozsma was found on a hilltop, was separated from the Kettőshatár I burial ground by about 60m distance. Therefore, Kettőshatár II was separated from the other two cemeteries concerning their physical location. Further differences were registered in the anthropological (e.g., the dominance of Mongoloid characteristics) and archaeological characteristics. Forty-three graves were unearthed, and the site is considered fully excavated. The grave pits (e.g., depth and length) were significantly larger compared to the average size in this period. Remains of wooden constructs and burial chambers were registered in several cases, and traces of wooden coffins (e.g., metal parts) were found in every grave. In addition, grave robbery was documented in 100% of the burials that damaged both the skeletal remains and the grave goods. Nevertheless, some artefacts still represent the original wealth of the cemetery (e.g., golden plate strap ends with tendril motifs, gilded bronze rosette-shaped ornaments, gold earrings, and gilded silver shield-shaped belts ornaments. The cemetery was dated to the second half of the 8th century CE, and it was used parallel with the Kettőshatár I cemetery. We involved four samples in our analysis.

**KK2-429: Grave No. 429:**
The robbed grave contained the skeleton of a young adult female with Mongoloid characteristics on the skull. A dog skeleton was found in the pit. Besides, a vessel, sheep bones, a knife, and a simple belt were unearthed. It was dated to the 2nd half of the 8th century CE.

**KK2-441: Grave No. 441:**
It was the robbed niche grave of a middle adult female with Mongoloid characteristics on the skull. The burial contained gold earrings, rosette-shaped gold dress fittings, a vessel, and bones of two sheep, cattle, and a dog. It was dated to the 2nd half of the 8th century CE.

**KK2-445: Grave No. 445:**
The skeletal remains of an old adult male with Europid characteristics on the skull were found in the robbed niche grave. A vessel and elements of a bow were documented in the burial, dated to the 2nd half of the 8th century CE.

**KK2-670: Grave No. 670:**
The robbed grave contained the skeleton of a middle adult male with Mongoloid characteristics on the skull. Traces of a coffin were registered in the pit. Besides, the remains of a bow, a vessel, and bones of a sheep and a horse were found in the burial. It was dated to the 2nd half of the 8th century CE.

**S1.4.17 Kunpeszér-Felsőpeszéri út, Homokbánya (KFP; Bács-Kiskun County)** [68–70]
Thirty-two graves were excavated at the outskirts of Kunpeszér village by Elvira H. Tóth. Fifteen graves were dated to the early Avar period (group I), and the others were dated to the 8th century CE. We involved group I in our study. The early Avar graves were on a large area that was a loose burial site. Three sub-adults and twelve adults were described, and six males and six females were identified in the series. Most of the individuals showed Mongoloid taxonomic features. The male burials were rich since they were buried with belts decorated with silver and gilded mounts, swords decorated with gold and silver plates, bow, quiver, and arrowheads. The female graves were less wealthy, and their burials contained only simple grave goods. We involved four samples in our analysis.

**KFP-6: Grave No. 6:**
An adult male with Mongoloid characteristics was buried in the NNW–SSE orientation grave. The grave goods were belts decorated with silver shield-shaped and gilded bronze rosette-shaped mounts, a bow, a quiver made of birch bark, and ten arrowheads. The grave was dated between 630–660 CE.

**KFP-7: Grave No. 7:**
The grave was oriented to NNW–SSE and contained the skeleton of an adult female. The burial yielded a few beads, an iron awl, a knife, a clay spindle-whorl, and a pair of rounded, gilded bronze Mezőszilas-type pendants. The grave was dated to 630–660 CE.

**KFP-30a: Grave No. 30a:**
The skeletal remains of an adult male with Mongoloid characteristics were found in the NNW–SSE oriented grave. The southern half of the pit was later in superposition with grave 30b. Silver earrings with a small spherical pendant, silver hair-clamps, belt decorated with silver round-shaped mounts, iron buckles, a bow, quiver with arrows, and a sword decorated with P-shaped silver ears were in the grave. The burial was dated between 630–660 CE.

**KFP-31: Grave No. 30b:**
The irregular-shaped pit was dug into grave No. 30a. The skeleton of an adult female with Mongoloid characteristics and traces of trauma on the skull was found. The deceased was lying on her left side in a contracted position (the right leg was less pulled up to the knees than the left). At the right thigh, a prism-shaped bone clamp with five holes at the narrower side was found. Based on the superposition, the woman was buried later than the individual 30a. The grave was dated to the middle of the 7th century CE (around the year 650).

**S1.4.18 Kiskőrös-Pohibuj Mackó dűlő (KPM; Bács-Kiskun County)** [71,72]
The archaeological site was south of Kiskőrös town, located in the middle region of the Danube–Tisza Interfluve, between the railroad and highway towards Kecel. Unfortunately, only a small part of the cemetery was explored since the diversity of the recovered material suggests the uninterrupted, long period use of the cemetery. Early excavations started in the 1890s, and by 1936 we had information on the material of 62 Avar graves. The earliest burials can be dated from the 630s CE, the latest from around 800 CE. The anthropological material was analysed by Pál Lipták. He could identify two sub-adults (juveniles) and ten adults, and he described three females and seven males in the population. The series contained both skulls with Europid and Mongoloid characteristics. We involved three samples in our analysis.

**KPM-14: Grave No. 14:**
There is no information about the grave that contained the skeleton of a young adult male with Europid characteristics and Mongoloid skull features. Only a grey, wheel-formed vessel is known from the burial. It was dated to the 2nd third of the 7th century CE (between 630 and 650).

**KPM-23: Grave No. 23:**
There is no information about the grave of a middle-adult male with Europid characteristics and Mongoloid skull features. Cylindrical braid ornaments, quadrangular cast bronze belt mounts with griffin-shaped decorations, a knife, and a wheel-formed yellowish-brown, narrow-necked jar were found. The grave was dated to the 2nd third of the 8th century CE (between 720–750).

**KPM-27: Grave No. 27:**
There is no information about the grave of a young-adult female with Mongoloid characteristics on the skull. The burial contained a biconical spindle-whorl. The fine dating of the findings was not achievable.

**S1.4.19 Kiskőrös-Vágóhídi dűlő (KV; Bács-Kiskun County)** [73,74]
Sixty-seven graves arranged in seven burial groups of a larger Avar-age cemetery were excavated on the outskirts of Kiskőrös town between 1934–1938. The population of the cemetery showed Mongoloid taxonomic features, however, the anthropological data on the skeletal remains of eleven

individuals were published by János Nemeskéri. The analysed material was composed of one sub-adult and ten adults. Besides, two females and eight males were identified in the material. Based on the numerous prestigious objects and the gold dress fittings, gold ornaments of coffins and shrouds, the cemetery is among the wealthiest middle Avar-age cemeteries of the Carpathian Basin. Assumingly, it was used by the community of the power elite. The site was dated between 660–720 CE. We involved four samples in our analysis.

**KV-3367: Grave No. LXXI (57):**
The skeletal remains of an adult male with Mongoloid (Tungid) characteristics were found in the grave. The deceased was laid to rest in a coffin decorated with gold plates. Headgear decorated with glass inlaid quadrangular gold fittings, cylindrical gold hair-clamps and gold hook around the head were found in the burial. The grave was dated between 670–700 CE.

**KV-3369: Grave No. I:**
It was the grave of a middle-adult male with Mongoloid (Sayanic) skeletal features. The burial contained headgear decorated with gold mounts and a belt ornamented with round-shaped mounts with glass fittings in the middle. Besides, a silver chalice was near the dead. Unfortunately, a sword was kidnapped from the grave. The deceased was buried between 660–700 CE.

**KV-3450: Grave No. IX (8):**
The skeleton of a middle-adult female was in the grave. Silverplate strap-end decorated with lily and gold foil ornaments of a coffin or shroud were found in the burial. The findings were dated approximately between 660–720 CE.

**KV-3456: Grave No. XLV (43):**
A middle-adult female with Mongoloid (Tungid) characteristics was found in the grave. Iron coffin clamps and a bronze earring with a blue bead pendant were found. The grave was dated to the 1st third of the 8th century CE (700–720).

**S1.4.20 Makó-Mikócsa-halom (MM; Csongrád-Csanád County)** [75,76]
During the preventive excavation of a factory on the outskirts of Makó, a complete early Avar-age cemetery with 251 graves was excavated by Csilla Balogh in 2009–2010. Based on the burial customs (NE–SW orientation, niche and end-niche graves, large number of partial sacrificial animals), the cemetery seems to be used by a population of Eastern European origin who joined the Avars in the Eastern European steppe region. Most of the skulls in the population showed Europid characteristics. Besides, some skulls had Mongoloid features. The cemetery was used by the community between 568–650 CE. We involved seven samples in our analysis.

**MM-61: Grave No. 61 (Obnr 218/ Snr 227):**
The niche grave contained the skeletal remains of a middle-adult male with Europid characteristics. The grave goods were a belt and shoes decorated with metal mounts, a sword decorated with silver plates, a bow, a quiver with arrowheads and some lamella of an armor. Besides the right leg of the dead, tools for wood, bone/horn and metal processing (e.g., saw, rasp) and uniquely, in a crucible made of iron plate semi-finished bone bow-applications were found. The grave can be dated to 580–620 CE.

**MM-131: Grave No. 131 (Obnr 388/ Snr 409):**
The skeleton of a middle adult male with Europid (Pamyrian) characteristics on the skull were found in the catacomb grave. The burial contained the partial skeletons of a horse and a sheep (ca. ten years old), a bone awl, closure ornament of a sabretache, horse-riding equipment (stirrup and girth buckle). It was dated to the 1st half of the 7th century CE.

**MM-240: Grave No. 240 (Obnr 518/ Snr 547):**
It was the niche grave of an old adult male with Mongoloid skull characteristics. The wealthy burial contained animal bones in high number, i.e., the whole skeleton of a horse with harness, partial skeletons of two horses, two cattle, a calf, and two sheep. Besides the stirrup and the bit, the horse-riding equipment contained sponge-shaped harness mounts and plain strap ends. In addition, implements of goldsmithing, bronze rivets and raw materials, a knife, remains of a wooden box, a dagger with a P-shaped suspension handle were unearthed. Elements of two belts were found (two iron buckles) on the skeleton, one of them was decorated with small, pressed rounded fittings and the other with pressed Martynovka type mounts. The grave was dated to 600–630 CE.

**MM-245: Grave No. 245 (Obnr 525/ Snr 554):**
The niche grave contained the skeleton of a middle adult male with Mongoloid skull characteristics. The grave goods of the wealthy burial were partial skeletons of a horse and a sheep, horse-riding equipment (bit, girth buckle, stirrup) with bronze harness ornaments, a bone hook, a Byzantine bronze measuring cup, implements of goldsmithing, and a set of belt with gilded silver Martynovka type ornaments. The burial was dated to 600–630.

**MM-80: Grave No. 80 (Obnr 256/ Snr 268):**
The skeletal remains of a young adult male with Mongoloid skull characteristics were recovered from the niche grave. The skeleton of a horse with rosette-shaped silver harness ornaments, and partial skeletons of a cattle, a calf, and a sheep were found in the pit. The grave goods were a belt with silver ornaments of North Italian origin and weapons, i.e., a sword and a bow. The burial was dated to 600–630 CE.

**MM-83: Grave No. 83 (Obnr 259/ SNR 271):**
It was the burial of a sub-adult individual (7–8 years old). A vessel and a string of beads were found in the niche grave. It was dated to the 1st half of the 7th century CE.

**MM-151: Grave No. 151 (Obnr 410/ Snr 434):**
The skeleton of an old adult female was found in the grave that was constructed with two narrow ledges along the longer sides of the pit. Assumingly, these were to hold the wooden planks that functioned as a coffin. The partial skeletons of a cattle and a sheep were recovered from the pit. In addition, a whetstone, a spindle-whorl, a bone needle case, and an iron buckle were found. The burial was dated to the 1st half of the 7th century CE.

**S1.4.21 Mélykút-Sáncdűlő (MS; Bács-Kiskun County)** [77,78]
Altogether 54 graves were excavated northeast of Mélykút in 1959 and 1968. In particular, 46 graves were dated to the Avar period and eight burials to the Hun period. The Avar graves were in a 100 x 50 m area located at the NW–SE part. The burials formed five irregular grave groups of different sizes. Based on the archaeological findings, the cemetery was used from 625/630 CE to 700/710 CE. The skeletal remains of 45 individuals were available for anthropological analysis. Antónia Marcsik described ten subadults and thirty-five adults in the material. Besides, she identified twenty-two females and seventeen males in the population. The series is composed of skulls with Europid characteristics. In an additional case, Mongoloid characteristics were described. We involved three samples in our analysis.

**MS-43: Grave No. 43:**
The skeleton of an adult female was found in the grave oriented in NNW–SSE directions. The burial contained silver earrings with a large spherical pendant, an iron buckle, a knife, a disc-shaped spindle-

whorl, and a wheel-formed pottery jar. Furthermore, cattle and chicken bones were found near the knees. The grave was dated 630–660 CE.

**MS-45: Grave No. 45:**
It was the burial of an adult male, oriented in the NNW–SSE directions. Silver earrings with a spherical pendant, an iron buckle, a belt decorated with pressed silver mounts, a knife, an iron ring, and a bard-box decorated with carved bone plates were found in the grave. The grave was dated between 630-660 CE.

**MS-50: Grave No. 50:**
The skeletal remains of an adult male were found in the NNW–SSE oriented grave. The burial contained iron buckles, rectangular-shaped pressed belt fittings, a pressed strap-end decorated with medallion, a strap-end decorated with a masque, an iron ring, an axe, and hand-made pottery. The grave was dated approximately to the 670s CE.

**S1.4.22 Madaras-Téglavető (MT; Bács-Kiskun County)** [79,80]
The cemetery, located in the outskirts of the Madaras village in the southern part of the Danube–Tisza Interfluve (30 km far from the Danube valley), was excavated between 1952 and 1962. However, ninety-two, Avar-period graves were found, the cemetery was not fully excavated. The known part of the cemetery was used in a relatively short period, between 650–720 CE. In the archaeological material of the cemetery, the ratio of the weapon founds was high, and sabers, bows and quivers decorated with bone plates were found. Based on the material, the cemetery was considered as the legacy of armed commoner people. The skeletal remains of eighty-eight individuals were available for anthropological analysis. Pál Lipták and Antónia Marcsik determined thirty sub-adults and fifty-eight adults, and they identified thirty-six males and twenty-four males in the material. Concerning the taxonomic data, the series was composed of skulls with Europid, Mongoloid and Europo-Mongoloid characteristics. In particular, a difference was detected between the sexes, for most of the male skulls were described with Europid characteristics, while Mongoloid features were frequently documented among the female skulls. We involved four samples in our analysis.

**MT-17: Grave No. 17:**
It was a burial of a middle-adult male, oriented in the NW–SE directions. Hair-clamps made from bronze and gold plates, earrings, a saber, arrows, iron buckles, a knife, and animal bones were found. The grave was dated between 670–700 CE.

**MT-23: Grave No. 23:**
The skeletal remains of a middle-adult male were found in the grave oriented to NW–SE directions. The grave goods were silver earrings, silver hair-clamps, silver belt mounts, an iron buckle, a knife, a bow, a saber, and five arrows. The assemblage dated the burial between 670–700 CE.

**MT-29: Grave No. 29:**
The skeleton of a middle-adult male was found in the grave oriented in the NW–SE directions. The burial contained an earring, a long iron knife, an iron buckle, strap-ends cut from undecorated bronze plates, belt mounts, and a quiver decorated with carved bone plates. The burial was dated between 650–670 CE.

**MT-74: Grave No. 74:**
The grave oriented in the NNW–SSE direction contained the skeleton of a middle-adult male. Pressed bronze four-lobed belt mounts, pressed round-shaped ornaments, mounts cut off from a bronze plate,

a knife, and arrowheads in a quiver decorated with carved bone plates were found. The burial was dated between 670–700 CE.

##### S1.4.23 Orosháza-Béke tsz., Homokbánya (OBH; Békés County) [81,82]

A middle-sized late Avar cemetery with 149 graves was partially excavated at the northern outskirt of Orosháza in 1967–1969. However, a significant part of the graveyard was destroyed by sand mining. The graves were oriented to the NW-SE directions. Certain burial customs, namely six graves with an end-niche and six burials with cattle bones are related to the Early Avar population of the Tiszántúl region. Iron nails and clasps were found in several graves, indicating that the deceased was buried in a coffin. Eight graves contained a belt set, and valuable painted yellow mugs were found in two burials. The cemetery was dated to the last third of the 7th century and the 8th century CE. We involved two samples in our analysis.

**OBH-37: Grave No. 37:**

The skeletal remains of a middle-adult female with Mongoloid taxonomical features were found in the end-niche grave. The burial contained animal bones and an iron knife. Based on the location and form of the grave, it was among the earliest burials of the cemetery.

**OBH-52: Grave No. 52:**

It was the burial of a young-adult female individual with Europid characteristics on the skull. The grave contained a fragmented bronze earring with a bead pendant. It was ornamented with granulations that were soldered to the bottom of the ring. The burial was dated to the 8th century CE.

##### S1.4.24 Orosháza-Bónum Téglagyár (OBT; Békés County) [81,82]

The cemetery was excavated south from Orosháza in 1966-1967. A total of 231 graves were unearthed, and 14 additional burials were destroyed before the excavation (only their findings were saved). Concerning the burial customs, five end-niche graves, nine horse burials – separated horse graves too – and traces of burial structures were documented. Fourteen graves contained belt sets, while one burial contained a sword. In addition, bronze phaleras with lion heads were found in a destroyed burial with horse-riding-related deposits. The site was mainly used from the last third of the 7th century CE to the end of the 8th or beginning of the 9th century CE. The archaeological material indicated that the population was composed of descendants of the original, Early Avar period population and the newcomers. We involved five samples in our study.

**OBT-3: Grave No. 3:**

The skeleton of a middle-adult male was found in the disturbed grave. The burial contained a set of belt fittings (e.g., shield-shaped mounts, buckles and strap-ends) ornamented with a tendril motif. It was dated to the 8th century CE.

**OBT-51: Grave No. 51:**

It was the burial of a middle-adult male with Mongoloid (Sinid) features. The grave goods were a sword and a cast bronze belt fitting with floral ornaments. The burial was dated to the end of the 7th century CE.

**OBT-56: Grave No. 56:**

The skeletal remains of a young adult male with Europid characteristics (Brachycran and some Pamirid features) were found in the grave. Wood remains with bronze nails were next to the skull. Besides, a knife, an egg and a cast bronze belt set were documented in the grave. The belt fittings were ornamented with griffins. The burial was dated to the 8th century CE.

**OBT-106: Grave No. 106:**
The burial contained the skeleton of an old adult male with Europid (Cromagnoid) characteristics on the skull. The bones of a larger animal (horse or cattle) were found above the deceased. In addition, food offerings and an iron buckle were documented in the burial. The grave could not be dated more precisely in the 7-8th centuries CE.

**OBT-108: Grave No. 108:**
The skeleton of a middle-adult female with Europid characteristics (with gracile Mediterranean features) was in the grave. The burial contained a biconical spindle-whorl and two iron buckles. The grave could not be dated more precisely in the 7-8th centuries CE.

**S1.4.25 Pitvaros-Víztározó (PV; Csongrád-Csanád County)** [82,83]
A middle-sized Avar cemetery with 225 graves was excavated southeast to Pitvaros in the 1990s. The burials were oriented to the NW-SE or NNW-SSE directions. Regarding the burial customs, graves with end-niche and burials with horse-riding-related deposits (i.e., two graves with horse burials) were documented in the cemetery. Besides, sheep bones were found in more than thirty graves. Twenty-five male individuals were buried with a silver or gilded bronze belt set. Another precious finding of the cemetery was in one of the horse burials that contained an ornamented harness with numerous bronze phaleras and other fittings. The use of the cemetery started in the last third of the 7th century CE and ended in the early 9th century CE. Archaeological data indicates that the community was composed of descendants of the original, Early Avar period population and the newcomers. The skeletal remains of 226 individuals were analysed by Erika Molnár. The generally low quality of the bones limited the evaluation of the data. She distinguished 66 sub-adults and 160 adults and identified 85 females and 85 males in the series. The material contained skulls mostly with Europid characteristics. However, several cases with Mongoloid features were registered too. We involved five samples in our study.

**PV-12: Grave No. 12:**
The skeleton of a young adult male with Europid characteristics (Cromagnoid) was found in the pit. Traces of a coffin (nails) were found in the grave. Besides, the burial contained a bronze belt set with griffin ornamented mounts. It was dated to the first half of the 8th century CE.

**PV-116: Grave No. 116:**
The grave contained the skeleton of a young adult female The grave goods were fragments of a bronze earring, a bronze earring with a glass bead ornament, and animal bones: the skeleton of cattle, sheep and hen bones. The burial was dated to the 8th century CE based on radiocarbon data.

**PV-125: Grave No. 125:**
It was the grave with an end-niche of a middle-adult male individual. Parts of horse-riding equipment (i.e., two stirrups, an iron bit and bronze harness ornaments) were found in the entrance pit. Besides, a yellow, wheel-formed pot was in the niche, and sheep bones were registered in the grave. The burial was dated to the second half of the 7th century CE. The dating was verified with radiocarbon data.

**PV-200: Grave No. 200:**
A middle-adult male with Mongoloid characteristics was buried in the elongated, rectangular-shaped grave, oriented to NW-SE directions. Skeletal remains of a harnessed stallion were found at the south-eastern end of the pit. The human remains laid at the north-western part of the grave, two-two post-

holes were observed on both sides of the skeleton. The deceased was buried in a wooden coffin strengthened with clamps.
Sheep bones were found next to the skull. In addition, a set of belt mounts were part of the inventory. The bronze belt fittings were cast with a tinned surface which was common in the 1st third of the 8th century CE. This date was verified with radiocarbon analysis as well.

**PV-205: Grave No. 205:**
The grave with an end-niche contained the skeleton of a middle-adult male with Europid characteristics and some Mongoloid features on the skull. The niche itself was disturbed. Concerning the burial customs, nails of the coffin and the skeleton of a foal were found. The grave finds were a set of bronze belt fittings. The background of the belt fittings was decorated with punched ornaments, and their surface was tinned. It was dated to the beginning of the 9th century CE; thus, this burial is one of the latest graves in the cemetery.

**S1.4.26 Sükösd-Ságod (SSD; Csongrád-Csanád County)** [84,85]
Three hundred-sixty-nine Avar period graves were excavated by Mihály Kőhegyi, György V. Székely and Erika Wicker near Sükösd village between 1967 and 1981. However, the cemetery was not fully excavated, and the archaeological material is only partially published. Based on the findings, the cemetery was opened after the 630's CE, and it was used continuously until around 750 CE. The finds of the cemetery are related to the Avar cemeteries of Eastern Transdanubia. The skeletal remains of 165 individuals were available for anthropological analysis. Altogether 56 sub-adults and 109 adults were identified. Besides, 68 females and 41 males were described in the series. According to the result of the taxonomic investigation, skulls with Europid characteristics (e.g., Cromagnoid, Mediterranean) are dominant in the material, and Mongoloid features were registered only in one case. We involved six samples in our study.

**SSD-17: Grave No. 17:**
It was the NW–SE oriented burial of an adult female with Mongoloid features on the skull (Pamirian-type). The grave goods were an earring with mounted spherical pendant, an earring with a large spherical pendant decorated with plate overlay, the bone mouthpiece of a hose, monochrome beads, an iron buckle, a knife, a biconical spindle-whorl, and hand-made pottery. The grave was dated between 630–660 CE.

**SSD-35: Grave No. 35:**
The skeletal remains of a middle-adult male with Mongoloid (Sinid-type) characteristics were found in the grave oriented in the NW–SE directions. The features of the grave and the grave goods are unknown.

**SSD-58: Grave No. 58:**
The grave oriented in the NW–SE directions contained the skeleton of a middle-adult female with Iranian characteristics on the skull. Earrings with a large spherical pendant, an iron buckle, a knife, and a biconical spindle-whorl were found in the burial. The grave was dated between 630–660 CE.

**SSD-144: Grave No. 144:**
A middle-adult male was found in the grave oriented in the NW–SE directions. The burial contained an iron buckle, pressed round-shaped belt fittings, undecorated large and small strap-ends, and a knife. The grave was dated approximately to the 670s CE.

**SSD-151: Grave No. 151:**

In the NW–SE orientation grave, the skeleton of a middle-adult male with Iranian characteristics was found. The grave goods were belt fittings cut off from a bronze plate, an iron buckle, a knife, and a wheel-formed pottery jar. The burial was dated between 670–700 CE.

**SSD-198: Grave No. 198:**
The grave oriented in the NW–SE directions contained the skeletal remains of a sub-adult individual without any known grave goods. The fine dating of the burial was not achievable.

**S1.4.27 Szeged-Fehértó A (SZF; Csongrád-Csanád County)** [86,87]
A middle-sized Avar age cemetery with 376 graves was excavated by Ferenc Móra and Károly Cs. Sebestyén between 1929 and 1932. Based on the orientation of the graves, two groups were distinguished in the cemetery, an eastern and a western group. Furthermore, differences were registered in the grave goods between the eastern and the western parts. The burials of the eastern group contained belts with pressed ornaments, weapons (archery equipment, swords, axe), vessels (e.g., hand-formed types) jewelry, such as earrings (e.g., earring with glass bead pendant), bracelets or strings of beads. On the other hand, artefacts characteristic of the late Avar period were found in the burials of the western part, such as belt mounts with griffon and tendril motifs and melon seed-shaped beads. The cemetery was dated to the last third of the 7th and 8th centuries CE. The skeletal remains (mostly skulls) of 204 individuals were available for analysis. In the fragmented material, 25 subadults and 176 adults were distinguished, and 88 females and 89 males were identified. Concerning the taxonomic data, the series composed of skulls with Europid characteristics (e.g., Brachycran groups, Cromagnoid, Mediterranean) and Mongoloid features were documented only in three cases. We involved four samples in our analysis.

**SZF-26: Grave No. 26**
It was the burial of a young adult male with Europid characteristics on the skull. The grave goods were antler/ bone bow plates, arrowheads, a sword, bone mouth-piece of a leather bottle, ornamented bone plates, silver earrings with hollow globe-pendants, glass beads, silver plate belt fittings, silver buttons, a knife, a vessel, and animal (pig) bones. It was dated to the 3rd quarter of the 7th century CE.

**SZF-43: Grave No.43**
Skeletal remains of an old adult male with Europid (Cromagnoid) characteristics on the skull were found in the grave. The burial contained archery equipment (bow plates, arrowheads), belt buckles, iron and silver plate belt mounts, fire lighting tools, a knife, and animal bones. It was dated to the 7th century CE.

**SZF-181: Grave No. 181**
The skeleton of a middle-adult female with Mongoloid features on the skull was found in the grave with glass beads. It was dated to the 7–8th centuries CE.

**SZF-371: Grave No. 371**
It was the burial of a middle-adult female with Europid (Brachycran) characteristics on the skull. The grave goods were silver earrings with hollow globe-pendants, glass beads around the head, an incomplete bronze ring, iron buckles, an ornamented narrow bone plate, iron chain-links, egg fragments, and animal (pig) bones. It was dated to the 2nd quarter of the 7th century CE.

**S1.4.28 Szeged-Kundomb (SZK; Csongrád-Csanád County)** [88,89]

After brickmakers found burials in 1924, 319 burials dated to the Avar age were recorded in several excavation periods. However, the cemetery was not fully explored. The archaeological material represented the wide scale of burial customs (e.g., graves with a niche) and artefacts characteristic for the whole Avar period. The grave goods composed of weapons (e.g., archery equipment, swords), ornamented belts, earrings (e.g., earrings with different types of pendants), beads, bracelets, finger-rings, tools and implements (e.g., knives, fire lightning tools, spindle-whorls, awls), coins (e.g., Roman coins), and pottery (e.g., vessels and bottles). The cemetery was dated from the last third of the 7th to the 9th centuries CE. The skeletal remains (mostly skulls) of 179 individuals were available for anthropological investigation. According to the result, 33 sub-adults and 146 adults were distinguished, and 72 females and 62 males were identified in the population. Concerning the taxonomic analysis, only skulls with Europid characteristics (e.g., Cromagnoid, Brachycran, Nordoid, Mediterranean) were described in the series. However, in certain cases, Mongoloid features were also registered. We involved five samples in our study.

**SZK-83: Grave No. 83**
The skeletal remains of a middle-adult male with Europid characteristics (Armenoid) were found in the grave. The burial contained bronze earrings (with missing pendants), a bronze belt buckle with shield-shaped plate, silver plate belt mounts, fire lighting tools, a knife, and glass beads. It was dated to the 2nd half of the 7th century CE.

**SZK-102: Grave No. 102**
It was the burial of a middle-adult male with Europid characteristics (Mediterranean). Traces (metal parts) of a wooden coffin were registered in the pit. The grave goods were Roman coins emitted by Constantinus Gallus II (351–354), Valens (364–378) and Constantinus II (337–361), fire lightning tools, a knife, gilded silver strap ends and silver belt mounts (e.g., shield-shaped mounts), silver plate fragments, a propeller-shaped iron plate, an iron buckle, and chicken bones. The burial was dated to the middle of the 7th century CE.

**SZK-130: Grave No. 130**
The burial contained the skeleton of a young adult male with Europid characteristics (Mediterranean-Pamirian). A vessel, a wheel-made yellow bottle, a set of bronze belt mounts (e.g., shield-shaped ornaments, mounts with pendants, propeller-shaped mounts, belt-hole guards) ornamented with tendril motifs, a knife, and a fowl bone was found in the grave. It was dated to the middle of the 8th century CE.

**SZK-180: Grave No. 180**
The skeleton of a young adult male with Europid characteristics was found in the grave. The burial contained a silver earring with a hollow globe-pendant, a sword, a long knife, archery equipment (bow plates, arrowheads), fire lighting tools, a set of bronze and silver belt ornaments (e.g., shield-shaped mounts), and strap ends. Besides, an iron buckle, S-shaped bronze chain-links, a Roman knee-brooch, Roman coins (the only preserved coin was emitted by Constantine II), a three-discs-shaped bone plate, a hand-made vessel, and animal (chicken and pig) bones were found. It was dated to the 2nd half of the 7th century CE.

**SZK-213: Grave No. 213**
It was the burial of a middle-adult female with Mongoloid features. The grave goods were a silver earring with a hollow globe-pendant, a rock-crystal bead, an iron buckle, a knife, a biconical clay spindle-whorl, and animal (pig and chicken) bones. The burial was dated to the middle of the 7th century CE.

##### S1.4.29 Székkutas-Kápolnadűlő (SZKT; Csongrád-Csanád County) [90,91]

The only fully excavated large-scale cemetery was found by K. B. Nagy in the vicinity of Székkutas. The site was next to an extensive prehistoric rampart which was probably used in the Avar period too. The excavation was carried out between 1965 and 1986, and 534 graves were recorded. The cemetery contained 59 graves with end-niche. Two horse burials and three graves with horse-riding equipment were found. Besides, other animal bones as food offerings (sheep and cattle) were also documented. These animal bones were found in a non-anatomical position, which is characteristic for the last third of the 7th century CE. The average middle and late Avar age material occurred in the series. However, the more prestigious objects (e.g., belt sets, clasps) are more common in the 8th century CE. A sword and arrowheads were also registered. Several burials contained typical early Avar period material. However, these objects were broken and repaired. Based on the archaeological material, L. Bende and G. Lőrinczy dated the use of the cemetery between the middle or second half of the 7th century and the 9th century CE. The skeletal remains of 518 individuals were available for anthropological analysis from which 147 sub-adults and 371 adults were distinguished. Besides, 188 females and 155 males were identified in the series. We involved five samples in our analysis.

**SZKT-62: Grave No. 62:**

A young adult male with crossed legs was found in the grave, probably wrapped into some textile. The grave did not contain any known grave goods; thus, the fine dating of the burial is not achievable.

**SZKT-70: Grave No. 70:**

The skeletal remains of a young adult male were found in the grave. Traces of a wooden coffin (metal parts) were documented in the pit. The grave contained unknown iron fragments, a knife, belt buckles, and a set of belt mounts with ornaments (e.g., griffin and tendril motif). The burial was dated to the 8th century CE.

**SZKT-89: Grave No. 89:**

It was the burial of a young adult female. She was buried with a painted yellow mug, a pair of bronze earrings with dark-blue bead pendants, four melon-seed shaped beads, and a biconical spindle-whorl. The burial was dated to the 8th century CE.

**SZKT-265: Grave No. 265:**

An old adult male was buried in the grave. The remains (iron nails) of a wooden coffin were documented. The skeleton of a cattle and the skull and legs of a sheep were found above the deceased. Besides, the burial contained two iron buckles, a knife and further animal bones. The grave could not be dated more precisely.

**SZKT-311: Grave No. 311:**

The skeleton of an old adult male was found in the grave. The remains of a wooden coffin were documented. A whole skeleton of a cattle and bones of a sheep were found between the deceased and the pit wall, but the bones were not in anatomical order. The grave contained an iron ring, a knife and two iron buckles. The burial was dated to the end of the 7th century CE.

##### S1.4.30 Szeged-Makkoserdő (SZM; Csongrád-Csanád County) [92,93]

An Avar age cemetery with 267 graves was unearthed in 1930 and 1942. However, the site is considered partially excavated. The burial customs of a grave pit with a niche, using wooden coffins, food offerings, horse burials are characteristic of the cemetery. Similarly to the Szeged-Kundomb series, the archaeological material contains the variety of the typical Avar period grave goods types, i.e., weapons (e.g., archery equipment, sword), jewelry (e.g., earrings with small bead-pendants, penannular hair rings, hairplait-ornaments, melon seed-shaped beads, and bracelets), dress fittings (e.g., sets of belt fittings), horse-riding equipment (e.g., stirrups, bits), pottery (e.g., hand-made and wheel-made vessels), and implements (e.g., fire lightning tools, knives, disentangling hooks). The cemetery was dated from the last third of the 7th to the beginning of the 9th century CE. The skeletal remains (mostly skulls) of 182 individuals were saved and available for anthropological investigation. Among them, 41 sub-adults and 111 adults were identified, and 52 females and 58 males were distinguished. The series contained mostly skulls with Europid characteristics (e.g., Cromagnoid, Brachycran groups, Mediterranean). However, several cases (15.8%) with Mongoloid features were also described. We involved five samples in our study.

**SZM-24: Grave No. 24:**
It was the grave with a niche of a young adult male with Mongoloid characteristics on the skull. Animal bones, i.e., a horse burial, calf and foal bones were found in the burial. The grave goods were a set of bronze and silver belt fittings, silver hairplait-ornament and Byzantian silver coins emitted by Konstans II (641–668) and Konstantinos IV (668–685). The burial was dated to the last third of the 7th century CE.

**SZM-38: Grave No. 38:**
It was the burial of an adult male with Europid characteristics and Mongoloid features on the skull. The grave with a niche contained a horse burial, horse-riding equipment, gold earrings with pendants, penannular silver earrings, bronze and silver belt fittings, a short and a long knife, fire lighting tools, and animal bones (chicken and calf). The burial was dated to the middle of the 7th century CE.

**SZM-255: Grave No. 255:**
The skeletal remains of a middle adult male with Europid characteristics on the skull were found in the grave with a niche. The burial contained iron buckles, fragments of a Byzantine buckle, a fragmentary glass bead, a vessel, and chicken bones. It was dated to the second half of the 7th and beginning of the 8th centuries CE.

**SZM-259: Grave No. 259:**
The skeleton of an adult male was found in the grave. The grave goods were bronze earrings with glass bead ornaments, a string of beads, melon seed-shaped beads, a knife, an iron buckle, and animal bones. The burial was dated to the 7th century CE.

**SZM-332: Grave No. 332:**
It was the burial of a sub-adult (infantia II) individual. The grave did not contain any known grave goods, and it was dated to the first half of the 8th century CE.

##### S1.4.31 Szegvár-Oromdűlő (SZOD1; Csongrád-Csanád County) [94–96]

On a range of sandhills, the 467 graves of an early Avar period cemetery were excavated. The archaeological data is partially published. The archaeological material was characterized by the grave pits with a niche, the great number of animal bones (horse, cow and sheep), hand-made pottery and the typical early Avar period grave-good types (e.g., pressed breast discs, pressed belt and harness fittings). Besides, solidus emitted by Heraclius (610–641) and Heraclius Constantinus (reigned in 641) were also found in the graves. The cemetery was dated from the last third of the 6th century CE to the last third of the 7th century CE. The skeletal remains of 298 individuals were available for anthropological investigations. The cemetery contained sub-adult burials in a high ratio (ca. 64,4%), and 192 sub-adults and 106 adults were identified. Concerning the result of the sex determination, 82 females and 51 males were described in the series. Taxonomic analysis was not performed due to the low qualitative preservation of the bones. Therefore, the given data in the descriptions below are the results of the re-examination by our anthropologist colleagues. We involved four samples in our analysis.

**SZOD1-76: Grave No. 76:**
It was the burial of an adult male with Europid (Cromagnoid) characteristics on the skull. The grave did not contain any known grave goods, and it was dated to the 1st half of the 7th century CE.

**SZOD1-127: Grave No. 127:**
The skeletal remains of a young-adult female with Mongoloid characteristics and an artificially deformed skull were found in the grave without known grave goods. The burial was dated to the 1st half of the 7th century CE.

**SZOD1-187: Grave No. 187:**
It was the burial of a middle-adult female with Mongoloid characteristics and an artificially deformed skull. The grave did not contain any known grave goods. The burial was dated to the 7th century CE.

**SZOD1-554: Grave No. 554:**
The skeleton of a middle-adult male with Europid characteristics was found in the grave with a niche. The burial contained the partial skeleton of a cattle and bones of three sheep. It was dated to the 2nd quarter of the 7th century CE.

**SZOD1-829: Grave No. 829:**
The grave with a niche contained the skeleton of an adult female (though it turned out to be genetically a male) with Europid characteristics. The grave goods were a knife, an earring, a spindle-whorl, a tube, and an earpick. Besides, the bones of cattle were found in the pit. It was dated to the 2nd quarter of the 7th century CE.

**S1.4.32 Szarvas-Grexa téglagyár (SZRV; Békés County)** [41,54,97]
Four hundred twenty-two graves were found during the excavation led by Irén Juhász in 1983, 1984 and 1986. Concerning the burial customs, horse burials, graves with niche and food offerings (animal bones and vessels) were documented in the cemetery. The archaeological material was composed of jewelry, dress and belt fittings (sets of belt mounts, buckles), weapons (archery equipment), horse-riding equipment (e.g., stirrups), and implements. One of the most emblematic findings of the graves is a needle case decorated with runiform writings. The cemetery was dated to the 7-9th centuries CE. The skeletons of 423 individuals were involved in the anthropological analysis. One hundred and nineteen sub-adults and 299 adults were identified, and 166 females and 160 males were described in the series. Although the low qualitative preservation of the bones limited the evaluation of the taxonomic data, the material contained skulls both with Europid and Mongoloid characteristics too. We involved eight samples in our analysis.

**SZRV-54: Grave No. 54:**
A middle-adult female with Mongoloid features was buried in the grave with end-niche. The grave goods were a pot with a handle, a necklace, a pair of silver earrings with globular pendant, a spindle-whorl, an iron buckle, and tweezers. It was dated to the mid-7th century CE, the earliest period of the cemetery.

**SZRV-67: Grave No. 67:**
The skeletal remains of an old-adult female were in the grave. A spindle-whorl and a bone needle case with runiform writings on the surface were found. The burial was dated to the 8th century CE.

**SZRV-147: Grave No. 147:**
It was the burial of a middle-adult male. The grave contained horse-riding equipment (a pair of stirrups and an iron bit), a vessel beneath the skull and pressed bronze belt fittings (e.g., rectangular mounts with griffin ornaments). The burial was dated to the second half of the 7th century CE.

**SZRV-168: Grave No. 168:**
The skeleton of a young adult male was found in the grave. Cattle bones, an undecorated bronze buckle and an openwork small strap end were found in the burial. It was dated to the end of the 7th century CE.

**SZRV-212: Grave No. 212:**
The grave contained the skeletal remains of a sub-adult individual (6-8 years old) without any known grave goods. The burial was dated to the 8th century CE.

**SZRV-266: Grave No. 266:**
It was the grave of a middle-adult male buried with a belt ornamented with a set of cast bronze mounts. The mounts were decorated with floral motives. The grave was dated to the 8th century CE.

**SZRV-277: Grave No. 277:**
The skeletal remains of a middle-adult male with Europid characteristics were found in the grave. The burial contained an iron buckle and a knife. It was dated to the 8th century CE.

**SZRV-316: Grave No. 316:**
The skeleton of a young adult female with Mongoloid characteristics on the skull was found in the grave. The burial did not contain any known grave goods. Based on the location of the grave, it was dated to the 8th century CE.

##### S1.4.33 Tiszafüred-Majoros-halom (TMH; Jász-Nagykun-Szolnok County) [98]
An extensive cemetery with 1283 graves was excavated by Éva Garam at the outskirt of Tiszafüred between 1965 and 1972. It is the largest known Avar period burial site east of the Tisza River. Among the graves, 70 separated horse burials with horse-riding equipment were found. The grave goods of the male burials were composed of weapons (i.e., sabres, axes, spearheads, or antler bow plates), pressed and cast belt fittings. The most characteristic types of female jewelry were the perforated cast bronze discs, which hung off from the belt. Besides them, various earrings, beads, arm- and finger rings were found. In addition, a stamp mould for belt fittings with bird ornaments was also excavated. The cemetery had close analogies with the iconic cemetery of Zamárdi-Réti-földek (Somogy County) from Transdanubia. The Tiszafüred-Majoros-halom series was dated between the last third of the 7th century and the beginning of the 9th century CE. The anthropological data is not published yet. We involved six samples in our study.

**TMH-199: Grave No. 199:**
It was the burial of an adult male with Europid characteristics (Nordoid-x) and Mongoloid skull features. A belt set with cast bronze belt fittings, a bronze chain, a knife, and two iron buckles were found in the grave. It was dated to the Late Avar period.

**TMH-388: Grave No. 388:**
The grave contained the skeleton of an adult male with Europid (Nordoid and Cromagnoid) characteristics. The burial inventory consisted of a belt set of pressed bronze fittings and a knife. The grave was dated to the Middle Avar period.

**TMH-509: Grave No. 509:**
The skeletal remains of an adult male with Europid (Cromagnoid-A) characteristics were found in the grave. The grave was at some distance from the other burials and contained a belt set of cast bronze fittings, an iron awl and two iron buckles. It was dated to the latest phase of the late Avar period.

**TMH-756: Grave No. 756:**
An adult female with Europid (Brachycran) characteristics was found in the grave. The burial contained an iron buckle, a spindle-whorl, a knife, a handmade pot, and a comb. The fine dating of the burial was not achievable.

**TMH-798: Grave No. 798:**
The disturbed grave contained the skeletal remains of an individual with unknown sex (we could show to be a male based on genetic data) and Europid characteristics with Mongoloid features on the skull. The iron nails of a wooden coffin were documented in the pit. A bronze buckle was the only registered artefact in the grave. The fine dating of the burial was not achievable.

**TMH-1273: Grave No. 1273:**
It was the burial of an adult female. The grave contained a pair of bronze earrings, glass beads, and a perforated bronze disc. The grave was dated to the end of the 7th century CE.

##### S1.4.34 Tatárszentgyörgy-Szabadrétpuszta (TTSZ; Pest County) [70,99]

The site was at a dune, near the road between Tatárszentgyörgy and Kunpeszér. Ilona Kovrig excavated 51 graves in 1951. The cemetery was used between the middle of the 8th and the beginning of the 9th centuries CE. Compared to other series from this period, the cemetery contained burials with weapons in high number (11 of the 18 male burials). The weapon findings were composed of archery equipment, swords and long knives. In addition, the archaeological material was characterized by the presence of gilded bronze shroud plates, gilded bronze belt fittings and gilded bronze jewelry. Based on the findings, the cemetery was used by a late Avar period military elite population. However, the anthropological analysis was highly limited due to the disturbance and robbery of the burials that damaged the bones, and especially the skull. Fourteen skulls were available for investigation, from which three sub-adults and eleven adults were identified, and eight females and three males were described. The series contained skulls with Europid and Mongoloid characteristics too. We involved one sample in our analysis.

**TTSZ-43: Grave No. 43:**
The skeleton of a young adult female with Mongoloid features on the skull was found in the grave, oriented in the NW–SE directions. Cast bronze lobed-shaped belt fittings, a knife, a vessel, and sheep bones were found in the burial. It was dated to the end of the 8th and beginning of the 9th centuries CE.

#### S1.5 General archaeological description of the Hungarian Conquest Period

The nomadic power structure of the Magyars from the eastern steppe region appeared at the Lower Danube region in the 830s CE [100]. Their armies, consisting of mounted archers, were hired in many battles by the different European kingdoms. During their campaigns in the second half of the 9th century CE, they reached even the Carpathian Basin (e.g., in 862 and 881 CE). So it is possible, that small groups might have already settled down there before the occupation of the new territory under the leadership of Álmos and his son Árpád at the turn of the 9th and 10th centuries CE (based on historical sources, it was dated between 895 and 900 CE) [101].

After the fall of the Avar Khaganate at the beginning of the 9th century CE, the Carpathian Basin was not empty, but it had a peripheral status. The Trans Danube region was controlled by Franks, remnants of the Avar population lived at the Great Hungarian Plain, Bulgarians occupied Transylvania, and Slavic tribes were at the northern and southern peripheries [101,102]. The lack of unity and a sole power centre allowed the Magyars to conquer the Carpathian Basin in a short time and to integrate these local populations into their society. The conquering Hungarians established a complex military and political system strong enough to unite and control the whole Carpathian Basin, which ultimately resulted in the foundation of the Christian Hungarian Kingdom at the turn of the 10th and 11th centuries CE. The new kingdom was ruled by the descendants of Árpád, i.e., the members of the Árpád-dynasty for three centuries.

Historically the Conquest period lasted just for a few decades, however the archaeological material cannot be dated with such accuracy, so most of the 10-11th century material is considered the Hungarian Conquest Period horizon [103,104]. Until now, more than 30.000 graves were dated to this period [105]. Earlier, historians assumed that the Conquerors counted several hundred thousand people, however, the relatively low number of excavated graves suggests that their ratio could be much smaller [106].

In the 10th century CE, new burial customs appeared in the Carpathian Basin. These included different types of partial horse burials with harness, various categories of jewelry and dress fittings (e.g., earrings with cast-beadrow pendant, cast opened-work braid disc ornaments), weapons (e.g., sabers, the components of the bows, ornamented bow cases,), so-called prestigious artefacts (e.g., belt with mounts), and horse trappings (e.g., pear-shaped stirrups with flat arches). Their analogies can be found east from the Carpathian Basin, for example in the Ural region (Türk, 2014), in the Volga region [107], or the Caucasus (Lezsák, 2020). These mostly appeared as individual burials or burial grounds with a relatively small number of graves and were usually considered as the 'classical' or 'typical' 10th-century-CE Hungarian burial horizon [105,108].

On the other hand, large cemeteries with a small percentage of burials with horse-riding-related or weapon-related grave goods are also part of the horizon. These series are characterized by the generally low number of grave goods, simpler jewelry types, and the presence of the coins emitted by the Kings of the Árpád-dynasty. Certain cemeteries were continuously used in the Avar and Hungarian Conquest periods (e.g., the multi-period cemetery of Vörs-Papkert B). Besides, in some cases, burial customs showed analogies with the material of earlier periods, for instance, with late Avar period cemeteries (e.g., the graves with a niche in the 10–11th-century cemetery of Homokmégy-Székes) supporting that the integration of the local populations was indeed successful.

Archaeological investigation revealed regional differences concerning the characteristics of the cemeteries and artefacts [105,109]. The wealthiest sites dated to the 10th century CE were found in the highest number at the Upper Tisza region (e.g., Karos I, II, and III, Kenézlő–Fazekaszug) that suggests a vertically centralized system with a sole power centre in this region [105,108]. Later, in the last third of the 10th century CE, it was moved next to the Danube, to the central region of Buda and Esztergom.

Main cultural changes occurred at the end of the 10th and in the 11th centuries CE, basically related to the spread of Christianism and the reforms by king Stephen I (1000–1038). The pagan grave goods and burial customs (e.g., weapons and partial horse burials) started to vanish, and new, uniformed types of jewelry appeared in the graves (e.g., beads with gold and silver foils, S-terminalled hair rings). Many cemeteries that were used in the 10th century have been closed and new burial grounds were opened in the 11th century. Besides, continuous 10–11th-century-CE cemeteries (e.g., Püspökladány-Eperjesvölgy, Ibrány-Esbóhalom, and Alba Iulia-Izvorul Împăratului) are also known. The distribution of these types of cemeteries show regional differences suggesting inner migrations and population changes [105]. However, these were rather long-term changes that occurred by the end of the 11th and beginning of the 12th centuries under the rule of king László I (1077–1095) and king

Kálmán I (1095–1116). Their reign ultimately meant the end of the old era, and the new Christian traditions became widespread in the whole kingdom.

In the present study we adapted the cemetery grouping systems of Kovács (Kovács, 2013) and Gáll (Gáll, 2019), which are based on general characteristics, e.g., number of burials, burial customs, ratio of graves with partial horse burials, weapons, prestigious grave goods, jewelry, dress fittings, and artefacts made of precious metals. As a result, we distinguished two groups labelled as 'elite', characterized by the lower total number of burials, higher ratio of graves with partial horse burials, weapons, prestigious grave goods, jewelry, dress fittings, and artefacts made of precious metals) and 'commoner', characterized by the higher total number of graves in the cemetery, higher number of graves without grave goods, lower ratio of graves with partial horse burials and weapons, and simpler jewelry and dress fittings. A grave was attributed to the Árpádian period if it could be unambiguously dated to the 11th-century often distinguished by Christian type graves without pagan grave goods, and coins minted by the early Árpádian kings.

#### S1.6 Descriptions of the studied 10-11th-century cemeteries and graves

##### S1.6.1 Algyő-258. kútkörzet (AGY; Csongrád–Csanád County) [110,111]

Eighty-three graves were excavated by Béla Kürti between 1973 and 1976. The site is considered fully excavated. Although the archaeological material is only partially published, two groups were distinguished (northern/middle group and southern group) based on the distribution of the graves and grave goods. Different types of implements (knives, equipment for fire lightning) and jewelry of the head (e.g., hair rings, earrings with cast-beadrow pendant, and braid ornaments), neck (leaded cross), and arm and hand (bracelets and rings) have been found most frequently in the graves. However, the material is characterized by the clothing ornaments in the female burials and the high number of graves with horse-riding- (21 graves) and weapon-related (18 graves) artefacts. Based on the archaeological material, the cemetery was dated to the 10th century CE. The anthropological material was analyzed by Antónia Marcsik and her colleagues. The skeletons of 77 individuals were available and fit for the analysis. They recorded a significant male surplus (41:20) in the series, which was explained by the possible military status of the population. Only Europid types of skulls were described in the series. However, the low state of preservation limited interpretations. Further chemical-analytical examinations were carried out on the material by Imre Lengyel. Similarly to the archaeological analysis, he distinguished two groups in the cemetery (northern and southern groups).
Classification of the cemetery: 10th-century quarter cemetery (type 4) in the Kovács system; group A in the Gáll system. In our analysis, the Algyő-258. kútkörzet samples are categorized as Conqueror elite.
Four burials dated to the 10$^{th}$ century CE were involved in our analysis:

**AGY-49: Grave No. 49:**
Located in the northern part of the cemetery. The burial of an adult female with a Europid type of skull contained a partial horse burial with horse-riding equipment. Additionally, animal bone (considered as a food offering), hair rings, bracelets, and clothing ornaments were found.

**AGY-75: Grave No. 75:**
It has been found in the northern group, near to grave No. 49. A middle-adult male with a Europid type of skull was buried in the grave with a bronze bracelet.

**AGY-87: Grave No. 87:**
It was in the middle group of the cemetery. The adult female individual with a Europid type of skull was found in the grave. Animal bone (considered as a food offering) and a hair ring were found in the grave during the excavation.

**AGY-92: Grave No. 92:**
Located in the southern group of the cemetery. The burial of a middle-adult male with a Europid type of skull was among the richest in the series and contained partial horse burial, horse riding-related equipment, archery equipment, animal bone considered as a food offering, gold hair ring, and silver funerary eye piece.

**S1.6.2 Bugyi-Kisványpuszta /Bugyi-Felsővány (BK; Pest County)** [112]
Local people with metal detectors have found 10th-century stray founds (e.g. gilded silver dress fittings with pendants, belt mounts, S-terminalled hair rings), indicating that many burials were destroyed because of intense agricultural field works. In 2011, twenty graves were found during a rescue excavation, but most of these did not contain any known grave goods. The jewelry was composed of simple types of hoops around the head (e.g., penannular hair rings), finger rings (band rings), and bracelets (e.g., wire bracelets). Weapons (i.e., archery equipment) were found in two graves, besides one burial contained horse bones and horse-riding equipment was found in two graves. However, grave No. 2 made the cemetery unique since this is the first burial with a sabretache plate in the region. The cemetery was dated to the 2nd half of the 10th century CE. The results of the anthropological analysis are not published yet, and only data on the skeleton from grave No. 2 is known. The series was classified as group B in the Gáll system and as Conqueror elite in our study. We involved one sample, the burial No. 2, in our analysis.

**BK-2: Grave No. 2:**
The skeletal remains of a young adult male with Europid characteristics and Mongoloid features were found in the grave. The grave goods were partial horse burial, horse-riding equipment, archery equipment, mounted belt with gilded silver ornaments, and gilded silver sabretache plate. The burial is considered as the richest in the cemetery and the micro-region.

**S1.6.3 Szeged-Csongrádi út (CSU; Csongrád–Csanád County)** [111,113]
The excavation was carried out by the leading of Béla Kürti between 1974 and 1987. The cemetery consisted of 13 graves which were dated to the 10th century. Three groups were distinguished based on the location of the graves (western, middle, and eastern groups). Jewelry (hair rings, bracelets, string of beads) and clothing ornaments (ball buttons, pressed ornaments, and rhombic shaped ornaments) were found in the subadult and female graves. However, two graves did not contain any inorganic material. The male burials have more importance concerning the quality and the quantity of the grave goods: archery equipment, fire lightning tools, partial horse burials, horse-riding equipment, knives, hair rings, sabres, a sword with sabre hilt, and coins (the coin of Constantine VII Porphyrogenitus & Romanos II /945–953/ and the coin of Hugh of Provance & Lothar II /931–945/) were found. The cemetery was dated to the 10th century CE, and especially on the 2nd half of it. The anthropological analysis was carried out by Antónia Marcsik. The preservation of the bones is low that highly limited the possible conclusions and our sampling. A total of 12 skeletons were available for the research, from which five males, five females, and two subadults were described. According to the taxonomical analysis, the Europid types are dominant in the population. In addition, mongoloid morphological characteristics were detected on three skulls. Based on the archaeological material and the distribution of the graves, it is considered as a burial ground for a few families. Classification of the cemetery: 10th-century quarter cemetery (type 4) in the Kovács system; group A in the Gáll system. One sample was involved in our analysis categorized as Conqueror elite:

**CSU-11: Grave No. 11:**
The burial of an adult female with Mongoloid morphological characteristics on the skull. The grave did not contain any known grave goods. The grave was dated to the 10$^{th}$ century CE.

##### S1.6.4 Homokmégy-Székes (HMSZ; Bács–Kiskun County) [114,115]

The excavation of the site was carried out by Zsolt Gallina and Sándor Varga between 1996-2002. The cemetery consisted of 206 graves. It is divided into northern and southern parts based on the type and the orientation of the graves, the grave goods, and the position of the arms. The northern part is further split into east and west sides based on the density of the graves. Due to the characteristics of the soil, the shape of the grave pits has been preserved well: several grave pits with sidewall niches have been found, as well as traces of other Avar age burial customs, which are rare in the 10th century (e.g., patterns of post-holes in the grave pit). The archaeological findings include jewelry and clothing ornaments, for example, hoops around the head (e.g., S-terminalled and penannular hair rings, earrings), neck jewelry (e.g., neckrings, beads), and arm jewelry (e.g., bracelets, rings), dress fittings, as well as weapons (archery equipment, axe) and implements (e.g., fire-lighting equipment, knife).
Based on the composition of the findings, the cemetery dates to the period from the first third of the 10th century CE to the first third of the 11th century CE.
The state of preservation of the anthropological material is generally good or medium, excluding the sub-adult skeletons. During the anthropological examinations, 136 adult and 50 sub-adult remains were distinguished, of which 63 was described as male and 83 as female. Based on the distribution of taxonomic features, in addition to the predominance of Europids (88%), a smaller proportion of components belonging to the Mongoloid race also appears (9.3%). Based on biological distance calculations, the population shows similarities to other 10th century and Avar-era series.
Classification of the cemetery: 10th-century village cemetery (type 5) in the Kovács system; group B2 in the Gáll system. In our analysis, the samples are categorized as Conqueror commoner.
Nine burials dated to the 10–11th centuries CE were involved in our analysis:

**HMSZ-5: Grave No. 5:**
It was in the north and eastern part of the cemetery. An old adult female with Europo-Mongoloid characteristics was laid to rest in the grave. The only known grave goods were two hoops around the head (possibly hair rings).

**HMSZ-43: Grave No. 43:**
Located on the southern edge of the northern group in the cemetery. An old adult male with Mongoloid characteristics was buried in the grave. Patterns of post-holes were recorded in the grave pit, but no grave goods were found during the excavation.

**HMSZ-50: Grave No. 50:**
The grave was in the middle/ western part of the northern group. It is the burial of a young adult male with mongoloid characteristics and without any known grave goods.

**HMSZ-86: Grave No. 86:**
The burial of a middle adult male with Mongoloid characteristics, located at the west side of the northern group in the cemetery. Traces of quiver and arrowheads were found in the grave.

**HMSZ-88: Grave No. 88:**
The grave was in the northern group, at the southern edge of the west side. It is the burial of a middle adult male with Mongoloid characteristics and symbolic trephination on the skull. The grave was without any known grave goods.

**HMSZ-157: Grave No. 157:**

Located in the northern group of the cemetery, at the southern edge of the west side. The burial contained the skeleton of a middle adult male individual with Mongoloid characteristics. Arrowheads and a knife were found in the grave.

**HMSZ-229: Grave No. 229:**
The burial of a young adult female with Europid characteristics, located at the southern part of the cemetery. The grave goods composed of jewelry and clothing ornaments, i.e., penannular hair rings, neckring with hook and eye-terminal, ball button, band bracelet, band ring, and animal-headed bracelet were found in the grave.

**HMSZ-231: Grave No. 231:**
The grave was in the southern part of the cemetery. Patterns of post-holes in the grave pit were recorded during the excavation. An old adult male was buried there without known grave goods.

**HMSZ-245: Grave No. 245:**
It was found in the southern part of the cemetery. In the burial of an old adult female with Mongoloid characteristics, patterns of post-holes were recorded, but no grave goods were found.

**S1.6.5 Ibrány-Esbóhalom (IBE; Szabolcs–Szatmár–Bereg County)** [116,117]
The excavation of the cemetery was carried out between 1985 and 1990 under the leadership of Eszter Istvánovits. The remains of 274 individuals were found in 269 graves. Based on the finds, the 10$^{th}$-century CE part of the cemetery can be well characterized by the low number of burials with weapon- and horse riding-related grave goods and dress fittings (e.g., pendent mounts), as well as simple wire jewelleries (e.g., bracelets) and implements (e.g., knives, fire-lightning equipment). Within the 10$^{th}$-century part, a group with different ethnicities is also distinguished, based on the grave orientation, burial customs, and findings. In addition to the coins related to the reign of the kings of the Árpád Dynasty, the burials of the 11$^{th}$ century CE are characterized by S-terminalled hair rings, beads, and rings. In virtue of the archaeological material, continuity was assumed between the two parts of the site. The cemetery was dated to 940-1075. The state of preservation of the anthropological material is medium, often poor. In anthropological studies 98 men, 82 women, 74 children (inf I-II) and 20 young (juvenile) individuals have been distinguished, thus there is a male surplus for adults. Taxonomic analysis revealed a predominance of Europeans, Mongoloid traits were observed in case of 4 individuals. Craniometric analysis showed a discrepancy between the 10$^{th}$ and 11$^{th}$ century parts, raising the question that the two cemetery parts may hide different populations.
Classification of the cemetery: 10-11/12th-century village cemetery (type 6)in the Kovács system; group B2 in the Gáll system. In our analysis, the samples are categorized as Conqueror and Árpádian age commoner. Nine burials were involved in our analysis:

**IBE-90: Grave 90:**
The burial of an old adult male, without known grave goods. The burial was dated to the 11th century CE.

**IBE-106: Grave 106:**
A middle-old adult female was buried in the grave. Grave goods: an S-terminalled silver hair ring and a silver denar of King Stephan I. The burial was dated to the 11th century CE.

**IBE-107: Grave 107:**
The grave of a middle-old adult male without known grave goods. The burial was dated to the 11th century CE.

**IBE-116: Grave 116:**
The grave of a middle-old adult female with Mongoloid characteristics. Two bronze S-terminalled hair rings were found in the grave. The burial was dated to the 11$^{th}$ century CE.

**IBE-154: Grave 154:**
A young adult male was buried in the grave with the trace of a trepanation on the skull.
A silver penannular hair ring and some iron fragments were in the grave. The burial was dated to the 10$^{th}$ century CE.

**IBE-161: Grave 161:**
The grave of an infant (ca. 10 years old). Traces of wooden coffin, i.e., the iron pants were found. Besides, two penannular hair rings and an obsidian fragment were in the grave. The burial was dated to the 10th century CE.

**IBE-176: Grave 176:**
It was the grave of a middle adult male without known grave goods. The burial was dated to the 10$^{th}$ century CE.

**IBE-206: Grave 206:**
The grave of an infant (ca. 6–8 years old). Grave goods: silver penannular hair ring. two bronze braid ornaments, gilded silver ornaments, neckring, a band bracelet and a wire bracelet. The burial was dated to the 10th century CE.

**S1.6.6 Karos-Eperjesszög I, II, III (K1, K2, K3; Borsod–Abaúj–Zemplén County)** [105,118,119]
The cemeteries were on a hill, oriented in the north-south directions, ca. in a 150-200 m distance from each other. The Karos-Eperjesszög I, II, and III cemeteries are considered as the wealthiest known-group of sites in the 10th-century Carpathian Basin. Characteristics of the cemetery I are the least known since 40-50 graves were destroyed before the excavations. Altogether 13 graves were rescued by Tibor Horváth in 1936. Later, eight graves were found and verified by László Révész in 2001. The saved and stray founds clearly show that the number of graves with horse riding- and weapon-related artefacts was, indeed, high (at least 31 burials with horse riding-related grave founds, four sabers, pieces of archery equipment). The surviving remained belt ornaments, sabretache ornaments, harness ornaments (e.g., rosette-shaped pieces), and Arabian coins (the coins of al-Mustaʿīn billāh from 863/864 CE and the coins of Ismáil ibn Ahmad from 905/906 CE) still indicated the high value of the burials.
Seventy-three graves of the cemetery II were excavated under the leadership of László Révész between 1986 and 1990. However, the cemetery – uniquely in the region- is considered as fully excavated, the total number of the graves assumed to be 84–88. Concerning the different types of jewelry, hair rings (penannular hair rings), earrings with cast-beadrow pendants, beads, the string of beads, braid ornaments, bracelets (e.g., bracelet with coiled terminals, wire bracelet), and finger rings (e.g., penannular plain rings, bezelled rings) were found in the graves. However, the cloth fittings made of precious metal were relatively rare in the cemetery though shoe/boot ornaments can be found in the material. The most characteristic pieces of the objects are, indeed, the so-called prestige grave goods, such as a set of belt ornaments, sabretache ornaments and plates, sabers with precious metal fittings, and ornamented bow cases. The number of burials with horse bones (32 graves) or horse-riding- (37 graves) and weapon-related equipment (22 graves) (e.g., archery equipment, axes, sabers) is very high. Different types of implements (knives, fire-lighting tools, bone tools) and food offerings (pots and animal bones) were also found in a relatively high number of the graves. Altogether, 11 Arabian dirhems (dated between 893 and 923 CE) and 33 Western-European coins (dated between 888 and 924 CE) were in the burials.

The excavation of the cemetery III was led by László Révész between 1988 and 1990. They discovered 19 burials on the site forming a line. The types of grave goods were highly similar to those that were found in cemetery II. However, a set of mounted belts were found only in two cases, and the weapon-related artefacts were composed of archery equipment and two axes. Similarly, jewelry and clothing ornaments, i.e., braid ornaments and shoe/ boot ornaments were found. The number of partial horse burials (7) and graves (5) with horse riding-related equipment is high. Based on the archaeological material, a strong relationship was assumed between the three cemeteries and the populations behind them. According to the results of László Révész, the cemeteries can be dated from the turn of the 9-10th centuries CE to the second third of the 10th century CE.
The anthropological material of the cemeteries was analyzed by Ágnes Kustár. According to the results of the sex and age-at-death determinations, there were no 0–1-year-old infants in the series, assumingly because of the damage caused by the intense and systematic agricultural fieldworks. The ratio of the subadult individuals was 30,39%, while the ratio of the adults was 69,61%. Concerning the adult groups, a significant male surplus was recorded (64,4%/ 35,6%). Based on the results of the taxonomical analysis on 35 skulls, Europid characteristics are dominant in the series (77,1%), but skulls with Europo-Mongoloid morphological patterns are also found in the material (22,9%).
The elite cemetery was classified as a 10th-century quarter cemetery (type 4) in the Kovács system and group A in the Gáll system. In our analysis, the samples are categorized as Conqueror elite.
Three burials from cemetery I (K1), seven burials from cemetery II (K2), and further three burials from cemetery III (K3) were involved in our analysis. All of them have been dated to the 10th century CE.

**K1-1: Grave No. 1:**
It was the grave of a young adult male. The burial contained horse-riding-related equipment (stirrup, girth buckle), antler bow plates, traces of quiver, arrowheads, bronze dress fittings, belt ornaments, and ball buttons.

**K1-10: Grave No. 10:**
A middle adult male was buried in the grave with a knife.

**K1-3286: without grave number**
The burial was destroyed before the excavation and the skeletal remains of a middle adult male were saved.
**K2-16: Grave No. 16:**
Middle-old adult male with Europid (Cromagnoid) characteristics. The grave goods were composed of partial horse burial, horse-riding-related equipment, an axe, archery equipment (i.e., antler bow plates, traces of quiver, antler stiffening plaque of a bow case, arrowheads), and belt mounts.

**K2-18: Grave No. 18:**
The burial of an old adult male. Partial horse burial, horse-riding-related equipment, and a knife were found in the grave.

**K2-26S: Grave No. 26:**
A young adult male was buried with an arrow and a whetstone.

**K2-29: Grave No. 29:**
The burial of a young adult male. The grave contained partial horse burial, horse-riding-related equipment with gilded silver ornaments, food offering (sheep bone), two silver penannular hair rings, band bracelet, a glass bead, a knife, arrowheads, traces of quiver, a set of gilded silver belt ornaments, sabretache plate.

**K2-33: Grave No. 33:**
The juvenile was buried with food offering (sheep bone), a knife, an iron ring (under the mandible), and arrowheads.

**K2-52: Grave No. 52:**
The skeletal remains of a middle adult male with Europid characteristics (Armenoid-Turanian-Pamirian) were in the grave. The richest burial of the cemetery contained: partial horse burial, horse riding related equipment with gilded silver ornaments, gold penannular hair ring, silver wired bracelets, bezelled ring, Arabian dirhems (of Ismail ibn Ahmed /904-905/) and Western-European coins (of Louis IV the Child /899-911/), a set of gilded silver belt ornaments, silver sabretache plate, fire-lighting tools, a knife, saber with gilded mounts and fittings, archery equipment with ornamented bow case, traces of quiver, arrowheads, and antler bow plates.

**K2-61: Grave No. 61:**
An old adult male with Europo-Mongoloid characteristics (Pamirian-Turanian) was buried in the grave. The grave goods were partial horse burial, horse-riding-related equipment, food offering (unidentified animal bone fragments), a bead, antler bow plates, arrowheads, silver band bracelet, fire-lighting tools, a knife, a set of gilded silver belt ornaments, sabretache ornaments.

**K3-6: Grave No. 6:**
The young adult female with Europid characteristics (Atlanto-Mediterranean) was buried in the grave. The burial contained gilded silver belt ornaments or dress fittings, a silver band bracelet, ball buttons, boot mounts, and horse-riding-related equipment with mounts.

**K3-12: Grave No. 12:**
The burial of a middle adult male with Europid characteristics (Cromagnoid-Pamirian) and trepanation on the skull. The grave contained partial horse burial and skull fragments of cattle. In addition, a green metal plaque was recorded on the human bones (e.g., on the radius, ulna, forehead, and teeth).

**K3-13: Grave No. 13:**
The skeletal remains of a middle adult male buried with silver penannular hair ring, two silver plates on the forehead, silver band bracelets, antler bow plate, traces of quiver, arrowheads, a knife, disentangling bone hook, fire-lighting tool, and horse-riding-related equipment with silver ornaments.

**S1.6.7 Kenézlő-Fazekaszug I (KeF1; Borsod–Abaúj–Zemplén County)** [105,120,121]
The burial ground lies on a sandhill called Fazekaszug dűlő, located north of the village. In 1913, after the ploughing of the soil damaged three graves, András Jósa excavated a further 22 burials. In 1919-1920, the landowner uncovered at least eight more graves (e.g., containing partial horse burials and sabretache plates) to the south of the area investigated by Jósa. In the western part of the cemetery, 14 graves (No. 12–25) formed into a line. However, the control excavation performed by László Kovács and László Révész in 1990 revealed that the whole cemetery was not arranged in regular grave rows. The cemetery is considered partially excavated, and the 25 graves account only ca. a third or one-quarter of the burial ground. Eight male burials were equipped with weapons, and only three female burials were identified based on the grave goods. Besides, nine graves did not contain any artefacts.
However, precious metal ornaments were found relatively in a small number, male burials frequently contained weapons and accessories of the belt (sabers, quivers, arrowheads, belt ornamented with mounts, sabretache plates). Additionally, the ratio of partial horse burials in the graves is also high (8).

The dirhams (emitted by Ismail ibn Ahmad between 892–975 CE and 903–904 CE and by Nasr ibn Ahmad Samanid Emir between 916–917 CE) and Western-European denars (e.g., emitted by *rex* Berengar I between 888–915 CE; by Hugh of Provance & Lothar II between 931–945 CE) dated the cemetery to the 2nd third and the middle of the 10th century CE. The anthropological analysis was carried out by András Bíró and Erzsébet Fóthi. The preservation of the bones was low, 4 males and 1 female were described in the population. Besides, the quality of the documentation mostly did not allow to match the archaeological and anthropological data.
The elite cemetery was classified as a 10th-century quarter cemetery (type 4) in the Kovács system and group A in the Gáll system.
One sample was involved in our analysis categorized as Conqueror elite.

**KeF1-10936: Grave No. 9?:**
The easternmost grave in the cemetery did not contain known grave goods. The skeletal material belonged to an adult male individual. The grave was dated to the second third of the 10th century CE.

**S1.6.8 Kenézlő-Fazekaszug II (KeF2; Borsod–Abaúj–Zemplén County)** [105,121]
Another 25 burials were excavated by Nándor Fettich 150 m from the first burial ground assuming, that they are continuous. Thus, the numbering of the graves started with 26. However, the control excavation led by László Kovács and László Révész revealed that the graves belonged to a separate, 2nd cemetery. The graves have formed a single long grave row, and one grave was a double burial.
The special characteristic of the cemetery was the high ratio of partial horse burials (nine male, five female, and one subadult grave). Besides, eight graves contained archery equipment and one a saber. Dress fittings were found only in a few cases. In two female graves, richer inventories have been documented: beads, bracelets, and boot mounts. Two coins were found in the cemetery (emitted by Rudolph of Burgundy between 922–926 CE and by Ismail ibn Ahmad between 907/908–911/912 CE). The anthropological material contained the skeletal remains of twelve males, five females, and one subadult individual.
The elite cemetery was classified as a 10th-century quarter cemetery (type 4) in the Kovács system and group A in the Gáll system.
One sample was involved in our analysis categorized as Conqueror elite.

**KeF2-1045: Grave No. 50 (or 25):**
The grave was in the northernmost part of the cemetery. The skeletal remains of an adult male individual with Mongoloid characteristics were found with partial horse burial, silver penannular hair ring, silver band bracelet, a set of belt mounts, antler bow plates, arrowheads, and horse-riding-related equipment. The burial was dated to the second third of the 10th century CE.

**S1.6.9 (Szeged-) Kiskundorozsma-Hosszúhát (KH; Csongrád–Csanád County)** [122,123]
In 2004, ten 10th-century graves were found during the excavation at the trails of highway M5. The anthropological material was examined by Antónia Marcsik. Five males, two females, and three subadult individuals were described in the series. According to the results of the taxonomical analysis, two individuals showed dominant Europid characteristics (Pamirian) and Europo-Mongoloid characteristics were registered on another group of individuals. One of the male burials contained partial horse burial, horse-riding- (e.g., trapezoid shape stirrups) and weapon-related equipment. The other males, similarly to the subadult individuals, were buried with simpler grave goods such as penannular hair rings, wire bracelets, and food offerings (animal bones) or even without known grave goods. One female was buried with partial horse burial and horse-riding-related equipment, precious metal dress fittings (e.g., pressed ornaments, boot ornaments), jewelry (e.g., braid ornaments, bezelled gold finger ring). The other female burial contained dress fittings, bracelets, and earrings with cast-beadrow pendants. Based on archaeological, radiocarbon and archaeometric data, archaeologists dated

the cemetery, and especially the grave No. 595 to the 2nd third of the 10th century CE (around the period of 940–960).
Classification of the cemetery: 10th-century quarter cemetery (type 4) in the Kovács system; group A in the Gáll system. Two samples were involved in our analysis categorized as Conqueror elite.

**KH-500: Grave No. 500:**
The grave contained the skeletal remains of an adult male with Europo-Mongoloid characteristics, buried with horse-riding-related equipment (e.g., trapezoid shaped stirrups), archery equipment, fire-lighting tool, and a knife. This was the richest among the male burials. The burial was dated to the 2$^{nd}$ half of the 10$^{th}$ century CE.

**KH-596: Grave No. 595:**
The burial of an adult female with Europo-Mongoloid characteristics is the richest grave in the cemetery. The grave goods were partial horse burial, horse-riding-related equipment with gilded silver ornaments, food offerings (pots and sheep bone), gilded silver chaplet (?) and braid ornaments, gold and gilded silver dress fittings, finger ring, silver band bracelets, bronze twisted ankle rings, silver boot ornaments. The burial was dated to the 2$^{nd}$ half of the 10$^{th}$ century CE.

##### S1.6.10 Ladánybene-Benepuszta (LB; Bács–Kiskun County) [99,124]

**LB-1432**
The first-ever known burial in the Carpathian Basin, dated to the 10th century. It was found by shepherds in 1834. The grave was destroyed, and only a part of the grave goods and human bones were brought to the museum. The anthropological analysis was carried out by Pál Lipták, who identified the bones as the skeletal remains of a middle adult male with Europid (Turanid) characteristics and traces of trepanation on the skull. The grave originally contained rich grave goods, i.e., partial horse burial (lost), saber (lost), gilded silver belt ornaments, harness ornaments, unknown metal ornaments, and coins. According to written records, at least 30-40 coins were found, but only 12 of them were given to the museum (emitted by king Berengar I between 888-915 CE and emperor Berengar I between 915-924 CE). The coins were pierced through, and assumingly these were used as dress fittings or harness ornaments. The burial was dated on the 1st half of the 10th century CE. It was categorized as group B2 in the Gáll system and as Conqueror elite in our analysis.

##### S1.6.11 Magyarhomorog-Kónyadomb (MH; Hajdú–Bihar County) [125,126]

The systematic excavation of the cemetery was started by István Dienes between 1961 and 1971 and was finished by László Kovács between 1985 and 1988. The cemetery has a "pagan" segment with 17 graves from the 10th century CE, characterized by a high number of burials with weapon- and horse-riding-related grave goods and a significant male surplus. The larger part consists of 523 "Christian" graves of the 11–12th centuries CE in which the gender rate is balanced. The cemetery is one of the few sites where the burial custom of giving grave goods, which was forbidden by the Christian liturgy, can be observed after the turn of the millennium and the adoption of Christianity (e.g., weapon- and horse-riding-related grave goods in the burials dated with coins related to the reign of the Árpád dynasty kings). Different types of jewelry were the most common archaeological findings: hoops around the head (e.g., S-terminalled hair rings), neck jewelry (e.g., beads), bracelets, and various types of rings. Based on the location of the tombs containing coins, the cemetery started from a central core and expanded evenly toward its edges. During the archaeological analysis, the possibility of discontinuity between the two parts of the cemetery arose. Therefore, archaeologists suggest that the cemetery separately has a 10th-century quarter cemetery (MH1) and an 11–12th-century village cemetery (MH). During the anthropological analysis, eleven men, three women, and three children of unknown sex were identified in the 10th century part. The state of preservation was generally of medium quality. Based on the taxonomic analysis, the Europid groups dominated (about 60%), but,

overall, the proportion of those showing Europo–Mongoloid traits is significant with one individual being classified as Mongoloid type. In paleopathological alterations, developmental abnormalities predominated, which address several further questions about kin relationships within the group. Concerning the 11–12th-century village cemetery, the state of preservation of the skeletons was generally low. A total of 126 males, 174 females, 187 adolescents (infantia I–II), and 36 juvenile individuals were distinguished. According to the craniometric analysis, the dolichocranial skulls were dominant, and the taxonomic analysis revealed that skulls with Europid characteristics were in the highest number, but there were also Europo–Mongoloid characteristics and in six cases Mongoloid types were present (the state of preservation highly limited the classification). Classification of the cemeteries: 10$^{th}$-century quarter cemetery (type 4) and 11-12$^{th}$-century village cemetery (type 7) in the Kovács system; group A and group B2 in the Gáll system.
In our analysis, following the results of the archaeological investigation, the burials from the 10th-century part of the burial ground, i.e., from the quarter cemetery are categorized as Conqueror elite and burials from the 11-12th-century village cemetery are labelled as Árpádian-age commoners. We involved ten samples in our analysis.

**MH1-4: Grave No. 4:**
The burial of a young adult male with Europid characteristics. Shroud fragments, arrowheads, traces of quiver, a girth buckle, a knife, and fire-lighting equipment were found in the grave. The burial was dated to the 10th century CE.

**MH1-9: Grave No. 9:**
A middle adult male with European characteristics (Cromagnoid) was buried in the grave. The grave goods were partial horse burial with a girth buckle, antler bow plates, traces of quiver, and fire-lighting tools. The burial was dated to the 10th century CE.

**MH1-14: Grave No. 14:**
A young adult female with Europid characteristics was found in the grave. The burial contained: silver dress fittings, a string of beads with shells, and a knife. The burial was dated to the 10th century CE.

**MH1-23: Grave No. 23:**
The grave contained the skeletal remains of a middle adult male with Europid (Pamirian) characteristics. Partial horse burial, horse-riding-related equipment, shroud fragments, silver penannular hair ring, silver fragments, antler bow plates, arrowheads, traces of quiver, lyre-shaped belt buckle, fire-lighting tool, and a knife were in the grave. The burial was dated to the 10th century CE.

**MH-88: Grave No. 88:**
It was the grave of a middle adult male without known grave goods. The burial was dated to the 11th century CE.

**MH-106: Grave No. 106:**
It was the burial of an old adult female. The grave goods were: penannular hair rings, glass beads, bracelets, finger rings, food offering (animal bone), and a coin of king Stephen I. The burial was dated to the first half of the 11th century CE.

**MH-107: Grave No. 107:**

A young adult female was buried in the grave. The burial contained gilded silver and silver dress fittings, gilded silver ornaments, a ball button, pendant earring with bead pendant, glass beads, and gilded bronze braid ornaments. The assemblage was dated to the 11th century CE.

**MH-137: Grave No. 137:**
The skeletal remains of an old adult male were found in the grave with arrowheads and a coin emitted by king Stephen I. The burial was dated to the first half of the 11th century CE.

**MH-153: Grave No. 153:**
The burial of a subadult (infantia I) individual. The grave goods were S-terminalled hair rings, neckring, lunula, a string of beads, a bracelet, a finger ring, and a coin emitted by king Stephen I. The burial was dated to the first half of the 11th century CE.

**S1.6.12 Nagykőrös-Fekete dűlő (NK; Pest County)** [127,128]
Two 10$^{th}$-century graves were found in 1950 by construction workers, and unfortunately both burials were destroyed before a possible archaeological investigation. Most of the information are based on the description of the eyewitnesses at the field. The graves contained partial horse burials, horse-riding equipment, weapons (archery equipment and a saber); precious metal jewellery (penannular hair rings), silver dress fitting, gilded silver belt ornaments, and coins emitted by king Berengar I and emperor Berengar I between 888 and 924 CE. The graves were dated to the 10$^{th}$ century CE. Both burials contained the skeletal remains of adult males with Europid characteristics and in one case additional Mongoloid features. The site was classified as group A in the Gáll system. We involved the burial No. 2 in our analysis categorized as Conqueror elite.

**NK-2: Grave No. 2:**
The burial of a middle adult male with Europid (Protoeuropean) characteristics. The grave goods were partial horse burial, horse riding equipment, archery equipment, a saber, gold penannular hair rings, a set of gilded silver belt ornaments, and fragments of a fire-lighting tool.

**S1.6.13 Nagytarcsa-Homokbánya (NTH; Pest County)** [129,130]
In 1967, 21 graves were excavated by László Kovács, and there are data on another seven disturbed graves, but the cemetery can only be considered as partially excavated (estimated at 40–50 graves). The poor archaeological findings consisted of penannular hair rings, twisted neck rings, wire bracelets, rings, and ball buttons. Two burials with weapon-related grave goods (archery equipment, an axe) and four burials with horse-riding-related deposits (e.g., pear-shaped stirrups) were excavated in the cemetery. Based on the findings, the cemetery was dated to the second half of the 10th century CE. The state of preservation of the anthropological material is moderate, often low. The anthropological analysis identified the skeletons of 8 males, 15 females, and four unspecified children. Five skulls belonging to the Europid type proved to be suitable for taxonomic analysis. The site was classified as group B2 in the Gáll system. Four samples were involved in our analysis categorized as Conqueror commoners.

**NTH-1: Grave No. 1:**
An adult male with Europid (Nordoid) characteristics was buried in the grave. The grave goods were a penannular hair ring, horse-riding-related equipment, a knife, an arrowhead, and an axe. The burial was dated to the second half of the 10th century CE.

**NTH-2: Grave No. 2:**
The skeletal remains of an adult female with Mongoloid features were found in the grave with a bracelet and a banded ring. The burial was dated to the second half of the 10th century CE.

**NTH-19: Grave No. 19:**
It was the burial of a middle adult male with Europid (Cromagnoid) characteristics without known grave goods. The grave was dated to the 2nd half of the 10th century CE.

**NTH-20: Grave No. 20:**
The skeletal remains of an adult female with Europid (gracile Mediterranean) characteristics were found in the grave with a penannular hair ring. The burial was dated to the second half of the 10th century CE.

**S1.6.14 Püspökladány-Eperjesvölgy (PLE; Hajdú–Bihar County)** [131,132]
The excavation was carried out under the leadership of Ibolya M. Nepper and Márta Sz. Máthé between 1977 and 1982. A total of 637 graves were found in the cemetery, but due to double burials, the remains of 641 individuals were identified. Based on the findings, the cemetery can be divided into two parts. The western part (about one-third of the cemetery) dating back to the 10th century CE is characterized by burials with weapon- (e.g., archery equipment, sabres, swords) and horse-riding-related (pear-shaped stirrups) grave goods, as well as jewelry, such as hoops around the head (e.g., penannular hair rings), necklaces (e.g., beads), and arm/hand jewelry (e.g., bracelets, rings), dress fittings, and implements (e.g., knives, fire-lighting equipment). The other part was likely used after the statement of the Christian Hungarian Kingdom in the 11th century CE, based on the more common occurrence of coin-dated burials and relatively late grave-good types (e.g., twisted and braided rings, foil beads, S-terminalled hair rings). The anthropological material is of medium or poor preservation. During the anthropological analysis of the cemetery, 191 male, 163 female, and 256 sub-adult (with undetermined sex) individuals were described. According to studies on craniometric and body height data, continuity was assumed between the 10th century and 11th-century parts of the cemetery. Classification of the cemetery: 10-11/12th-century village cemetery (type 6) in the Kovács system; group B2 in the Gáll system. Fourteen samples were involved in our analysis categorized as Conqueror and Árpádian-age commoners.

**PLE-23: Grave No. 23:**
It was the robbed grave of a middle adult male with Europid characteristics. The burial contained a Roman coin from the 4th c. CE, a knife, a whetstone, fire-lighting equipment, a disentangling bone hook, arrowheads, and horse-riding equipment. The grave was at the eastern border of the 10th-century part of the cemetery. The burial was dated to the second part of the 10th century CE.

**PLE-28: Grave No. 28:**
The burial contained the skeleton of a young adult male. The grave goods were a knife, an arrowhead, and horse-riding equipment (bit fragments). The grave was in the north-eastern area in the 10th-century part of the cemetery. It was dated to the second part of the 10th century CE.

**PLE-38: Grave No. 38:**
It was the grave of a young adult male with Europid characteristics. It was at the eastern border of the 10th-century part of the cemetery. A hammer axe, arrowheads, and horse-riding equipment were found in the burial. The assemblage was dated to the second part of the 10th century CE.

**PLE-57: Grave No. 57:**
The skeletal remains of an adult female with Europid characteristics were found in the grave with a penannular hair ring and a band fingerring. The burial was dated to the second half of the 10th century CE, and the grave was in the middle of the 10th-century part of the cemetery.

**PLE-95: Grave No. 95:**

The burial of an infant (ca. 5 years old) was at the borderline between the 10th- and 11th-century parts of the cemetery. The grave contained an iron rattle, glass beads, a cross (pendant), and clothing ornament fragments. It was dated to the first half of the 11th century CE.

**PLE-115: Grave No. 115:**
It was the grave of a middle adult male with Europid characteristics and without known grave goods. The burial was at the borderline between the 10th- and 11th-century parts of the cemetery and it was dated to the first half of the 11th century CE.

**PLE-195: Grave No. 195:**
The skeletal remains of an old adult male were found in the grave with horse-riding equipment, an awl, and food offering (pig humerus and radius). It was at the borderline between the 10th- and 11th-century parts of the cemetery, dated to the second half of the 10th century CE.

**PLE-200: Grave No. 200:**
The burial contained the skeleton of a young adult male. The grave goods were penannular hair rings, a belt buckle, a knife, antler bow plates, traces of quiver, arrowheads, horse-riding equipment (e.g., trapezoid-shaped stirrups), and food offering (sheep/goat bones). The grave was at the north-western corner of the cemetery, and it was dated to the second half of the 10th century CE.

**PLE-216: Grave No. 216:**
The skeleton of a middle adult male with Europid characteristics was in the grave. The burial contained arrowheads, a belt buckle, horse-riding equipment (e.g., saddle mounts and trapezoid-shaped stirrups), and food offering (sheep humerus). The grave was dated to the second half of the 10th century CE, and it was at the western border of the cemetery.

**PLE-327: Grave No. 327:**
It was the burial of a young adult male. Silver S-terminalled hair rings, a twisted silver ring, and an iron fragment were found in the grave. It was at the north-eastern area of the 11th-century part of the cemetery. The burial was dated to the 2nd third – 3rd quarter of the 11th century CE.

**PLE-384: Grave No. 384:**
The skeletal remains of a young adult male with Europid characteristics were found in the grave with a coin emitted by king Solomon (1063–1074). The burial was dated to the last third of the 11th century CE, and it was at the northern border of the cemetery.

**PLE418: Grave No. 418:**
It was the grave of a young adult female with Europid characteristics. The grave goods were silver penannular hair rings, a silver S-terminalled hair ring, a coin emitted by king Bela I (1060–1063), and an iron mount. The burial was dated to the last third of the 11th century CE, and it was at the north-eastern corner of the cemetery.

**PLE-441: Grave No. 441:**
The grave contained the skeleton of an infant (ca. 3 years old) without known grave goods. The burial was dated to the 11th century CE, and it was at the north-eastern corner of the cemetery.

##### S1.6.15 Szeged-Öthalom (1950) (SEO; Csongrád–Csanád County) [105,133,134]

10th-century burials were found at the site in 1879, 1950, and 2009, but they are not considered as one cemetery, rather different groups of graves. We could involve in our study the group that was found in 1950. However, Bálint Alajos rescued thirteen graves, he could not make the proper documentation of the burials because of military restrictions. Seven graves contained known grave goods. In two burials, horse bones, horse-riding equipment, and weapons (archery equipment) were found. Besides, a coin emitted by Berengar I (888-915 CE) was unearthed. The jewelry composed of penannular hair rings, band finger rings, and wire and band bracelets. The graves were dated to the 10th century CE. Ten skeletons were available for the anthropological analysis. All of them were identified as adults, respectively two males and eight females were distinguished. Only three cases were fit for the taxonomical analysis, and all three cases showed Europid (Pamirian and Mediterranean) characteristics. The cemetery was classified as group A in the Gáll system. We involved two samples in our study categorized as Conqueror elite.

**SEO-3: Grave No. 3/1950:**
The skeletal remains of a juvenile with undetermined sex were found in the grave. The burial did not contain any known grave goods.

**SEO-4: Grave No. 4/1950:**
A young adult male with Europid (Pamirian) characteristics was buried in the grave. The grave goods were partial horse burial, a coin emitted by Berengar I (888-915 CE), traces of quiver, antler bow plates, arrowheads, and fragments of an iron girth buckle (?).

**S1.6.16 Sándorfalva-Eperjes (SE; Csongrád–Csanád County)** [111,135]
The site was excavated by Márta Galántha and István Fodor between 1980 and 1981. The cemetery is considered as fully excavated and is composed of 105 graves. However, the archaeological material is only partially published. Assumingly, 10% of the graves were reportedly constructed with a sidewall niche. Food offering (a pot) was found in one case. The archaeological material consisted of weapons (archery equipment) from twelve graves, horse-riding equipment from two graves, gilded silver dress fittings from six graves (rectangular cast ornaments, pressed ornaments, rosette-shaped ornaments, ornaments with pendants), and jewelry (earrings with bead row pendant, finger rings, bracelets, and a lead cross). Only one coin, emitted in the 9th century CE (by Odo of France /887-898/) was found in the cemetery. Based on the archaeological material, the cemetery was dated to the 2nd third of the 10th century CE.
The skeletal remains of 104 individual were available for anthropological analysis. The preservation of the bones is generally poor. Altogether, forty sub-adults (22 infantia I, 13 infantia II, and 5 juvenile) and sixty-four adults (33 young adults, 29 middle adults, and 2 old adults) were described in the series. Twenty-four males and forty-three females were identified in the population. According to data on taxonomical properties, both the male and female skulls showed Europid characteristics. However, the bad state of preservation highly limited the interpretation of the results. Although the cemetery is fully excavated, the population is considered as fragmentary since the data of sex and age-at-death estimation differed from the normal distribution (e.g., lack of fetuses and newborns, significantly low general age-at-death of the adults (25-34 years), and the surplus of females in the cemetery).
Classification of the cemetery: 10th-century village cemetery (type 5) in the Kovács system; group B2 in the Gáll system. We involved four samples in our investigation categorized as Conqueror elite.

**SE-16: Grave No. 16:**
The burial of a middle adult female contained silver earrings with beadrow pendant and silver fragments of a shroud.

**SE-23: Grave No. 23:**

A middle adult male with Europid characteristics (Cromagnoid features) was found in the grave. The traces of quiver, arrowheads, antler bow plates, and horse-riding equipment were registered in the burial.

**SE-64: Grave No. 64:**
It was the grave with sidewall niche of a young adult female with Europid characteristics (Dolichocran). The grave goods were silver earrings with beadrow pendants, rectangular gilded silver dress fittings, and ball buttons.

**SE-114: Grave No. 114:**
The skeletal remains of a young adult male with Europid characteristics (Cromagnoid features) were found in the grave. The burial contained traces of quiver, arrowheads, antler bow plates, and a finger ring.

##### S1.6.17 Sárrétudvari-Hízóföld (SH; Hajdú–Bihar County) [105,131,136]

The site was excavated between 1980 and 1985 by Ibolya M. Nepper. The site, with 262 graves, is considered the largest 10th-century cemetery in Hungary. The cemetery contains a very high proportion of burials with weapon- (archery equipment, sabers, axe) and horse-riding-related (e.g., pear-shaped stirrups, trapezoid-shaped stirrups, bits, girth buckles) grave goods, and the further archaeological findings consist of jewelry, such as hoops around the head (penannular hair rings), neck jewelry (neck rings, beads), and arm jewelry (e.g., bracelets, beads), dress fittings, and implements (e.g., knives, fire-lighting equipment). The cemetery did not compose of grave rows, contained several groups of burials. Based on the composition of the findings and the lack of grave goods dated to the 11th century CE (e.g., coins), the cemetery is dated to the 10th century CE. During the extensive anthropological analysis, 265 individuals were determined, of whom 98 belonged to sub-adult and 162 to adult categories. The skeletal remains are of good/medium preservation. Based on the skulls suitable for taxonomic studies, the series shows European characteristics (e.g., Cromagnoid and Nordoid elements). Classification of the cemetery: 10th-century village cemetery (type 5) in the Kovács system; group B2 in the Gáll system. We involved twelve 10th-century samples in our analysis categorized as Conqueror commoners.

**SH-12: Grave No. 12:**
It was the grave of an adult female buried with silver penannular hair rings, a twisted bracelet, and a banded ring. The burial was at the western border of the cemetery.

**SH-41: Grave No. 41:**
A middle adult male with Europid characteristics was buried in the grave. The burial contained a silver penannular hair ring, traces of quiver, arrowheads, fire-lighting equipment, a knife, and horse-riding equipment (e.g., trapezoid-shaped stirrups with silver ornaments). It was at the southwestern border of the cemetery. The grave was dated to the 2$^{nd}$ half of the 10$^{th}$ century CE.

**SH-66: Grave No. 66:**
It was the grave of a middle adult male. The burial contained a silver penannular hair ring, a knife, arrowheads, a saber, and an antler bow plate. It was in the middle-southern part of the cemetery.

**SH-81: Grave No. 81:**
An old adult male was buried in the grave with partial horse burial, penannular hair rings, food offering (animal bone), a knife, traces of quiver, an arrowhead, and horse-riding equipment (e.g., trapezoid-shaped stirrups). It was in the western part of the cemetery and the burial can be dated to the 2$^{nd}$ half of the 10$^{th}$ century CE.

**SH-98: Grave No. 98:**
A juvenile individual (probably male) with Europid characteristics was buried in the grave. The burial contained fire-lighting equipment, a knife, traces of quiver, arrowheads, and antler bow plate fragments. It was at the western border of the cemetery.

**SH-103: Grave No. 103:**
A middle adult female was buried in the grave. The burial contained gold penannular hair rings, gold clothing ornaments, and horse-riding equipment (e.g., gilded silver and silver saddle ornaments, stirrup with straight foot plate). The richest female grave in the cemetery, located at the southern border of the cemetery.

**SH-106: Grave No. 106:**
The skeletal remains of a middle adult male with Europid characteristics were found in the grave. The grave goods were silver penannular hair rings, a knife, antler bow plates, traces of quiver, and arrowheads. It was in the southern part of the cemetery.

**SH-143: Grave No. 143:**
The skeletal remains of a sub-adult (infantia I) individual were found in the grave with a glass bead, a ball button, gilded silver clothing ornaments, and gilded silver plate ornaments. The grave was at the south-eastern corner of the cemetery.

**SH-175: Grave No. 175:**
It was the grave of a sub-adult individual (infantia II) that contained silver penannular hair rings and an awl. The burial was in the south-eastern part of the cemetery.

**SH-182: Grave No. 182:**
The burial of an old adult male. The grave goods were penannular hair rings, a banded ring, a knife, and arrowheads. It was in the northern part of the cemetery.

**SH-251: Grave No. 251:**
A sub-adult individual (infantia II) was buried in the grave. It contained partial horse burial, a silver penannular hair ring, gilded silver clothing ornaments, bracelets, belt buckle fragments, a knife, fire-lighting equipment, antler bow plates, arrowheads, horse-riding equipment (e.g., trapezoid-shaped stirrup fragments). One of the richest burials in the cemetery, located in the eastern part of the site. The grave was dated to the 2nd half of the 10th century CE.

**S1.6.18 Sárrétudvari-Őrhalom (SO; Hajdú–Bihar County)** [131,137]
The burial site called Őrhalom was located on the surface of a kurgan, near the cemetery of Sárrétudvari-Hízóföld. The field was seriously damaged in 1986 by the intense agricultural works. Five 10th-century graves (and a few prehistorical tombs) were saved during the rescue excavation conducted by Ibolya M Nepper. However, only a small part of the burial ground was investigated, and the initial number of the graves, probably, was around 40-50. Both the archaeological and anthropological data are fragmentary that highly limited the interpretation. The graves formed a North-South oriented row. Different burial customs, i.e., egg (as food offering or symbol of resurrection), partial horse burials in two cases, and traces of a coffin were registered in the cemetery. The fragmentary grave goods consisted of weapon parts (archery equipment and saber), dress fittings (caftan ornament) and belt mounts. The archaeological material was dated to the first half, 2nd third

of the 10th century CE. The anthropological material was analyzed by Luca Kis. The skeletal remains of a sub-adult (2-3 years old) and four adults, respectively two males and two undetermined (showing rather female characteristics) individuals were described. The cemetery was classified as group A in the Gáll system. We involved one sample in our investigation categorized as Conqueror elite.

**SO-5: Grave No. 5:**
An adult male was buried in the grave that contained partial horse burial, traces of a wooden coffin, a caftan ornament, iron fragments, and antler bow plates. The burial was dated to the first two-thirds of the 10th century CE.

**S1.6.19 Sárrétudvari-Poroshalom (SP; Hajdú–Bihar County)** [131,137]
The burial ground was on a small hill (i.e., a tell from the Bronz Age) (Poroshalom) located in a few kilometers from sites Hízóföld and Őrhalom. The investigation was conducted by Ibolya M. Nepper between 1991 and 1993, but it is only partially excavated. However, they found 17 graves, stray finds referred to further burials. Partial horse burials and food offerings were found in two-two cases. The archaeological material composed of weapons (archery equipment and a saber), horse-riding equipment (stirrups, bit, girth buckle) with harness ornaments and inlaid ornaments, dress fittings (belt buckles, silver and gilded rectangular ornaments), jewelry (penannular hair rings, beads, ball buttons, silver bracelets, finger rings), and implements (knife, fire-lighting tool). A set of belt ornaments made of precious metal were found in two graves. The cemetery was dated to the first half/ 2nd third of the 10th century CE. However, radiocarbon data suggests that it was used in the 2$^{nd}$ half of the 10$^{th}$ century CE as well. During the anthropological investigation, fourteen skeletons were available for analysis. Four sub-adult and ten adult individuals, respectively six males and four females were described by Luca Kis. The cemetery was categorized as group A in the Gáll system. We involved three samples in the analysis categorized as Conqueror elite.

**SP-2: Grave No. 2:**
An old adult male was buried in the grave. It is the wealthiest grave known in the cemetery and the region. The parallels are from the burials of the Upper-Tisza region (i.e., in the Karos cemeteries). The grave contained partial horse burial, food offering (sheep humerus), band bracelets, gold and silver plates (probably dress fittings), a knife, a set of gilded silver belt ornaments, antler bow plates, traces of quiver, arrowheads, and horse-riding equipment (stirrups, bit, girth buckle) with harness mounts. The grave was dated to the 2nd third of the 10th century CE.

**SP-9: Grave No. 9:**
The skeletal remains of an old adult female were found in the grave with a neck ring, a string of beads, rectangular dress fittings, ball buttons, and horse-riding equipment. It is only the 9th known female grave with horse-riding equipment in the region (from 110 burials with horse bones or horse-riding equipment). The grave was dated to the 2nd third of the 10th century CE.

**SP-10: Grave No. 10:**
A middle-old adult male was buried in the grave. The burial did not contain any known grave goods. It was dated to the 2$^{nd}$ third of the 10$^{th}$ century CE.

**S1.6.20 Szakony-Kavicsbánya (SZAK; Győr–Moson–Sopron County)** [138–140]
The site was excavated by Gyula Nováki and István Dienes in 1961. The cemetery is considered fully excavated, and seven graves were found. However, the archaeological material of the cemetery is only partially published yet. Burial customs such as partial horse burial and food offerings (animal bones) were registered in three graves. One grave was constructed with a sidewall niche which is more

frequent in Avar age cemeteries. The jewelry found in the graves consisted of hoops around the head (penannular hair rings), earrings (earring with bead row pendant), braid ornaments, beads (e.g., so-called olive beads that are very rare in the 10th-century graves of the Carpathian Basin), bracelets (band bracelet), finger rings (bezelled rings). Dress fittings (rosette-shaped ornament) were found in one grave, and different types of ball buttons were registered in three graves. Weapons (archery equipment) and implements (knife) were only in one grave. Horse-riding equipment was found in the three graves with partial horse burial. Besides the usual elements (stirrups, bit, girth buckle), harness ornaments such as the rosette-shaped ornaments and silver saddle plates were also represented in these burials. The cemetery was dated to the 10$^{th}$ century CE. Kinga Éry described four sub-adults and three adults, respectively she identified a male and two females in the anthropological material. The measurable skulls showed Europid (i.e., Cromagnoid, Pamirian, Turanid) features. Besides, the skeletal remains were analyzed with chemical methods by Imre Lengyel. He determined the sex of the sub-adult individuals (1 boy and three girls) and suggested a family relationship between the No.1 male, No. 7 female, and No. 2 and No. 4 sub-adult individuals. In addition, another family relationship was assumed between the No. 1 male, No. 6 female, and No. 3 and No. 5. sub-adult individuals.
The cemetery was considered as a burial ground for a small family. The burials were classified as group B2 in the Gáll system. We involved four samples in the analysis categorized as Conqueror elite.

**SZAK-1: Grave No. 1:**
A middle adult male with Europid characteristics (Cromagnoid) was buried in the pit. The grave contained partial horse burial, horse-riding equipment (e.g., harness ornaments), food offering (cattle bone), silver band bracelet, gold bezelled finger ring, archery equipment, and a knife.

**SZAK-4: Grave No. 4:**
The skeletal remains of an infant (4-5 years old) with undetermined morphological sex were found in the grave. The burial contained a disc-shaped bronze dress fitting.

**SZAK-6: Grave No. 6:**
It was the burial of a middle adult female with Europid (Turanid) characteristics. The grave goods were food offering (animal bone), partial horse burial, horse-riding equipment (e.g., saddle with silver ornaments, round-shaped stirrups with long loops), a gold penannular hair ring, beads (a string of beads or dress fittings), bone braid ornaments, silver bands (band bracelets or dress fittings), gold bezelled ring, and ball buttons.

**SZAK-7: Grave No. 7:**
The grave with a sidewall niche contained the skeletal remains of a young adult female with Europid (Pamirian and Cromagnoid) characteristics. Partial horse burial, horse-riding equipment (e.g., harness ornaments), an earring with bead row pendant, silver band bracelets, and ball buttons were found in the grave.

**S1.6.21 Szegvár-Szőlőkalja (SZA; Csongrád–Csanád County)** [141]
The 62 graves containing the burials of 63 individuals were excavated in 1979 by Katalin Hegedűs. A burial of opposite orientation (E–W) was found at 30–35 m from the cemetery array. The poor archaeological findings consist of penannular hair rings, beads, wire bracelets, ball buttons, and knives. Horse-riding-related equipment (a fragment of a bridle) and weapons (arrowheads) were unearthed only in one burial. Based on the composition of the archaeological material, the site was dated to the 10th century CE. During the analysis of the anthropological findings, 25 male and 25 female skeletons

were described. Based on the partially published taxonomic studies, the population composition is heterogeneous. The individuals belonged to the Europid, Mongoloid, and Europo–Mongoloid types. The cemetery was described as 10th-century villager cemetery (type 5) in the Kovács system; and group C in the Gáll system. We involved fourteen 10th-century samples in our analysis categorized as Conqueror commoners.

**SZA-7: Grave No. 7:**
The skeletal remains of a young-middle adult male with Europid and Mongoloid characteristics were found in the grave with iron rings and a knife. It was in the middle of the cemetery.

**SZA-20: Grave No. 20:**
The grave contained the skeleton of an adult female with a penannular hair ring. It was in the middle of the cemetery.

**SZA-29: Grave No. 29:**
It was the burial of a young adult female with Europid (brachycran) characteristics, without any known grave goods. The grave was in the middle/ eastern part of the cemetery.

**SZA-44: Grave No. 44:**
The skeleton of a juvenile (with female characteristics) with Europid and slight Mongoloid characteristics was found in the grave. The wealthiest grave in the cemetery contained silver clothing ornaments, a penannular hair ring, a cowry shell, horse-riding equipment (bit fragment), and food offering (fish bones). It was in the northeast corner of the cemetery.

**SZA-52: Grave No. 52:**
It was the grave of an old adult male with Mongoloid characteristics. Silver shroud (?) fragments and iron fragments were documented in the grave pit. It was in the northeast corner of the cemetery.

**SZA-154: Grave No. 154:**
A young adult female with Mongoloid characteristics was buried in the grave. Penannular hair rings, a silver ornament, and iron fragments were found in the grave that was at the northern edge of the cemetery.

**Szegvár-Oromdűlő (SZOD; Csongrád–Csanád County)** [142,143]
The site contained 372 graves which were excavated under the leadership of Gábor Lőrinczy between 1980/1983 and 1996, but many burials (about 75–85) were destroyed due to previous disturbances. Five additional graves were excavated 30–40 m away from the tight array of the cemetery. The archaeological material has consisted of jewelry, such as hoop jewelry around the head (penannular hair rings, coiled hair rings, S-terminalled hair rings, hoops with spiral pendants), neck rings, bracelets, finger rings, and, less frequently, implements (knives, fire-lighting equipment) and dress fittings (one grave). The cemetery was dated to the period between the second third of the 10th century CE and the middle of the 11th century CE. In anthropological studies, skeletons of 110 males, 114 females, and 148 sub-adults of indeterminate sex were described. During the taxonomic analysis, the predominance of Europid type skulls (Cromagnoid and Nordoid) was detected in both the 10th and 11th-century groups, but a small proportion of Mongoloid features were also described. Data on the craniometric properties suggested a population change occurred at the turn of the 10–11th centuries CE.
Based on its size and the composition of the grave goods, the cemetery was classified formerly as 10–11th-century village cemetery (type 6) in the Kovács system; and group C in the Gáll system. We involved four samples in our study categorized as Conqueror and Árpádian-age commoners.

**SZOD-376: Grave No. 376:**
It was the burial of a middle adult male with Europid characteristics. The grave contained a coin emitted by king Andrew I (1046-1060 CE). The burial, located in the northeast part of the cemetery, was dated to the second half of the 11th century CE.

**SZOD-394: Grave No. 394:**
The skeleton of a young adult male was found in the grave without known grave goods. The skull showed Europid (Nordoid) characteristics. The grave was in the southern part of the cemetery, and it was dated to the 10th century CE.

**SZOD-426: Grave No. 426:**
A young adult female with Mongoloid characteristics was buried in the grave. The wealthy burial contained an S-terminalled hair ring, bronze clothing ornaments, a string of beads with gold foil, and a finger-ring. The grave, located in the middle/ eastern part of the cemetery, was dated to the 11th century CE.

**SZOD-566: Grave No. 566:**
It was the burial of a middle adult male with Europo-Mongoloid characteristics. The grave did not contain any known grave goods. It was in the middle/ northern part of the cemetery, and the burial was dated to the 10th century CE.

##### S1.6.23 Tiszanána-Cseh-tanya (TCS; Heves County) [144,145]

Thirty-three skeletons from thirty-two graves were excavated by István Dienes in 1959. However, due to previous earthworks, many graves were destroyed, and the total number of graves was probably around 40–45. Twenty graves contained known grave goods. Burial customs, i.e., partial horse burials in three cases and food offerings (animal bones, a pot) in six cases were registered in the cemetery. Jewelry (penannular hair rings, coiled hair rings, earrings with beadrow pendant, iron neck ring, and band bracelet) and dress fittings (rosette-shaped ornaments, rounded and rectangular ornaments, boot ornaments) were less frequently found in the graves. Weapons (archery equipment) were in three graves. A sabretache plate was found in one grave. The ratio of the horse-riding-related grave goods was high since three burials with horse bones and horse-riding equipment and four more graves with only horse-riding equipment were registered. In two cases, the saddle was ornamented with silver mounts, and in one case, rosette-shaped harness ornaments were found. Coins, emitted in the 9th-10th century CE (by Charles II the Bald /840–875/; by king Berengar I /888–915/; and by Hugh of Provance /926–931/) were only in the burials of sub-adults. The cemetery was dated between the 1st third and the last decades of the 10th century CE. Based on the distribution of the grave goods, the chronological evolution of the burial ground followed a North to South direction.
The qualitative and quantitative preservation of the anthropological remains was frequently low. During the analysis, seventeen sub-adults (infantia I & II), a juvenile, and fourteen adults were distinguished, respectively six males and nine females were identified. The series contains only skulls with Europid characteristics. Most of the cases belonged to the Mediterranean (38,4%) and Brachycran groups (38,15%), followed by the Cromagnoid (15,4%) and Nordoid (7,7%) types.
Based on the number of the graves, the burial customs and grave goods the cemetery was classified as group A in the Gáll system. We involved three samples in our analysis categorized as Conqueror elite.

**TCS-2: Grave No. 2:**

A middle adult female with Europid (Cromagnoid) characteristics was buried in the grave with an infant (7-8 months old) near her right humerus. The grave contained partial horse burial, an earring with a bead-row pendant, beads, a band bracelet, iron belt buckle, boot mounts, silver sheets and horse-riding equipment (e.g., horse bit with sidebars and the metal parts of the saddle) with rosette-shaped gilded bronze harness ornaments. The burial was dated to the 2nd third of the 10th century CE.

**TCS-5: Grave No. 5:**
The skeletal remains of an old adult male with Europid (Nordoid) characteristics were found in the grave with food offering (sheep bone), an arrowhead, traces of quiver, and antler bow plates. It was dated to the 2nd third of the 10th century CE.

**TCS-18: Grave No. 18:**
It was the burial of an old adult male with Europid (Brachycran) characteristics. The grave contained food offerings (sheep or goat bone), and it was dated to the third quarter of the 10th century CE.

#### S1.7 Cemeteries with multiple periods

##### S1.7.1 Vörs-Papkert B (VPB; Somogy County) [146,147]

The multi-period cemetery was excavated between 1983 and 1993 under the leadership of László Költő, Szilvia Honti, and József Szentpéteri. The cemetery as a whole is still unpublished, but it is dated back to the turn of the 8–9th centuries CE to the turn of the 10–11th centuries CE. The 716 excavated burials are mostly from the Late Avar and Carolingian periods, but sporadic burials of 33 people can be dated to the time of the Hungarian Conquest. The study of the cemetery allowed archaeologists to open discussion concerning the supposed survival and continuity of the Transdanubian Late Avar population through the 9–10th centuries CE. Extensive anthropological and serological investigations were carried out on the skeletal remains of the cemetery. Contradictory results were obtained (e.g., concerning the sex determination), and the data is still only partially published.
We involved nine samples from the different periods in our analysis.

**Avar age samples:**
**VPB-31: Grave No. 31:**
The burial of a middle adult female, dated to the 8–9$^{th}$ century CE. The grave goods were a bronze earring with twisted wire ornament and a vessel.

**VPB-307: Grave No. 307/A:**
The grave No. 307 contained the skeletal remains of an adult (307/A) and three sub-adult (307/B, C, and D) individuals. The burial of the young adult male was robbed, and the human and animal bones were mixed. Partial horse burial and iron parts of a coffin were registered in the grave. Besides, gilded bronze belt mounts with scale ornaments, gilded iron harness ornaments (e.g., rosette-shaped bridle ornaments with gilded and silvered bronze inlays), antler bow plates, and three (two whole and fragments of a third) vessels were found in the grave. The burial was dated to the end of the 8$^{th}$ and beginning of the 9$^{th}$ century CE.

**Carolingian Age sample:**
**VPB-279: Grave No. 279:**
The skeletal remains of a juvenile male were found in the grave with a knife, a whetstone, fire-lighting equipment, an iron belt buckle, and a razor. The burial was dated to the 9$^{th}$ century CE.

**Hungarian Conquest period and early Árpádian Age samples:**

**VPB-118: Grave No. 118:**
A middle adult female was buried in the grave with a penannular hair ring and a finger ring. The burial was dated to the 10–11th century CE.

**VPB-167: Grave No. 167:**
It was the burial of a middle adult male. The grave goods were a sword (type H in Petersen's system), a lyre-shaped bronze buckle, ornamented strap-holder, and a knife. The burial was dated to the (1st quarter of the) 10th century CE.

**VPB-310: Grave No. 310:**
A young adult male was buried in the grave. The burial contained an arrowhead, pear-shaped stirrups, fire-lighting equipment, and fragments of an unidentified iron plate. It was dated to the 10th century CE.

**VPB-561: Grave No. 561:**
The grave contained the skeletal remains of an adult male. Traces of quiver with bronze quiver-belt ornaments, arrowheads, a knife, an iron belt buckle, and fire-lighting equipment were found in the burial. It was dated to the 10th century CE.

**VPB-588: Grave No. 588:**
The skeletal remains of an adult with undetermined sex were found in the grave without any known grave goods. Traces of trepanation were registered on the skull. The burial was dated to the 10–11th centuries CE.

**VPB-600: Grave No. 600:**
The burial contained the skeleton of an adult female and a knife. Traces of trepanation were registered on the skull. The grave was dated to the 10–11th centuries CE.

#### S1.8 Neolithic Cemeteries

##### S1.8.1 Alsónyék-Bátaszék, Mérnökségi telep (ANY; Tolna County) [148–150]

An extensive Neolithic settlement and cemetery with ca. 15000 features were unearthed, related to the preventive excavation of the M6 highway between 2006 and 2009. Although the entire material is not published yet, several studies aimed to examine certain parts of the site or specific questions and topics. The settlement and the burial grounds were in use for a long period, and the objects of the Starčevo, Central European Linear Pottery (LBK) and Lengyel Cultures were found. The greatest extent of the site was attained during the period of the Lengyel Culture (dated to the 5th millennium BCE, the Late Neolithic and Early Copper Age) for over 2000 burials and 100 buildings or pits were documented. Among the burials, a special group was identified, characterized by the high number of grave goods and traces of wooden pillars in the grave pit. Therefore, these burials were labelled as 'chief burials'. We involved one sample in our analysis.

**ANY-4027: Grave No. 4027:**
It was the 'elite' burial of a young adult male with severe pathological changes on the skeleton. The grave was framed with timber, and post-holes were at the bottom of the pit. Pottery and stone knife (?) were found in the grave. It belonged to the features of the Lengyel Culture, dated to the 5th millennium BCE.

##### S1.8.2 Hódmezővásárhely-Gorzsa (HGO; Csongrád–Csanád County) [151,152]

A Neolithic tell settlement with six layers was unearthed at Hódmezővásárhely-Gorzsa during multiple excavation periods led by Gyula Gazdapusztai in the 1950's and by Ferenc Horváth in the 1970's, 1980's and 1990's. However, only a low percentage of the site was investigated, and the material is only partially published. Besides the settlement objects, seventy-one burials were found (56 graves and partial remains of a further fifteen individuals from ditches, pits, houses and as stray finds). The graves were oriented to the SE-NW directions, and the deceased were buried in a contracted position laid on their left or right side. Common findings were the pottery (e.g., vessels) deposited near the skull. In addition, bone pins, a stone mace, jewelry (e.g., pendants, bracelets) made of different material (e.g., bone and shell), and several copper artefacts were recovered from the burial ground. The burials were connected to the Tisza Culture and dated to the first half of the 5th millennium BCE. The archaeological dating was confirmed by multiple radiocarbon data derived from the samples of the Gorzsa series (4932–4602 BCE with 95.4% probabilistic). According to the anthropological data, a third of the population was determined as sub-adult, and two-thirds of the adult individuals were described as females. We involved one sample in our study.

**HGO-26: Grave No. 26 (Anthropological ID: HGO-21):**

It was the burial of a young adult female dated to the Tisza Culture period. Pathological changes of infectious origin were detected on the bones.

##### S1.8.3 Vésztő-Mágori-halom (VM; Békés County) [153,154]

Neolithic burials were unearthed in two excavation periods on the Vésztő-Mágori-halom site, one of the largest tell settlements of the Great Hungarian Plain. Between 1972 and 1976, Katalin Hegedűs found 49 graves, one from the Szakálhát group (ca. 6th millennium BCE), thirty from the Tisza Culture (ca. 4900–4500 BCE) and eighteen from the Tiszapolgár Culture (ca. 4400–4000 BCE). János Makkay excavated further three graves from the Late Neolithic period and three burials from the Copper Age in 1986. Concerning the burial customs, the contracted position of the deceased and traces of coffins were frequently registered. The number of the grave goods was generally low, and beads, pottery, ochers (yellow and red), and stone axes were found in the pits. The anthropological analysis focused on the material from the excavations of Katalin Hegedűs. The low quality of the bone material limited the examination of the skeletons and evaluation of the data. Olga Spekker and her colleagues described 12 sub-adults (ten infants and two juveniles) and 18 adults. In addition, they identified five females and ten males in the series. We involved one sample in our study.

**VM-33: Grave No. 33:**

The skeleton of a juvenile with male features was in the grave. Pathological changes of infectious origin were registered on the skeleton. The body was in a contracted position laying on its right side and oriented to the E–SE directions. Traces of coffin and ocher were registered in the pit. The burial belonged to the Tisza Culture, dated to the first half of the 5th millennium BCE.

#### S1.9 Samples from the Caucasus region

##### S1.9.1 Anapa–Andreyevskaya Shhel (Anapa; Krasnodar Krai, Russia) [155]

Excavations were carried out by Andrej Novicsihin and Gabriella M. Lezsák near Anapa, at Andreyevskaya Shhel in two periods, between 1991 and 1992 and in 2019. Eleven graves were unearthed from which two graves were cremation, respectively eight of them were inhumation. Besides, the grave No. 5 was a horse burial. However, in the graves No. 10 and 11 weapons (arrowheads and a saber), while in graves No. 9 and 10 potteries were found, the quality and quantity

of the grave goods is rather poor. Additionally, the inhumation graves uncovered in 2019 did not follow the most important burial custom of the Eurasian nomads, namely laying the dead to rest with a horse or with parts of the horse. Nonetheless, in two graves weapon-related grave goods were found.
The evaluation and publication of the archaeological analysis is in progress. Furthermore, radiocarbon analysis was carried out in the case of graves No. 9, 10 and 11. Based on these results the site can be dated to the 10$^{th}$ (i.e., graves No. 10 and 11), and 11-12$^{th}$ centuries (i.e., grave No. 9) CE. We involved three samples in our analysis.

**Anapa-9: Grave No. 9:**
The skeletal remains of an infant (6–9 months old) were found in the grave pit, orientated from south to north. The burial contained 43 glass beads, 7 copper rattles and hand-made pottery. Based on radiocarbon analysis it was dated to the 11–12$^{th}$ centuries CE (with the highest probability between 1075–1154).

**Anapa-10: Grave No. 10:**
The grave contained the skeletal remains of a sub-adult individual (9-12 years old) oriented along the west-east axis. The grave goods were two copper buttons, six iron arrowheads and pottery. The burial was dated to the 10$^{th}$ century CE (based on the radiocarbon analysis with the highest probability between 886–992).

**Anapa-11: Grave No. 11:**
It was the burial of a juvenile (16–19 years old) individual oriented along from west to east. The grave finds were two copper buttons, fragment of a copper finger ring, a saber, an iron awl, and five arrowheads. It was dated to the 10$^{th}$ century CE (based on the radiocarbon analysis with the highest probability between 880–994).

#### S1.8 Archaeological references:


1. Tejral, J. *Einheimische und Fremde. Das norddanubische Gebiet zur Zeit der Völkerwanderung*. (Archäologisches Institut der Akademie der Wissenschaften der Tschechischen Republik, 2011).
2. Tejral, J. The Problem of the primary Acculturation at the Beginning of the Migration Period. in *Die spätrömische Kaiserzeit und die frühe Völkerwanderungszeit in Mittel- und Osteuropa* (eds. Mączyńska, M. & Grabarczyk, T.) 5–31 (2000).
3. Kazanski, M. Radaigaise et la fin de la civilisation de Černjahov. in *The Pontic-Danubian Realm in the Period of the Great Migration* (eds. Ivanišević, V. & Kazanski, M.) 381–391 (Collége de France CNRS Centre de Recherche D'Histoire et Civlisation de Byzance, 2012).
4. Vida, T. The Huns and the late antique settlement structure in Pannonia. in *Attila's Europe? Structural transformation and strategies of success in the European Hun period. Extended, annotated proceedings of the international conference organised by the Hungarian National Museum and the Eötvös Lorand University, Budapest, June 6-* (eds. Rácz, Z. & Szenthe, G.) 173–200 (Magyar Nemzeti Múzeum-Eötvös Loránd Tudományegyetem, 2021).
5. Mesterházy, K. Eine Gräbergruppe mit nordsüdlicher Grablegung im gepidischen Gräberfeld von Biharkeresztes–Ártánd-Nagyfarkasdomb. *Acta Archaeol. Acad. Sci. Hungariae* **60**, 73–95 (2009).
6. Párducz, M. Archäologische Beitrage zur Geschichte der Hunnenzeit in Ungarn. *Acta Archaeol. Acad. Sci. Hungaricae* **11**, 309–398 (1959).
7. Párducz, M. *Die ethnischen Probleme der Hunnenzeit in Ungarn*. (Akadémiai Kiadó, 1963).
8. Kontny, B. & Mączyńska, M. Ein Kriegergrab aus der frühen Völkerwanderungszeit von

Juszkowo in Nordpolen. in *Dying Gods: religious beliefs in northern and eastern Europe in the time of Christianisation* (eds. Ruhmann, C. & Brieske, V.) 241–261 (Niedersächsischen Landesmuseum Hannover, 2015).
9. Levada, M. To Europe via the Crimea: on possible migration routes of the northern people in the Great Migration period. in *Inter Ambo Maria : Contacts between Scandinavia and the Crimea in the Roman Period: collected papers* (eds. Khrapunov, I. & Stylegar, F.-A.) 115–137 ('Dolia' Publishing House, 2011).
10. Kaczanowski, P. & Radzińska-Nowak, J. The Huns on Polish lands – an attempt to summarise. *A Nyíregyházi Jósa András Múzeum Évkönyve* **55**, 431–450 (2013).
11. Quast, D. Die nördliche Grenzzone des Oströmischen Reiches und Skandinavien im 5. und 6. Jahrhundert. in *Kollaps – Neuordnung – Kontinuität. Gepiden nach dem Untergang des Hunnenreiches* (eds. Vida, T., Quast, D., Rácz, Z. & Koncz, I.) 333–368 (Archaeolingua, 2019).
12. Heather, P. *Fall of Roman Empire. A new history of Rome and the Barbarians*. (Oxford University Press, 2005).
13. Stickler, T. *Aëtius. Gestaltungsspielräume eines Heermeisters im ausgehenden Weströmischen Reich.* (2002).
14. Kim, H. J. *The Huns*. (Routledge, 2016).
15. Blockley, R. C. *The Fragmentary Classicising Historians of the Later Roman Empire II: Eunapius, Olympiodorus, Priscus and Malchus*. (Francis Cairns, 1981).
16. Knipper, C. *et al.* Coalescing traditions—Coalescing people: Community formation in Pannonia after the decline of the Roman Empire. *PLoS One* **15**, e0231760 (2020).
17. Tejral, J. Das Hunnenreich und die Identitatsfragen der barbarischen 'gentes' im Mitteldonauraum aus der Sicht der Archäologie. in *Barbaren im Wandel. Beiträge zur Kultur- und Identitätsbildung in der Völkerwanderungszeit* (ed. Tejral, J.) 55–119 (Archäologisches Institut der Akademie der Wissenschaften der Tschechischen Republik, 2008).
18. Rácz, Z. Zwischen Hunnen- und Gepidenzeit. Frauengräber aus dem 5. Jahrhundert im Karpatenbecken. *Acta Archaeol. Acad. Sci. Hungaricae* **47**, 301–359 (2016).
19. Rácz, Z. Who were the Gepids and Ostrogoths on the middle Danube in the 5th century? An archaeological perspective. in *The Migration Period between the Oder and the Vistula. Vol. 1–2* (eds. Bursche, A., Hines, J. & Zapolska, A.) 771–789 (Brill, 2020).
20. Tomka, P. Az árpási 5. századi sír (The Grave of Árpás from the 5th century). *Arrabona* **39**, 161–188 (2001).
21. Nagy, M. A Hun-Age Burial with Male Skeleton and Horse Bones Found in Budapest. in *Neglected Barbarians* (ed. Curta, F.) 137–175 (Brepols, 2010).
22. Ny. Kovacsóczy, B., Rácz, Z., Mozgai, V., Marcsik, A. & Bajnóczi, B. Archaeological and natural scientific studies on the Hun-period grave from Kecskemét-Mindszenti-dűlő. in *Attila's Europe? Structural Transformation and Strategies of Success in the European Hun Period. Extended, annotated proceedings of the international conference organised by the Hungarian National Museum and the Eötvös Loránd University, Budapest, June 6–* (eds. Rácz, Z. & Szenthe, G.) 305–326 (Magyar Nemzeti Múzeum-Eötvös Loránd Tudományegyetem, 2021).
23. Dobos, A., Gál, S. S., Kelemen, I. & Neparáczki, E. 5th-century burials from Sângeorgiu de Mureș-Kerek-domb (Mureș County, Romania). in *Attila's Europe? Structural transformation and strategies of success in the European Hun period. Extended, annotated proceedings of the international conference organised by the Hungarian National Museum and the Eötvös Lorand University, Budapest, June 6-* (eds. Rácz, Z. & Szenthe, G.) 327–360 (Magyar Nemzeti Múzeum-Eötvös Loránd Tudományegyetem, 2021).
24. Vörös, G. Hunkori szarmata temető Sándorfalva-Eperjesen. *A Móra Ferenc Múzeum Évkönyve* **1982–1983**, 129–172 (1985).

25. Gulyás, B., Rácz, Z., Bajnok, K. & Gait, J. A solitary 5th century burial from Szilvásvárad-Lovaspálya, North-East Hungary. in *Kollaps – Neuordnung – Kontinuität. Gepiden nach dem Untergang des Hunnenreiches* (eds. Vida, T., Quast, D., Rácz, Z. & Koncz, I.) 431–458 (Archaeolingua, 2019).
26. Barfield, T. J. The shadow empires: imperial state formation along the Chinese–Nomad frontier. in *Empires. Perspectives from Archaeology and History* (eds. Alcock, S. E., D'Altroy, T. N., Morrison, K. D. & Sinopoli, C. M.) 10–41 (Cambridge University Press, 2001).
27. Szenthe, G. The 'Late Avar Reform' and the 'Long Eighth Century': A Tale of the Hesitation Between Structural Transformation and the Persistent Nomadic Traditions (7th to 9th Century Ad). *Acta Archaeol. Acad. Sci. Hungaricae* **70**, 215–250 (2019).
28. Daim, F. Avars and Avar Archaeology. An Introduction. in *Regna and Gentes. The Relationship between Late Antique and Early Medieval Peoples and Kingdoms in the Transformation of the Roman World* (eds. Goetz, H.-W., Jarnut, J. & Pohl, W.) 463–570 (Brill, 2003).
29. Haldon, J. F. Commerce and Exchange in the Seventh and Eighth Centuries. Regional Trade and the Movement of Goods. in *Trade and Markets in Byzantium* (ed. Morrison, C.) 99–124 (Dumbarton Oaks, 2012).
30. *The Long Eighth Century. Production, Distribution and Demand*. (Brill, 2000).
31. Pohl, W. *The Avars. A Steppe Empire in Central Europe, 567–822*. (Cornell University Press, 2018).
32. Daim, F. Vom Umgang mit toten Awaren. Bestattungsgebräuche im historischen Kontext. in *Erinnerungskultur im Bestattungsritual. Archäologisch-Historisches Forum* (eds. Jarnut, J. & Wemhof, M.) 41–60 (Wilhelm Fink Verlag, 2003).
33. Csiky, G. *Avar-Age Polearms and Edged Weapons. Classification, Typology, Chronology and Technology*. (Brill, 2015).
34. Szenthe, G. Crisis or Innovation? A Technology-Inspired Narrative of Social Dinamics in the Carpathian Basin during the Eighth Century. in *Between Byzantium and the Steppe. Archaeological and Historical Studies in Honour of Csanád Bálint on the Occasion of His 70th Birthday* (eds. Bollók, Á., Csiky, G. & Vida, T.) 351–370 (Institute of Archaeology, Research Centre for the Humanities, Hungarian Academy of Sciences, 2016).
35. *Archäologische Denkmäler der Awarenzeit im Mitteldonaubecken*. (Institute of Archaeology, Research Centre for the Humanities, Hungarian Academy of Sciences, 2002).
36. Kiss P., A. *"…ut strenui viri…" A gepidák Kárpát-medencei története. The History of the Gepids in the Carpathian Basin*. (Szegedi Középkorász Műhely-Szegedi Tudományegyetem Történelemtudományi Doktori Iskola, 2015).
37. Koncz, I. A Historical Date and its Archaeological Consequences. *Acta Archaeol. Acad. Sci. Hungaricae* **66**, 315–340 (2015).
38. Vida, T. Local or foreign Romans? The problem of the late antique population of the 6th-7th centuries in Pannonia. in *Foreigners in Early Medieval Europe. Thirteen international studies on early medieval mobility* (ed. Quast, D.) 233–259 (Römisch-Germanisches Zentralmuseum, 2009).
39. Vida, T. Conflict and coexistence. The local population of the Carpathian Basin under Avar rule (sixth to seventh century). in *The Other Europe in the Middle Ages. Avars, Bulgars, Khazars and Cumans* (ed. Curta, F.) 13–46 (Brill, 2008).
40. Kovrig, I. *Das awarenzeitliche Gräberfeld von Alattyán*. (Akadémiai Kiadó, 1963).
41. Juhász, I. *Das awarenzeitliche Gräberfeld in Szarvas-Grexa-Téglagyár, FO 68*. (2004).
42. Preiser-Kapeller, J. The climate of the khagan. Observations on palaeoenvironmental factors in the history of the Avars (6th–9th century). in *Lebenswelten zwischen Archäologie und Geschichte. Festschrift für Falko Daim zu seinem 65. Geburtstag* (eds. Drauschke, J. et al.)

311–324 (Römisch Germanisches Zentralmuseum, 2018).
43. Hakenbeck, S. Migration in archaeology: Are we nearly there yet? *Archaeol. Rev. from Cambridge* **23**, 9–26 (2008).
44. Kővári, I. & Szathmáry, L. A továbbélés megítélése az Ároktő, Csík-gát lelőhelyen feltárt 5–9. századi csontvázleletek alapján. *A Herman Ottó Múzeum Évkönyve* **42**, 135–164 (2003).
45. Wenger, S. Données osteomatriques sur le matériel anthropologique du cimetière d'Alattyán-Tulát, provenant de l'époque avare. *Crania Hungarica* **2**, 1–55 (1957).
46. Lipták, P. Historisch-antropologische Auswertung der im Awarenzeitlichen Gräberfeld von Alattyán erschlossenen Skelettreste. in *Das awarenzeitliche Gräberfeld von Alattyán* 245–258 (Akadémiai Kiadó, 1963).
47. Cseh, G. & Varga, S. Avar kori temetkezések Apátfalva-Nagyút-dűlőn (M43 43. Lh.) – Burials from the Avar Age at the site Apátfalva-Nagyút-dűlő (Site M43 43.). in *Úton a kultúrák földjén. Az M43-as autópálya Szeged–országhatár közötti szakasz régészeti feltárásai és a hozzá kapcsolódó vizsgálatok* (eds. T. Gábor, S. & Czukor, P.) 443–477 (Móra Ferenc Múzeum, 2017).
48. Marcsik, A. & Varga, S. Az Apátfalva-Nagyút‐ dűlőn feltárt avar kori temetkezések antropológiai adatainak összefoglalása. in *Úton a kultúrák földjén. Az M43-as autópálya Szeged–országhatár közötti szakasz régészeti feltárásai és a hozzá kapcsolódó vizsgálatok* (eds. T. Gábor, S. & Czukor, P.) 479–487 (Móra Ferenc Múzeum, 2017).
49. Gáll, E. & Szenthe, G. The problem of "structural integration". A case study of the 9th–10th century burials (Graves 49 and 50) at Hortobágy – Árkus. *Mater. şi Cercet. Arheol.* **16**, 181–197 (2020).
50. Garam, É. *Katalog der awarenzeitlichen Goldgegenstände und der Fundstücke aus den Fürstengräbern im Ungarischen Nationalmuseum*. (Magyar Nemzeti Múzeum, 1993).
51. Nagy, M. *Awarenzeitliche Gräberfelder im Stadtgebeit von Budapest*. (Magyar Tudományos Akadémia Régészeti Intézete, 1998).
52. Balogh, C. Kora avar kori ún. propeller alakú övveret a kunpeszéri 3. sírból. — Frühawarischer sog. propellerförmiger Gürtelbeschlag aus Grab 3 in Kunpeszér. *A Móra Ferenc Múzeum Évkönyve – Stud. Archaeol.* **12**, 257–276 (2011).
53. Wicker, E. Csólyospálos – Felsőpálos, Budai-tanya. in *Archäologische Denkmäler der Awarenzeit in Mitteleuropa* (ed. Szentpéteri, J.) 100 (Magyar Tudományos Akadémia Régészeti Intézete, 2002).
54. Marcsik, A., Balázs, J. & Molnár, E. Zománc hypoplasia megjelenése és kronológiai eloszlása egy avar kori széria embertani leletein. *Folia Anthropol.* **10**, 93–98 (2011).
55. Lantos, A. Kora avar sírok Dunavecséről – Early Avar burials from Dunavecse. in *Hatalmi központok az Avar Kaganátusban – Power centres of the Avar Khaganate* (eds. Balogh, C., Szentpéteri, J. & Wicker, E.) 97–114 (Kecskeméti Katona József Múzeum, 2019).
56. Balogh, C. & Kőhegyi, M. Fajsz környéki avar kori temetők II. Kora avar kori sírok Fajsz-Garadombon. — Awarenzeitliche Gräberfelder in der Umgebung von Fajsz. Frühawarenzeitliche Gräber von Fajsz-Garadomb. *A Móra Ferenc Múzeum Évkönyve – Stud. Archaeol.* **10**, 333–363 (2001).
57. Balogh, C. A Felgyő, Ürmös-tanyai avar kori temető – The Avar cemetery at Felgyő, Ürmös-tanya. in *Felgyő, Ürmös-tanya. Bronz kori és avar kori leletek László Gyula felgyői ásatásának anyagában* (eds. Balogh, C. & P. Fischl, K.) 185–381 (Móra Ferenc Múzeum, 2010).
58. Marcsik, A. Felgyő, Ürmös-tanya avar kori temető humán csontvázmaradványai - The Human Skeletal Remains from the Avar Cemetery at Felgyő. in *Felgyő, Ürmös-tanya. Bronz kori és avar kori leletek László Gyula felgyői ásatásának anyagában* (eds. Balogh, C. & P. Fischl, K.) 383–391 (Móra Ferenc Múzeum, 2010).
59. Balogh, C. Hajós-Cifrahegy – Avar cemetery at Hajós-Cifrahegy. in *Avar kori temetők Bács-*

*Kiskun megyében 1 – Avar-age cemeteries in Bács-Kiskun County 1* 37–124 (Kecskeméti Katona József Múzeum, 2018).
60. Marcsik, A., Balázs, J. & Molnár, E. Anthropological analysis of an Avar age cemetery from the Duna-Tisza interfluve (Hajós-Cifrahegy). in *The Talking Dead. New Results from Central and Eastern European Osteoarchaeology* (ed. Gál, S. S.) 65–78 (Mega Publishing House, 2016).
61. Lipták, P. Homokmégy-Halom avarkori népessége. *Anthropol. Kozl.* **4**, 25–42 (1956).
62. László, G. Jegyzetek a Homokmégy-halomi késő avar temetőhöz (Függelék). *Anthropol. Kozl.* **4**, 43–46 (1956).
63. Wenger, S. L 'anthropologie du cimetière de Jánoshida-Tótkérpuszta. *Ann. Hist. Musei Nat. Hungarici* **4**, 231–244 (1952).
64. Erdélyi, I. A jánoshidai avarkori temető. *Régészeti Füzetek* **2**, (1958).
65. M. Nepper, I. A Kaba-bitozugi avar temető. *Commun. Archaeol. Hungariae* 93–123 (1982).
66. Marcsik, A. *et al.* Adatok a lepra, tuberculosis és syphilis magyarországi paleopatológiájához. *Folia Anthropol.* **8**, 5–34 (2009).
67. Samu, A. & Szalontai, C. Az avar kori sírrablásokról három kiskundorozsmai temető kapcsán. in *Hadak útján XXIV. A népvándorláskor fiatal kutatóinak XXIV. konferenciája Esztergom, 2014. november 4-6.* (eds. Türk, A., Balogh, C. & Major, B.) 763–811 (Archaeolingua, 2017).
68. H. Tóth, E. Korai avar vezetőréteg családi temetője a kunbábonyi kagán szállásterületén. *Múzeumi Kut. Bács-Kiskun Megyében* 10–20 (1984).
69. Marcsik, A. A kunpeszéri avar kori széria humán csontanyagának feldolgozása (Felsőpeszéri út-Homokbánya) – Human bone material of the Avarian Age series from Kunpeszér (Felsőpeszéri út-Homokbánya). in *„In terra quondam Avarorum…" Ünnepi tanulmányok H. Tóth Elvira 80. születésnapjára* (eds. Somogyvári, Á. & V. Székely, G.) 175–190 (Bács-Kiskun Megyei Önkormányzat Katona József Múzeuma, 2009).
70. Balogh, C. A Duna–Tisza köze avar kori betelepülésének problémái – Problems of colonization in the territory between Danube and Tisza Rivers in the Avar Age. (Eötvös Loránd Tudományegyetem, 2013).
71. Török, G. The Kiskőrös Pohibuj-Mackó-dűlő Cemetery. in *Avar Finds in the Hungarian National Museum. Cemeteries of the Avar Periods (567–829) in Hungary I* (ed. Kovrig, I.) 283–304 (Akadémiai Kiadó, 1975).
72. Lipták, P. Contributions à l'anthropologie des temps avars de la région de Kiskőrös. *Crania Hungarica* **1**, 47–52 (1956).
73. László, G. *Études archéologiques sur l'historie de la société des Avars*. (Akadémiai Kiadó, 1955).
74. Nemeskéri, J. Étude anthropologique des squelettes du clan princier avar découverts au cimetiére de Kiskőrös-Vágóhíd. in *Étude arcltéologiques sur l'histoire de la société des Avars* 189–210 (Akadémiai Kiadó, 1955).
75. Balogh, C. Egy Maros-völgyi kora avar kori közösség életmódjáról a régészeti adatok alapján – Makó–Mikócsa-halom. in *Környezettörténet. Tanulmányok Sümegi Pál professzor 60 éves születésnapi köszöntésére* (eds. Törőcsik, T., Gulyás, S., Molnár, D. & Náfrádi, K.) 129–142 (Szegedi Tudományegyetem Földrajzi és Földtudományi Intézet, 2020).
76. Balogh, C. A Byzantine Gold Cross in an Avar Period Grave from Southeastern Hungary. in *Lebenswelten zwischen Archäologie und Geschichte. Festschrift für Falko Daim zu seinem 65. Geburtstag* (eds. Drauschke, J. et al.) 25–42 (Römisch-Germanisches Zentralmuseum, 2018).
77. Balogh, C. Mélykút–Sánc-dűlő – Avar-age cemetery at Mélykút-Sánc-dűlő. in *Avar kori temetők Bács-Kiskun megyében 1 – Avar-age cemeteries in Bács-Kiskun County 1* 135–166 (Kecskeméti Katona József Múzeum, 2018).
78. Marcsik, A. A mélykúti avarkori temető embertani leleteinek vizsgálata. *Anthropol. Kozl.* **15**, 87–95 (1971).

79. Rácz, Z. A madaras-téglavetői avar temető (Kőhegyi Mihály ásatása 1959–62) – Das awarische Gräberfeld von Madaras-Téglavető (Ausgrabungen von Mihály Kőhegyi 1959–62). *A Móra Ferenc Múzeum Évkönyve – Stud. Archaeol.* **5**, 347–395 (1999).
80. Lipták, P. & Marcsik, A. A Madaras-Téglavető avar temető csontvázmaradványainak embertani jellemzése. *Cumania* **4**, 115–140 (1976).
81. Juhász, I. *Awarenzeitliche Gräberfelder in der Gemarkung von Orosháza*. (1995).
82. Bende, L. *Temetkezési szokások a Körös–Tisza–Maros közén az avar kor második felében*. (Pázmány Péter Katolikus Egyetem Bölcsészet és Társadalomtudományi Kar Régészeti Intézet, 2017).
83. Molnár, E. Pitvaros-Víztározó avar kori temető emebertani vizsgálatának eredményei. in *Temetkezési szokások a Körös–Tisza–Maros közén az avar kor második felében* 505–532 (Pázmány Péter Katolikus Egyetem Bölcsészet és Társadalomtudományi Kar Régészeti Intézet, 2017).
84. Kőhegyi, M. Sükösd-Ságod. in *Archäologische Denkmäler der Awarenzeit in Mitteleuropa* (ed. Szentpéteri, J.) 332–333 (Magyar Tudományos Akadémia Régészeti Intézete, 2002).
85. Kőhegyi, M. & Marcsik, A. The Avar Age Cemetery at Sükösd. *Acta Antiq. Archaeol.* **14**, 87–94 (1971).
86. Lipták, P. & Vámos, K. A 'Fehértó-A' megnevezésű avar kori temető csontvázanyagának embertani vizsgálata. *Anthropol. Kozl.* **13**, 3–30 (1969).
87. Madaras, L. *The Szeged-Fehértó 'A' and 'B' cemeteries.* (Kossuth Lajos Tudományegyetem Néprajzi Tanszék Ethnica Alapítványa, 1995).
88. Lipták, P. & Marcsik, A. Szeged-Kundomb avar kori népességének embertani vizsgálata. *Anthropol. Kozl.* **10**, 13–56 (1966).
89. Salamon, Á. & Cs. Sebestyén, K. The Szeged-Kundomb Cemetery. in *Das Awarische Corpus. Avar Corpus Füzetek* (eds. Kovrig, I. & Madaras, L.) 8–109 (Kossuth Lajos Tudományegyetem Néprajzi Tanszék Ethnica Alapítványa, 1995).
90. B. Nagy, K. *A székkutas-kápolnadűlői avar temető*. (Móra Ferenc Múzeum, 2003).
91. Marcsik, A. & Szalai, F. Néhány megjegyzés a fülkés sírokban eltemetett egyének embertani arculatáról. *A Móra Ferenc Múzeum Évkönyve – Stud. Archaeol.* **1**, 453–458 (1995).
92. Vámos, K. 'Szeged-Makkoserdő' avar kori népességének embertani vizsgálata. *Anthropol. Kozl.* **17**, 29–39 (1973).
93. Salamon, Á. The Szeged-Makkoserdő cemetery. in *Das Awarische Corpus. Avar Corpus Füzetek* (eds. Kovrig, I. & Madaras, L.) 109–207 (Kossuth Lajos Tudományegyetem Néprajzi Tanszék Ethnica Alapítványa, 1995).
94. Farkas, G., Marcsik, A. & Oláh, S. Történeti idők embere Szegváron. *Anthropol. Kozl.* **35**, 7–37 (1993).
95. Lőrinczy, G. Vorläufiger Bericht über die Freilegung des Gräberfeldes aus dem 6-7. Jh. in Szegvár-Oromdülő. Weitere Daten zur Interpretierung und Bewertung der partiellen Tierbestattungen in der frühen Awarenzeit. *Commun. Archaeol. Hungariae* 81–124 (1992).
96. Lőrinczy, G. Fülkesírok a Szegvár-oromdűlői kora avar kori temetőből. Néhány megjegyzés a fülkesíros temetkezések változatairól, kronológiájáról és területi elhelyezkedéséről. *A Móra Ferenc Múzeum Évkönyve – Stud. Archaeol.* **1**, 399–416 (1995).
97. Molnár, E. & Marcsik, A. Paleopatológiai elváltozások egy avar kori széria (Szarvas 68. lelőhely) embertani anyagában. *A Békés Megyei Múzeumok Közleményei* **24–25**, 411–428 (2003).
98. Garam, É. *Das awarenzeitliche Gräberfeld von Tiszafüred. Cemeteries of the Avar period (567-829) in Hungary vol. 3.* (Akadémiai Kiadó, 1995).
99. Lipták, P. Awaren und Magyaren im Donau-Theiss Zwischenstromgebiet. *Acta Archaeol. Acad. Sci. Hungaricae* **8**, 199–268 (1957).
100. Tóth, S. L. *A magyar törzsszövetség politikai életrajza. A magyarság a 9–10. században*.

(Belvedere Meridionale, 2015).
101. Szőke, B. M. A Kárpát-medence a Karoling-korban és a magyar honfoglalás. in *Magyar őstörténet. Tudomány és hagyományőrzés* (eds. Sudár, B., Szentpéteri, J., Petkes, Z., Lezsák, G. & Zsidai, Z.) 31–42 (MTA Bölcsészettudományi Kutatóközpont, 2014).
102. Révész, L. *The Era of the Hungarian Conquest*. (Hungarian National Museum, 2014).
103. Hampel, J. A honfoglalás kor hazai emlékei. in *A magyar honfoglalás kútfői* (eds. Pauler, G. & Szilágyi, S.) 507–826 (Nap Kiadó, 1900).
104. Langó, P. *Amit elrejt a föld... A 10. századi magyarság anyagi kuktúrájának régészeti kutatása a Kárpát-medencében*. (L'Harmattan, 2007).
105. Révész, L. *A 10–11. századi temetők regionális jellemzői a Keleti-Kárpátoktól a Dunáig*. (Szegedi Tudományegyetem Régészeti Tanszéke–Magyar Tudományos Akadémia Régészeti Intézete–Magyar Nemzeti Múzeum–Martin Opitz Kiadó, 2020).
106. Takács, M. Három nézőpont a honfoglaló magyarokról. *Dolg. az Erdélyi Múzeum Érem- és Régiségtárából* **1**, 67–98 (2006).
107. Gáll, E. & M. Lezsák, G. A view from the West to the East. An analysis of the characteristics, chronology, and distribution of the sabretache plates in the 10th century. *Zeitschrift für Archäologie des Mittelalters* **46**, 85–111 (2018).
108. Gáll, E. *A hatalom forrása és a magyar honfoglalás – hódítás és integráció. A korai magyar történelem egy régész szemszögéből*. (Magyarságkutató Intézet, 2019).
109. Gáll, E. *Az Erdélyi-medence, a Partium és a Bánság 10–11. századi temetői*. (Szegedi Tudományegyetem Régészeti Tanszéke–Magyar Nemzeti Múzeum–Magyar Tudományos Akadémia Régészeti Intézete, 2013).
110. Kürti, B. Az algyői temetőről röviden. Leletek és jelenségek előfordulási megoszlása a temető területén. *Múzeumi Kut. Csongrád Megyében* 15–36 (1998).
111. Marcsik, A., Just, Z. & Szalai, F. Honfoglalás kori csontmaradványok a Duna-Tisza köze déli területéről (Szeged-Algyő, Sándorfalva-Eperjes). in *Régészeti és Természettudományi Adatok a Maros-torkolat nyugati oldalának 10. századi történetéhez.* (eds. Türk, A., Lőrinczy, G. & Marcsik, A.) 377–418 (Archaeolingua, 2015).
112. Füredi, Á. Honfoglalás kori tarsolylemez Pest megyében. A Bugyi-felsőványi 2. sír. *Archaeol. Értesítő* **137**, 207–234 (2012).
113. Kürti, B. Honfoglaló magyar sírok Szeged-Csongrádi úton. in *Honfoglaló magyarság Árpád-kori magyarság. Antropológia-Régészet-Történelem* (eds. Pálfi, G., Farkas, G. & Molnár, E.) 59–64 (JATE Press, 1996).
114. Gallina, Z. & Varga, S. *A Duna-Tisza közének honfoglalás és kora Árpád-kori temetői, sír- és kincsleletei I. A Kalocsai Sárköz a 10–11. században*. (Szegedi Tudományegyetem Régészeti Tanszéke–Magyar Tudományos Akadémia Régészeti Intézete–Magyar Nemzeti Múzeum–Viski Károly Múzeum, 2016).
115. Marcsik, A., Bereczki, Z. & Molnár, E. Homokmégy-Székes 10-11. századi temető csontvázanyagának vizsgálata. in *A Duna-Tisza közének honfoglalás és kora Árpád-kori temetői, sír- és kincsleletei I. A Kalocsai Sárköz a 10–11. században* 179–207 (Szegedi Tudományegyetem Régészeti Tanszéke–Magyar Tudományos Akadémia Régészeti Intézete–Magyar Nemzeti Múzeum–Viski Károly Múzeum, 2016).
116. Istvánovits, E. *A Rétköz honfoglalás és Árpád-kori emlékanyaga*. (Jósa András Múzeum–Magyar Nemzeti Múzeum–Magyar Tudományos Akadémia Régészeti Intézete, 2003).
117. Szathmáry, L. Az Ibrány-Esbó-halom X–XI. századi temetőjének csontvázleletein végzett vizsgálatok eredményeinek összefoglalása. in *A Rétköz honfoglalás és Árpád-kori emlékanyaga* 385–391 (Jósa András Múzeum–Magyar Nemzeti Múzeum–Magyar Tudományos Akadémia Régészeti Intézete, 2003).
118. Révész, L. *A karosi honfoglalás kori temetők. Régészeti adatok a Felső-Tisza-vidék X. századi történetéhez*. (Herman Ottó Múzeum–Magyar Nemzeti Múzeum, 1996).

119. Kustár, Á. A Karos I-II-III. honfoglalás kori temetők embertani vizsgálata. in *A karosi honfoglalás kori temetők. Régészeti adatok a Felső-Tisza-vidék X. századi történetéhez* 395–456 (Herman Ottó Múzeum–Magyar Nemzeti Múzeum, 1996).
120. Jósa, A. Honfoglalás kori emlékek Szabolcsban. *Archaeol. Értesítő* **34**, 69–184; 303–340 (1914).
121. Bíró, A. & Fóthi, E. A Kenézlő-Fazekaszug I-II. honfoglalás kori temetők embertani vizsgálata. in *IV. Kárpát-medencei Biológia Szimpózium* (ed. Korsós, Z.) 51–56 (Magyar Biológiai Társaság, 2005).
122. Türk, A. & Lőrinczy, G. Régészeti adatok és természettudományi eredmények a Maros-torkolat nyugati oldalának 10. századi történetéhez. in *Régészeti és Természettudományi Adatok a Maros-torkolat nyugati oldalának 10. századi történetéhez* (eds. Türk, A., Lőrinczy, G. & Marcsik, A.) 15–23 (Archaeolingua, 2015).
123. Marcsik, A. Szeged-Kiskundorozsma, Hosszúhát és Kistelek M5 57. (27/71) lelőhelyen feltárt humán csontvázanyag. *A Móra Ferenc Múzeum Évkönyve– Stud. Archaeol.* **12**, 493–504 (2011).
124. Hampel, J. *Alterthümer des frühen Mittelalters in Ungarn I–III.* (1905).
125. Kovács, L. *Magyarhomorog–Kónya-domb 10. századi szállási és 11–12. századi falusi temetője*. (Szegedi Tudományegyetem Régészeti Tanszéke–Magyar Nemzeti Múzeum–Magyar Tudományos Akadémia Régészeti Intézete, 2019).
126. Marcsik, A., Balázs, J., Hajdu, T., Molnár, E. & Szeniczey, T. A Magyarhomorog-Kónya-dombi 10. és 11-12. századi temető embertani anyaga. in *Magyarhomorog–Kónya-domb 10. századi szállási és 11–12. századi falusi temetője* 557–610 (Szegedi Tudományegyetem Régészeti Tanszéke–Magyar Nemzeti Múzeum–Magyar Tudományos Akadémia Régészeti Intézete, 2019).
127. Dienes, I. Honfoglaló magyarok sírjai Nagykőrösön. *Archaeol. Értesítő* **87**, 177–180 (1960).
128. Lipták, P. New Hungarian Skeletal Remains of the 10th Century from the Danube-Tisza Plain. *Ann. Hist. Musei Nat. Hungarici* **3**, 177–287 (1952).
129. Kovács, L. Honfoglalás kori sírok Nagytarcsán II: A homokbányai temetőrészlet. Adatok a nyéltámaszos balták, valamint a trapéz alakú kengyelek értékeléséhez. *ommunicationes Archaeol. Hungariae* 93–121 (1986).
130. Lotterhof, E. he Anthropological Investigation of the Tenth Century Population Excavated at Nagytarcsa. *Anthropol. Hungarica* **12**, 41–62 (1973).
131. M. Nepper, I. *Hajdú-Bihar megye 10–11. századi sírleletei I–II.* (Déri Múzeum–Magyar Nemzeti Múzeum–Magyar Tudományos Akadémia Régészeti Intézete, 2002).
132. Lenkey, Z. *et al.* Tizenöt 8-13. századi népesség kraniológiai elemzése. in *Árpád előtt- Árpád után. Antropológiai vizsgálatok az Alföld I-XIII. századi csontvázleletein* (ed. Szathmáry, L.) 27–40 (JATE Press, 2008).
133. Bálint, C. Honfoglalás kori sírok Szeged-Öthalmon. *A Móra Ferenc Múzeum Évkönyve* 47–89 (1968).
134. Marcsik, A. Embertani adatok a Maros-torokkal szembeni mikrorégió 10. századi történetéhez. in *Régészeti és Természettudományi Adatok a Maros-torkolat nyugati oldalának 10. századi történetéhez* (eds. Türk, A., Lőrinczy, G. & Marcsik, A.) 433–464 (Archaeolingua, 2015).
135. Fodor, I. Honfoglaláskori temető Sándorfalván (Előzetes közlemény). *Acta Antiq. Archaeol.* **5**, 17–33 (1985).
136. Oláh, S. Sárrétudvari–Hízóföld honfoglalás kori temetőjének történeti embertani értékelése. (József Attila Tudományegyetem, Szeged, 1990).
137. Kis, L. Bioarchaeológiai adatok Sárrétudvari-Őrhalom és Sárrétudvari-Poroshalom 10. századi lelőhelyek társadalomrégészeti megítéléséhez. (University of Szeged, Szeged, 2018).
138. Dienes, I. *A honfoglaló magyarok*. (Hereditas Corvina Kiadó, 1972).

139. Éry, K. Honfoglaló magyar csontvázleletek Szakonyról. *Arrabona* **19–20**, 177–182 (1978).
140. Horváth, C. Szakony-Kavicsbánya cemetery from the age of the Hungarian conquest. *Ephemer. Hungarol.* **1**, 289–314 (2021).
141. Lőrinczy, G. Szegvár–Szőlőkalja X. századi temetője. *Commun. Archaeol. Hungariae* 141–162 (1985).
142. Bende, L. & Lőrinczy, G. A Szegvár-Oromdűlői 10-11. századi temető. *A Móra Ferenc Múzeum Évkönyve– Stud. Archaeol.* **3**, 201–285 (1997).
143. Marcsik, A. Szegvár-Oromdűlő 10. és 11. századi embertani leleteinek vizsgálata. *A Móra Ferenc Múzeum Évkönyve– Stud. Archaeol.* **3**, 287–322 (1997).
144. Révész, L. *Heves megye 10–11. századi temetői*. (Magyar Nemzeti Múzeum–Magyar Tudományos Akadémia Régészeti Intézete, 2008).
145. Kustár, Á. A tiszanánai honfoglaláskori temető embertani vizsgálata. *Anthropol. Kozl.* **35**, 119–140 (1993).
146. Költő, L. & Szentpéteri, J. A Vörs-Papkert "B" lelőhely VIII–XI. századi temetője. in *vezredek üzenete a láp világából. Régészeti kutatások a Kis-Balaton területén 1979–1992* (eds. Költő, L. & Vándor, L.) 115–121 (Somogy Megyei Múzeumok Igazgatósága—Zala Megyei Múzeumok Igazgatósága, 1996).
147. Költő, L., Szentpéteri, J., Bernert, Z. & Papp, I. Families, finds and generations: An interdisciplinary experiment at the early medieval cemetery of Vörs–Papkert B. in *Mensch, Siedlung und Landschaft im Wechsel der Jahrtausende am Balaton. People, Settlement and Landscape on Lake Balaton over the millennia. Castellum Pannonicum Pelsonense (CPP4)* (eds. Heinrich-Tamáska, O. & Straub, P.) 361–390 (Verlag Marie Leidorf GmbH, 2014).
148. Köhler, K. *et al.* A Late Neolithic Case of Pott's Disease from Hungary. *Int. J. Osteoarchaeol.* **24**, 697–703 (2014).
149. Osztás, A., Zalai-Gaál, I. & Bánffy, E. Alsónyék-Bátaszék: a new chapter in the research of Lengyel culture. *Doc. Praehist.* **39**, 377–396 (2012).
150. Zalai-Gaál, I., Osztás, A. & Köhler, K. Totenbrett oder Totenhütte? Zur Struktur der Gräber der Lengyel-Kultur mit Pfostenstellung in Südtransdanubien. *Acta Archaeol. Acad. Sci. Hungaricae* **63**, 69–116 (2012).
151. Masson, M. *et al.* 7000 year-old tuberculosis cases from Hungary - Osteological and biomolecular evidence. *Tuberculosis* **95**, S13–S17 (2015).
152. Horváth, F. Hódmezővásárhely-Gorzsa: A settlement of the Tisza culture. in *The Late Neolithic of the Tisza Region* (ed. Raczky, P.) 31–46 (Szolnok Megyei Múzeumok, 1987).
153. Makkay, J. *Vésztő-Mágor. Ásatás a szülőföldön*. (Békés Megyei Múzeumok Igazgatósága, 2004).
154. Spekker, O. *et al.* New cases of probable skeletal tuberculosis from the Neolithic period in Hungary – A morphological study. *Acta Biol. Szeged.* **56**, 115–123 (2012).
155. Lezsák, G. A magyar őstörténet kaukázusi forrásai. in *Magyar Őstörténeti Műhelybeszélgetés* (ed. Neparáczki, E.) 195–246 (Magyarságkutató Intézet, 2020).

#### S2 Supplementary Methods

##### S2.1 Sampling strategy

We present genome-wide data of 271 ancient individuals from the Migration Period of the Carpathian Basin between the 5th and 11th centuries and 3 individuals from the 9-10th century Caucasus with archaeological affinity to the Conquering Hungarians. We also sequenced 3 Neolithic individuals from the Carpathian Basin (summarized in Table S1a).

The 265 Migration Period samples represent the following time range:
9 samples from the Hun Period (5th century),
40 from the early Avar Period (7th century),
33 from the middle Avar Period (8th century),
70 from the late Avar Period (8-9th century).
48 from 10th century Conquering Hungarian elite cemeteries,
65 from commoner cemeteries of the Hungarian conquer-early Árpádian Period (10-11th centuries).

The majority of samples were collected from the Great Hungarian Plain (Alföld), the westernmost extension of the Eurasian steppe, which provided favorable ground for the arriving waves of nomadic groups. Cemeteries and individual samples were chosen on archaeological, anthropological and regional basis. We made an effort to assemble a sample collection from each period representing a) all possible geographical sub-regions, b) all archeological types, c) all anthropological types. From large cemeteries we selected individuals with the same criterions and possibly from all part of the cemetery with the following bias: We preferably choose samples with good bone preservation, archaeologically well described ones (with grave goods) and males (for Y-chromosomal data). Nevertheless, we took care to also include females and samples without grave goods, though these are definitely underrepresented in our collection.

##### S2.2 Population genetic analysis strategy and methods

As the studied samples represent three archaeologically distinguishable periods from three consecutive historically documented major migration waves into the Carpathian Basin, we evaluated Hun, Avar and Conqueror period samples separately.
The newly sequenced genomes were merged and co-analyzed with 2367 ancient (Table S3) and 1397 modern Eurasian genomes (Table S8), most of which were downloaded from the Allen Ancient DNA Resource (Version v42.4)[1]. We also downloaded the Human Origins dataset (HO, 6,2K SNP-s) and/or the 1240K data sets published in[2–4]. As the HO SNPs are fully contained in the larger 1240K set we filtered out the HO data when only the 1240K data set was published. In case the Reich data set contained preprint data of the same individual published later, we always used the published genotypes. Since some data set contained diploid- and mixed call variants we performed random pseudo haploidization of all data prior to downstream analysis.
Most of the analysis was done with the HO dataset, as most modern genomes are confined to this dataset, however, we run some of the f-statistics with the 1240K data if these were available. We used the nomenclature of the Allen Ancient DNA Resource database (Version v42.4) throughout our study, except Russia_IA, which was subdivided into Russia_IA_Tarand (Master ID; VIII5, VIII9, VIII7, VIII8, VII15, and VIII6) and Russia_IA_Tuva (Master ID; RISE504, RISE492, RISE602, RISE600 and RISE601) according to the markedly different origin of these samples. As we regrouped many of the published genomes with Hierarchical Ward clustering (see below) resulting in an increased number of genetic groups from identical cultures, these were distinguished throughout

the analysis with a short form of the original group name plus a serial number added to each subgroup, as listed in Table S3a and 3b.
First line of analysis included PCA[5] and unsupervised Admixture[6], which are the methods of choice to compare genomes in a hypothesis independent manner. Before running Admixture and population genetic analyses we removed close relatives from the dataset (Table S9).
We used Hierarchical Clustering (ward.D2 in cluster R package)[7], 2-dimensional f4-statistics[8] and qpAdm[9] to identify genetically similar individuals which could be grouped for subsequent analysis. Homogeneous genetic groups were further characterized with admixture-f3- and outgroup f3-statistics[10] as well as “distal” and “proximal” qpAdm analysis, in which pre-Iron Age and post-Bronze Age samples were used as sources respectively. Many of our samples were part of genetic clines between East and West Eurasia (Extended Data Fig 1a). In order to reveal the genetic ancestry of individual samples within genetic clines the identified genetic groups at the eastern and western extremes were also added to the Left-populations as sources. Many of the samples within clines could be modelled as simple two-way admixture of these two populations, or three-way admixtures with a third source. The remaining individuals were considered genetic outliers, which could be modelled from different sources. When it was possible, we estimated admixture time with the DATES algorithm[8].

##### S2.3 qpAdm analysis strategy

Next to Hierarchical Clustering we used qpAdm to group genetically similar individuals with shared ancestry as well as to reveal the plausible sources of the studied genomes from each time period. We used the qpAdm tool from the ADMIXTOOLS software package (version 6.0)[9] for modeling admixture sources. Since all of our Test populations were Eurasian samples we used a suitable outlier (Ethiopia_4500BP_published.Sg) as Right Base throughout our analyses.
We have to note that our Middle Age samples are highly admixed (Fig. 2b and Table S4), likewise most of the Iron Age populations from which they possibly derived contain similar genomic components obtained from various admixture histories of related populations. In such cases qpAdm expectedly results in multiple feasible alternative models, most of which is very difficult to exclude. As we wished to include all relevant potential source populations in the analysis, despite their obviously similar genome histories, excluding suboptimal models was the largest challenge of the analysis. Thus, we set out to optimize our qpAdm strategy in multiple steps.
Based on PCA, outgroup f3-statistics and unsupervised Admixture data first we assembled a large set of plausible pre-Bronze Age and early Bronze Age Right populations containing different ancestry components present in our Test populations, for being able to measure shared drift along each component. Next, a large set of Left populations were assembled on the same basis from published genomes of the entire Eurasian region, the number of which was further increased by our PC50 regrouping described above. As the number of possible source populations was very high, testing every possible combination of sources (Left populations) and outgroups (Right populations) rendered this approach impossible. Instead, we chose to run the analysis just with source combinations of 2 and 3 (rank 2 and 3). Theoretically, from a fully representative database (both in time range and geography) all genomes ought to be modelled from one or two proximal sources (rank 1 or 2). As qpWave is integrated in qpAdm, the nested-P values in the log files indicate the optimal rank of the model, that is if P-value for the nested model is above 0.05, the Rank-1 model should be considered[9].
For revealing past population history of the Test populations from different time periods, we run two separate qpAdm analysis. In the so called distal analysis we wished to identify the most plausible distant sources, thus a wide range of pre-Bronze Age and Bronze Age populations were included as sources. Next we run a so-called proximodistal analysis, in which just the most relevant distal sources were left in the Left population list, supplemented with a large number of available post-Bronze Age populations. In latter runs potentially more relevant proximal sources competed with

distant Bronze Age sources, and plausible models with distal sources indicated the lack of relevant proximal sources.
As the number of tested source populations was very high (in the proximodistal analysis it often exceeded 100), th e number of tested 3 source models was on the order of several hundred thousands. This was further increased by a similar number of models in the model competition runs, so in the entire study we run several million qpAdm models.

##### S2.3.1 Optimizing the Right populations

In the first step we performed an iterative optimization of our Right populations based on the log analysis of qpAdm. After the initial qpAdm run for each right populations we collected the number of significant Z scores, the number of models with at least one significant Z score, from the detailed "gendstat" lines of the log files of all qpAdm models. We also counted how many models we would not reject if we excluded the F4 statistics (significant Z scores) of a given right population. Based on this information we could test if all the right populations were needed to reject the models. If some of the Right populations were redundant we removed only one that had significant Z score in the least amount of models (having the smallest prediction power). Next we repeated the qpAdm analysis with the optimized Right populations until all Right populations were needed to reject all rejected models. According to our results the repeated qpAdm analyses with the stricter Right populations always resulted in more models rejected. As an important exception, we always kept Right populations that measured the main genetic components of our test population.
We also moved some Left populations to the Right list if they did not give valid models in any of the initial runs and their PCA position and Admixture composition were similar to our samples, thus were expected to provide significant Z-scores in dscore/gendstat statistics to reject suboptimal models. In most cases we also included modern populations in the Right set, for which post admixture gene flow could be ruled out, as they efficiently rejected suboptimal models. In some cases, it was also unavoidable to use modern populations as sources, because the more relevant ancient sources were seemingly unavailable.

##### S2.3.2 Excluding suboptimal models with model-competition:

In this way we assembled an optimal Right population set for each analysis, giving the smallest number of plausible models, however even the best Right sets still resulted in too many models. In order to further exclude suboptimal models, finally we applied the "model-competition" approach described in[8], the following way: From the Left populations present in the plausible models we moved one at a time to the Right set and rerun qpAdm for each model. This was repeated with each Left population which appeared in any of the plausible models. As the best sources have the highest shared drift with the Test population, including these in the Right list is expected to exclude all models with similar suboptimal sources. This way we were able to filter out most suboptimal models and identify the most plausible ones, which were not excluded by any of the Right combinations.
As each model-competition run gave a different P-value with different standard deviations, we deemed it more informative to provide the maximum, minimum and average P-values for the best final models, instead of the P-values and standard deviations of the original models. We also listed the extra Right populations which minimized or maximized P-values or excluded models, because these are informative.
In some cases, model-competition excluded all 3 source models, indicating that additional sources are required for optimal modeling. However, running 4 source models with numerous sources is not feasible, thus we run 4 source models with a reduced set of Left populations. To select the best candidate subset of sources we evaluated the gendstat data from the log files of the 3 source models and identified populations which excluded the best models. Latter populations presumably had

additional shared drift with the Test, not shared by any of the sources. Thus, in the rank 4 models we included just the best sources from the rank 3 models, plus their excluding populations.

##### S2.3.3 Modeling members of genetic clines:

We qpAdm modelled each sample, most of which were part of genetic clines. First we tried to model cline members as simple two-way admixtures of populations from the two extremes of the clines (as admixed descendants of local EU_Core and immigrant Asia_Core groups, see results). Many samples could be modelled this way, but a large number of samples could not, as they obviously required either a third source or entirely different sources. We applied another strategy to model latter samples.

As three source modeling of large number of samples from large number of sources is infeasible, we decreased computation time the following way. First we decreased the sources to around 50, by selecting one or a few representatives from similar populations in similar PCA locations within the same PC50 cluster. For example, from the large number of eastern Scythian groups we selected two Pazyryk, two Sagly and one Tagar (Table S6d and 7e). Second, we ran constrained 3 source models, by fixing one source at a time, and swapping only the rest of the Left populations. We repeated these constrained runs in 3 combinations, once fixing EU_Core, second fixing Conq_Asia_Core and third fixing Avar_Asia_Core. Finally, we compared the 3 different models for each sample and selected the ones with best P-values. Obtaining the same high P-value models from two different constrained runs, one with EU_Core fixed, the other with Asia_Core fixed, rendered it very likely that the optimal model was found. In cases when EU_Core fixed, and Asia_Core fixed runs gave different models, but all Asia_Core fixed models indicated the presence of EU_Core, we accepted the best EU_Core fixed models. Finally, we also tested whether samples at the Asian side of the cline require entirely different sources by running unconstrained 3 source models for these few samples.

We noticed that model-competition cannot exclude alternative qpAdm models with similar minor components, therefore a given source is best identified from samples which carry it in large fraction. We had several within cemetery PCA clines with varying fractions of the same components, as nearly each member could be modelled from individuals at the extremes of the cemetery cline (Table S6e). In these cases the best Asian source could be accurately identified from the unconstrained 3 source models, which also applies to all members of the cemetery cline.

### S3 Supplementary Results

##### S3.1 Kinship relations:

PCAngsd identified altogether 43 kinship relations and most identified relatives had identical maternal or paternal lineages (Table S9). The few exceptions were distant (>2$^{rd}$ degree) relatives, and uniparental markers never contradicted the potential kin relations.

As expected, most of the relatives were found within cemeteries, but 4 kin pairs were also identified from different cemeteries. Kinship patterns imply patrilocality, as most within cemetery kins were males, while inter-cemetery kins included females, but exceptions indicate that women could also stay within their community. Patrilocality was especially conspicuous in the Hortobágy-Árkus late Avar elite graveyard from which we sequenced 17 genomes, 9 of which were males and 7 of these carried the same Q1a2a1 Y-haplogroup, besides they belonged to two kin-groups, one with 3 brothers. It seems that within the Árkus late Avar elite cemetery most individuals could have belonged to the same extended family.

One example of the inter-cemetery relations is especially relevant from historical perspective, as the Sárrétudvari-Őrhalom (SO) small Conqueror elite cemetery is located close to the Sárrétudvari-

Hízóföld (SH) large commoner cemetery, and the SO-5 man from Őrhalom proved to be a first degree relative of the SH-3 woman from Hízóföld, providing direct evidence for the ongoing admixture between the 10th century Conqueror elite and local commoners.
Another informative example of the inter-cemetery relations came from the neighboring Karos-1 and Karos-2 Conqueror elite cemeteries. Individual K1-3286 from Karos-1 vas a first degree relative of K2-61 from Karos-2, nevertheless they had strikingly different genome profile, indicating a first generation admixture of individuals with different background. While K1-3286 could be modelled from Hun_Asia_Core and Iranian ancestries, K2-61 unambiguously carried 50% Conq_Asia_Core ancestry supplemented with Hun- and Iranian-like elements (Table S7e). Nevertheless, K2-61 could be modelled as first generation admixed progeny of K1-3286 and Conq_Asia_Core (Table S7d), confirming the ongoing admixture of immigrants with Hun-related and Conqueror ancestries still in the 10th century.
We kept just one of the related individuals with best genome coverage for further analysis.

##### S3.2 Defining EU_Core groups

On PCA most of our samples from each period map to the European side (Fig. 2 and Extended Data Fig.1b) overlapping with ancient and modern European populations. It is conspicuous that our samples at the European side form a South-North cline (Extended Data Fig 1b), which we termed the EU-cline. The southern border of this cline corresponds the PCA position of modern Greeks, Albanians, Italians as well as ancient Neolithic and Chalcolithic samples from the Carpathian Basin. The northern border of the cline well corresponds to the position of modern Hungarians and Late Bronze Age-Iron Age samples from Hungary. It is also remarkable that the southern half of this cline is dominated by Avar Period samples while the northern half by Conqueror Period samples (Extended Data Fig.1b).
PC50 clustering identified 4 homogenous groups within the EU-cline (Table S3). We performed a preliminary distal qpAdm analysis for each EU-cline sample (data not shown), which supported PC50 clustering and indicated that the largest cluster can be subdivided. As a result, we established five EU-cline groups representing five different genome compositions. Next, we selected a few representatives from each group, with most equivalent qpAdm models, and merged these genomes under the name of EU_Core1 to EU_Core5 respectively.
For unsupervised ADMIXTURE we present the lowest cross validation error model, K=7. The 7 admixture components are maximized in the following populations: modern Han (best represented by She), modern Nganasan, European Western Hunter-Gatherers (WHG), Ancient North Eurasians (ANE, represented by Afontova Gora 3, MA1 and Botai), Iranian early farmers (Iran_N), Anatolian early farmers (Anat_N), and European early farmers (EU_N), though latter two categories obviously overlap (Fig. 2b and Table S4).
Samples in the EU-cline typically contain negligible Asian (Nganasan and Han/She) ADMIXTURE components, and their ADMIXTURE compositions are comparable to that of Bronze and Iron Age populations from Europe and the Carpathian Basin, as well as to modern Europeans. (Fig. 2 and Table S4).

**EU_Core1:** This group is located at the southernmost part of the EU-cline, and their PC2 position overlaps with modern Greeks, Albanians, Italians and European Neolithic-Chalcolithic samples. This group is best represented by samples; ALT-224, KK1-251, KK1-252, SZK-83, SZK-180 and SZOD-376 (Extended Data Fig.1b), all of them from the Avar period, except for the 11th century commoner SZOD-376.
On PC50 clustering EU_Core1 clusters together with Langobards from Hungary[11], Iron Age, Imperial and Medieval individuals from Italy[12], as well as with Minoans and Mycenaeans from Greece[13] (Table S3), indicating an ancient southern European genetic affinity of this group.

**EU_Core2:** This group is located on PCA just above EU_Core1, and is best represented by samples HH-102, SZKT-265, SZKT-311, ALT-414 and ARK-38, all from the Avar period. On PC50 clustering they are clustered among others with Hungary_Langobard[11], Hungary_Maros_EBA, Hungary_MBA_Vatya[14] and Croatia_EIA[15] individuals, pointing at the presence of similar genomes in the Carpathian Basin already in the Bronze Age.

**EU_Core3:** This group is located on PCA just above EU_Core2, and is best represented by samples PLE-195, SH-175, SZF-43, and TMH-388, the first two from 11th century commoner, the last two from Avar period cemeteries. On PC50 clustering they are clustered among others with Czech_BellBeaker[16], Hungary_EBA_Protonagyrev[16], Hungary_MBA_Vatya[14] and Italy_IA_Republic[17] individuals, allowing the same conclusion as before.

**EU_Core4:** This group is located on PCA just above EU_Core3 and is best represented by samples TCS-18 (Conqueror elite), SZA-7 (11th century commoner), KDA-485 (middle Avar) and OBT-3 (late Avar). On PC50 clustering they are clustered among others with Bronze Age European Bell Beaker individuals from the Czech Republic, Poland, Iberia and Germany[16].

**EU_Core5:** This group is located on PCA next to EU_Core4, at the northernmost part of the EU-cline, overlapping with modern Hungarians. EU_Core5 is best represented by samples SE-64 (Conqueror elite), IBE-206, VPB-600 (both 11th century commoners) and TMH-199 (late Avar). On PC50 clustering they are clustered among others with Hungary_EBA_BellBeaker[16], Germany_EMedieval[18] and Hungary_Scythian[19] individuals.

As comparable ancient genomes had been present in the Carpathian Basin and the surrounding region before the Migration Period, it is very likely that all EU_Core samples represent local residents. Unsupervised ADMIXTURE revealed a gradient-like shift of genomic components along the EU-cline (Fig. 2b) with increasing Ancient North Eurasian (ANE) and Western Hunter-Gatherer (WHG) and decreasing Iranian early farmer (Iran_N) and European early farmer (EU_N) ancestries from South to North. All genomes within the EU-cline form transitions between the core groups, as they could be qpAdm modelled as admixtures of these (Table S6f and 7f).
In the subsequent analysis we considered the EU_Core groups as local residents of the Carpathian Basin. Though similar European genomes could be present on the Medieval Pontic Steppe as well, at present this possibility cannot be distinguished with available methods.

##### S3.3 The Conqueror cline:

Conquest period samples form a characteristic genetic cline on PCA, which we named the "Conqueror-cline" (Conq-cline, green and blue on Fig.2a and Extended Data Fig.1a). The Conq-cline largely overlaps with the modern "steppe-forest cline"[20], but it runs from Europe just to the middle of the "steppe-forest cline" corresponding to the PCA position of modern Bashkirs and Tatars. The Conq-cline principally overlaps with the ancient steppe cline published in[19], comprised of Bronze-Age Okunevo, Karasuk, Mezhovskaya and Iron Age Scytho-Siberian, Saka, Sarmatian, Wusun, Cimmerian, as well as younger Kangju, Late Xiongnu, TienShan Hun, and Medieval Nomad samples.
Unsupervised ADMIXTURE revealed that Conqueror genomes are very diverse and complex, incorporating all 7 components in different proportions (Fig. 2 and Table S4). About half of the samples contain the Nganasan Siberian component, which is present both in the elite and the commoners, and which is largest in samples at the Asian extreme of the cline, while, as shown above, samples at the European side lack Nganasan and Han/She elements.

#### S3.4 Defining Conqueror_Asia_Core

##### S3.4.1 Hierarchical clustering

PC50 clustering grouped members of the Conq-cline into several clusters, and at the Asian extreme of the cline 12 individuals were tightly clustered together, indicating that they shared highly similar genomes (Table S3). We termed this population Conq_Asia_Core, which we subsequently divided into two subgroups, as 4 samples were definitely shifted downwards from the rest on PCA (Fi. 2a, Extended Data Fig. 1a) due to their somewhat higher Han and Ancient North-East Asian (ANA) related ancestries, as revealed by two-dimensional f4-statistics (see below).

Members of Conq_Asia_Core1 are: MH1-23, KeF2-1045, KeF1-10936, TCS-2, LB-1432, SZAK-4, SZAK-6 and SZAK-7. Members of Conq_Asia_Core2 are SZAK-1, K3-6, K2-29 and SZA-154. These groups together consist of 6 males and 6 females derived from 9 different cemeteries and 11 of the 12 individuals belonged to the Conqueror elite according to archaeological evaluation (Table S1a).

On PC50 clustering Conq_Asia_Core clusters together with Bronze Age Okunevo and Karasuk samples from the Minusinsk Basin[14], Iron Age Scythians from Tuva, Pazyryk and the Altai[4], Sargat from Trans-Urals[4], Sagly-Uyuk from the Mongolian Altai[2], Central-Saka from Kazakhstan[4,19] and Xiongnu from Western Mongolia[2,19] (Table S3). The Conq_Asia_Core group very plausibly represented 10$^{th}$ century immigrants.

##### S3.4.2 PCA and unsupervised ADMIXTURE

On PCA closest to Conqueror_Asia_Core we find modern Bashkir and Tatar groups: Bashkir_Kugarchinsky, Bashkir_Ishimbai, Bashkir_Baimaksky, Tatar_Siberian_Yalutorovskiy, Tatar_Volga, Tatar_Siberian_Tyumen and Tatar_Siberian_Tomsk (Fig. 2a) in agreement with our previous results from uniparental markers[21,22], which identified Baskirs and Tatars as the most similar populations to the Conqueror elite.

Unsupervised ADMIXTURE detected 33-40% Nganasan, 27-34% ANE, 11-16% EU_N, 7-12% Anat_N, 1-8% Han, 1-8% Iran_N and 0-4% WHG components in Conq_Asia_Core individuals, and they showed the highest similarity with populations in the same PC50 cluster (Fig. 2b, Table S4). These results indicated a potential affinity of the Conq_Asia_Core to eastern Scythians, as well as a likely geographic location between the Urals and the Altai somewhere in northern Kazakhstan.

##### S3.4.3 f3-statistics

Admixture f3-statistics in the form f3(Conq_Asia_Core; X, Y) measured most negative f3 values with modern Nganasan and ancient or modern European populations (Extended Data Fig. 5a), Nganasan-MLBA_Sintashta having the 4$^{th}$ largest Z-score (-21,153). This result indicates that the main admixture sources of the Conq_Asia_Core population were Steppe_MLBA and Siberian populations, and best proxy of latter is modern Nganasan.

Next, we performed outgroup f3 statistics in the form f3(Mbuti; Conq_Asia_Core, Y) to measure shared drift between Conq_Asia_Core and all available modern and ancient populations. We found that modern and ancient Siberian populations share highest drift with Conq_Asia_Core (Extended Data Fig. 5c), moreover the top 4 of these are modern populations speaking Uralic languages; Mansi (Ugric), Nganasan (Samoyedic), Selkup (Samoyedic), Enets (Samoyedic). This result revealed that Conq_Asia_Core shared evolutionary past with language relatives of modern Hungarians, and the closest language relatives, Mansis had one of the the highest shared drift.

It is notable, that along the P1 PCA axis the Conqueror_Asia_Core group falls not far from modern Mansis, however along the P2 axis Mansis are further away, they are members of the Siberian genetic cline, while Conquerors fall in the steppe-forest cline (Fig. 2a). Mansis also have significantly different Admixture components from Conquerors, as they lack the Han/She, Iran_N, and Anat_N components present in the Conquerors (Fig. 2b). We also performed outgroup f3 statistics in the form f3(Mbuti; Mansi, Y) and the results revealed that the top list of populations sharing highest drift with Mansis are comparable to that of Conqueror_Asia_Core (Extended Data Fig. 5e), however latter is ranked much lower ($118^{th}$) in the list, consistent with the genome differences shown above. These results exposed that genetic relations support language relations, and Mansis need to be co-analyzed with Conquerors. Besides, ancestors of Mansis maybe ancestors of Conq_Asia_Core too, while the opposite is unlikely.

##### S3.4.4 Two-dimensional f4-statistics

We run several two-dimensional f4-statistics with the 1240K marker set to detect slight genetic differences between Conq_Asia_Core individuals, obtained via multiple gene flow. These f4 statistics in the form of *f4(Ethiopia_4500BP, Test ; Sintashta, Ulaanzuukh_SlabGrave)* versus *f4(Ethiopia_4500BP, Test ; Sintashta, Miao_modern)* indicated that Conq_Asia_Core individuals had different proportion of ancestry related to Miao (a modern Chinese group) and *Ulaanzuukh_SlabGrave*, latter representing Ancient North Asians (ANA)[2], and Conq_Asia_Core2 individuals had highest ancestry of both (Extended Data Fig 4a). Besides, individuals are arranged linearly along the Miao-ANA cline, suggesting that these ancestries covary in the Conqueror group, thus could have arrived together, most likely from present day Mongolia.
Along the Ancient North Eurasian (ANE)[10] axes represented by Botai[23], and the Bactria-Margiana Archaeological Complex (BMAC, represented by BA_Bustan[8]) axes the samples showed a more scattered arrangement (Extended Data Fig 4b), indicating that ANE and BMAC ancestry components poorly distinguish Conq_Asia_Core individuals, though Conq_Asia_Core2 individuals showed somewhat higher BMAC ancestry.

##### S3.4.5 distal qpAdm

While optimizing the Right populations in order to reveal the origin of the Conq_Asia_Core group (detailed in point S2.3.1) we run several preliminary qpAdm analysis with a diverse set of Left populations containing various Eurasian Bronze-Age and Iron Age samples. All analysis excluded proximal European sources, confirming that these individuals indeed represented new immigrants in the Carpathian Basin from outside Europe.
Next, we run a distal qpAdm analysis, in which we tested 44 potential Bronze Age sources in all combinations of 3 (rank 3), to identify potential distant sources of the two Conq_Asia_Core groups as well as for Mansis, with the following optimized Right population set (Table S7a):

*Ethiopia_4500BP, China_Tianyuan, ANE (AfontovaGora3+ Russia_MA1), Iran_GanjDareh_N, Kolyma_M, Anatolia_N, Latvia_HG, DevilsCave_N, Baikal_EN (Lokomotiv+Shamanka), WSHG (Tyumen+Sosnoviy), Russia_Steppe_Maikop, Karitiana, Chukchi*

We obtained 1056 plausible models for Conq_Asia_Core1, 539 models for Conq_Asia_Core2, and 85 models for modern Mansi. Next we performed a model-competition experiment[8] by rotating each Left populations to the Right list one by one, as described above in point S2.3.2. This way we obtained a single passing model for Mansis, which was the top P-value hit among the initial passing models.

***Mansi:***
*48% Russia_Mezhovskaya, 44% Nganasan, 8% Kazakhstan_Eneolithic_Botai*
(max P: 0.29, min P: 0.094, average P: 0.22)

For Conq_Asia_Core1 none of the models passed after model competition. The best model below was excluded only once, by the *Mongolia_LBA_CenterWest_4D* individual[3].

***Conq_Asia_Core1:*** *55% Russia_Mezhovskaya, 24% Nganasan, 21% Altai_MLBA_o*
(max P: 0.43, min P: 0.07, average P: 0.26)

Finally, we tested a 4-source model for Conq_Asia_Core1, selecting a set of 12 Left populations by the criteria detailed in the qpAdm analysis strategy section, and after model competition in the best passing model the former excluding *Mongolia_LBA_CenterWest_4*D individual was the 4$^{th}$ source, though out of the 44 competition-runs this model was also excluded once, by another *Mongolia_LBA_4_CenterWest* sample (Table S7b):

***Conq_Asia_Core1:***
*52% Russia_Mezhovskaya,SG, 20% Altai_MLBA_o, 15% Mongolia_LBA_CenterWest_4D, 13% Nganasan*
(max P: 0.23, min P: 0.09, average P: 0.16)

Our *Altai_MLBA_o* group consists of two ancient individuals, KHI001.A0101 and I13173 excavated not far from each-other in the Mongolian Altai. The unclassified Altai_MLBA outlier (KHI001.A0101), had been modelled as *30% Sintashta, 60% Baikal_EBA, 10% Gonur1_BA*[4]. The I13173 *Mongolia_MBA_Munkhkhairkhan_2* individual had been modelled as *30% WSHG, 50% Mongolia_East_N, 20% Turkmenistan_Gonur_BA_1*[3]. These genomes are more similar to Iron Age Scytho-Siberians than to contemporary MLBA genomes, as latter lacked Gonur_BA (BMAC) ancestry.
These results indicate that the Late Bronze Age Russia_Mezhovskaya and Nganasan populations could be common ancestors of Mansis and Conq_Asia_Core1, and their admixed descendants could indeed form a “proto-Ugric” community as inferred from linguistic data[24,25]. Above model also revealed the difference, a much higher Nganasan ancestry in Mansis, which was largely replaced by a Scytho-Siberian like ancestry in Conq_Asia_Core1, including BMAC components derived from the Altai-Mongolia region.

For Conq_Asia_Core2 the model-competition approach resulted in a single passing 3-source model (Table S7a):

***Conq_Asia_Core2:***
*63% Mongolia_LBA_MongunTaiga_3B, 27% Karasuk, 10% Magadan_BA*

The *Mongolia_LBA_MongunTaiga_3B* individual (I12976) was published in Wang et al. 2021[3]. Mongun Taiga type graves were excavated in the Mongolian Altai, and they belong to the wider Altai_MLBA group. The deceased were of Europoid anthropological type, with traces of Mongoloid features, related to Sintashta-Andronovo people. Some archaeologists consider them as the earliest stage of the Sagly-Uyuk Scytho-Siberian culture. The *Mongolia_LBA_MongunTaiga_3B* sample had been modelled as *50% Mongolia_East_N, 36% Russia_Sintashta_MLBA, 8% Russia_Afanasievo, 6% Turkmenistan_Gonur_BA*_1[3], therefore this result also indicates Scytho-Siberian like ancestry in Conq_Asia_Core2.

In the first run we had an alternative model for Conq_Asia_Core2, which is more comparable to that of Mansi and Conq_Asia_Core1:

***Conq_Asia_Core2:***
*54% Russia_Mezhovskaya.SG, 36% Nganasan, 10% Uzbekistan _BA_Bustan* (P= 0,445)

Though this model did not pass after model competition (Table S7a), yet it signifies the main differences between the two Conq_Asia_Core groups: higher Siberian and BMAC ancestry in Conq_Asia_Core2, which is also visible on unsupervised ADMIXTURE (Fig. 2b and Table S4) and 2-dimensional f4-statistics (Extended Data Fig 4).

In summary, distal qpAdm results supported the common Mezhovskaya-Nganasan related evolutionary past of Conq_Asia_Core and modern Mansis and pointed at an additional ancestry from present day Mongolia in Conquerors, which is absent from Mansis. For this reason, we included Mansis as sources in the subsequent proximodistal analysis.

##### S3.4.6 proximodistal qpAdm

In the proximodistal analysis we tested 99 potential source populations. Most of the sources came from the Iron Age Steppe, but we also included Mansis, and 9 populations from the Bronze Age, using the following optimized reference (Right) populations (Table S7c):

*Ethiopia_4500BP, ANE (AfontovaGora3+ Russia_MA1), Iran_GanjDareh_N, Anatolia_N, Latvia_HG, DevilsCave_N, Baikal_EN (Lokomotiv+Shamanka), WSHG (Tyumen+Sosnoviy), Russia_Steppe_Maikop, Finnish (modern), Ket (modern), Han (modern)*

This Right set contains three modern populations, which very efficiently excluded suboptimal models in our preliminary qpAdm experiments. From this list Ket has very similar ADMIXTURE profile to Mansis (Table S4), but never appeared as source in any of the preliminary plausible models. We did not obtain significant nested P-values in the 3-source models, indicating that 2 sources are insufficient to model Conq_Asia_Core from the used Left populations.

We obtained 624 plausible three source models for Conq_Asia_Core1, of which 17 passed after model competition (Table S7c). It is remarkable, that both the source populations and their proportions are very similar in each model. The first two majority sources are practically identical, while the third sources represent similar genomes from two different time periods, Scythians elevated ANA ancestry from the Iron Age, or Hun-related samples from later periods.

***Conq_Asia_Core1:***
*46-52% Mansi, 25-40% Early/Late Sarmatian, 12-26% Scyho-Siberian/Xianbei/Late Xiongnu.*
(MaxP: 0.8, MinP: 0.08, averageP: 0.63)

It is telling that all of the Early/Late Sarmatian samples were discovered around the southern Ural region[26,27], and had virtually identical genomes.
The 12-26% third populations included:

*Tasmola_Korgantas_300BCE* (KBO001.A0101 or BDY001.A0101)[4], *Pazyryk_Berel_50BCE_o* (BRE002.A0101 and I13504)[3,4],
*Tasmola_Birlik_640BCE* (BIR010.A0101 and BIR013.A0101)[4]
*SlabGrave* 1562-407 BCE ( PTO001.A0101, DAR001.B0101, ULN005.A0102, I6369 and I6368)[2,3]

*Xianbei_Hun_Berel_300CE* (BRE014.A0101) )[4],
*late_Xiongnu* 50 BCE - 100 CE (IMA007.A0101, IMA008.B0101, UVG001.A0101, CHN010.A0101 and I6228) )[2,3], or
*Kazakhstan_Nomad_Hun_Sarmatian* 334-535 CE (DA27)[19].

These groups are located close to each other on PCA and have very similar ADMIXTURE profile (Table S4).
Though early and late Sarmatians have similar genomes, they define two incompatible time-frames around 400-200 BCE versus 100-300 CE. The third populations also define two incompatible time-frames; around 640-50 BCE involving Scytho-Siberian populations, or around 50 BCE-500 CE involving Late-Xiongnu/Xianbei/Hun populations.

For Conq_Asia_Core2 we obtained 985 plausible models, of which 8 passed after model competition. Most of the passing models had very similar sources to that of Conq_Asia_Core1, with shifted proportions:

***Conq_Asia_Core2:***
*21-37% Mansi, 29-54% Sarmatian, 23-37% Xiongnu/Xianbei*
(MaxP: 0.47-0.79, MinP: 0.06, averageP: 0.35-0.67)

In three models the best proxy for Sarmatian was a late Xiongnu group from Mongolia (Xio_Sarm: UGU010.A0101, TMI001.B0101, BUR003.A0101), which had been shown to have indistinguishable genomes from Pontic Sarmatians[4] but in 3 models Kangju (DA121 and DA206), Kazakhstan_Karluk[19] and Ukraine_IA_WesternScythian[26], samples replaced Sarmatians.

The third populations included the same groups as in the Conq_Asia_Core1 models: Late Xiongnu (IMA007.A0101, IMA008.B0101, UVG001.A0101, CHN010.A0101, I6228) which had been modelled as *90% Ulaanzuukh_SlabGrave, 10% Chandman_IA/Sarmatian*[4],
*Xianbei_Hun_Berel_300CE* (BRE008.A0101 or BRE014.A0101), and *Tasmola Korgantas_300BCE (*BDY001.A0101)[4].
Alternatively the third populations included *Early Xiongnu* (SKT012.A0101)[2],
or Late Xiongnu_Han (BRU001.A0101) modelled as *41% Ulaanzuukh_SlabGrave, 32% Han, 26% Sarmatian* in the same paper.

It is reasonable to suppose that Conq_Asia_Core1 and Conq_Asia_Core2 individuals received the same influx from the same sources, but in different proportions, thus it is reassuring that the three source models identified very similar source populations for both groups. Moreover, the Conq_Asia_Core2 passing models rather support a Xiongnu/Xianbei/Hun_Sarmatian source than Iron Age sources. As the source populations in the models define inconsistent time periods, we performed DATES analysis to clarify admixture time.

##### S3.4.7 Dating admixture time with DATES

According to our experience DATES works best with high coverage genomes, or when multiple ancient genomes are merged for the analysis. Therefore, we merged all similar genomes identified in the 3 source qpAdm models, to find out the Mansi+Sarmatian versus Mansi+Scytho-Siberian/Xiongnu admixture times in the Conq_Asia_Core population.

First we dated the Mansi-Sarmatian admixture time, and for this end the following Early and Late Sarmatian genomes were merged from the passing qpAdm models:

Late Sarmatian: tem002, tem003, tem001, chy001[27]
Early Sarmatian: MJ-43, MJ-39, LS-13, MJ-41, MJ-56[26]

We also merged all Mansi genomes: Mansi48, Mansi76, Mansi94, Mansi43, Mansi79, Mansi91, Mansi84, Mansi56[28], S_Mansi-2.DG[29].

We also merged the two Conq_Asia_Core populations, as they reasonably acquired genetic influxes at the same time, moreover this way we could increase sample size.

For the Mansi-Sarmatian admixture we obtained 53.117 mean generation time (Extended Data Fig. 6), and calculating with a 26-30 years generation time[30] this corresponds to 1381-1593 years before death (which is estimated around 950 AD), or between 643-431 BCE admixture date, which corresponds to the early Sarmatian period.

For the Mansi-Scythian/Xiongnu/Xianbei DATES analysis we used the third populations from the Conq_Asia_Core1 three source models, and merged those genomes which gave comparable proportions (11-14%) and had adjacent locations on PCA. In this way we merged the following genomes:

*Tasmola Korgantas_300BCE* (KBO001.A0101),
*Pazyryk_Berel_50BCE_o* (BRE002.A0101, I13504) ,
*Tasmola_Birlik_640BCE* (BIR013.A0101),
*Slab Grave* 1562-407 BCE (PTO001.A0101, DAR001.B0101, ULN005.A0102, I6369, I6368),
*Xianbei_Hun_Berel_300CE* (BRE014.A0101),
*Late Xiongnu* 50 BCE - 100 CE (IMA007.A0101, IMA008.B0101, UVG001.A0101, CHN010.A0101, I6228)
*Kazakhstan_Nomad_Hun_Sarmatian* 334-535 CE (DA27).

The analysis was done with the same merged Mansi and Conq_Asia_Core genomes as listed above. For the Mansi-Scythian/Xiongnu admixture we obtained 24.431 mean generation time (Extended Data Fig. 6). Calculating with a 26-30 years generation time this corresponds to 635-733 years before death, or between 217-315 CE admixture date. From the identified 3rd source populations the archaeological age of *Xianbei_Hun_Berel_300CE* and *Kazakhstan_Nomad_Hun_Sarmatian* (334-535 CE) fit to these data.

Apparently this second admixture could have happened after the late Xiongnu period, during the formation of European Huns, before the Huns arrived at the Volga region and integrated local tribes east of the Urals, including Sarmatians and Conquerors. The same event had also been inferred from historical and archaeological data as described in[31]: "*In fact archaeological evidence from the Ural region seems to point to the expansion of the Huns into that area by the early fourth century AD, suggesting that the nations between the Altai and the Urals had succumbed to Hunnic conquest by the early fourth century"*

As a consequence, Conq_Asia_Core acquired a Scytho-Siberian like genome not directly from eastern Scythians, but from similar sources at a later time. These data are compatible with a permanent location of the Conquerors around the Ural region, in the vicinity of early Sarmatians, along the migration route of the Huns. Recently extensive phylogenetic connections were detected between the conquering Hungarians and individuals of the Kushnarenkovo-Karayakupovo culture from the Trans-Uralic Uyelgi cemetery [32], also confirmed by our previous study [33]. Our genome data seem consistent with this geographic location, so the Trans-Urals is the best candidate motherland of the Conquerors.

**S3.5 qpAdm modeling of Conqueror cline members**

PCA genetic clines generally represent a transition series of admixed genomes between genomes at the two extremes of the cline. In our case Conq_Asia_Core and EU_Core are located at the two ends of the cline, former certainly representing immigrant Conquerors, while latter very likely representing local residents. As most members of the Conqueror cline were derived from 10-11$^{th}$ century commoner cemeteries it was reasonable to suppose that many of these could have been 2$^{nd}$-5$^{th}$ generation admixed progenies of the immigrants and local residents. These freshly admixed individuals ought to be modelled as simple two-way admixtures of Conq_Asia_Core and EU_Core, or maybe three-way admixtures with a third source.
We tested this possibility with qpAdm in which Conq_Asia_Core, Avar_Asia_Core (see later) and all EU-Core groups were included in the Left population (Table S7d) using the following Right population set:

*Ethiopia_4500BP, Iran_GanjDareh_N, Anatolia_N, Romania_Mesolithic_IronGates, Latvia_HG, EEHG (Karelia+Samara+Popovo), modern Tajik, Han, Ket.*

Altogether 31 Conqueror Period samples from the Conqueror cline could be modelled as simple two-way admixtures of Conq_Asia_Core and EU_Core, of these 11 belonged to the elite, while 20 to the 10-11$^{th}$ century commoners (summarized in Table S1b).

The rest obviously required either a third source population or entirely different sources, and we analyzed these as described in the qpAdm strategy section.
Two other elite samples could be modelled from Conq_Asia_Core and EU_Core, plus a 3$^{rd}$ Hun/Xiongnu related source, and one commoner from Conq_Asia_Core and EU_Core plus an Iranian related source. Two elite samples did not show EU_Core components, and these were modelled from Conq_Asia_Core and Hun/Xiongnu related plus Iranian-related sources.

We call Hun-related sources *Hun_Asia_Core* (Fig 2b, see below), *Kazakhstan_Nomad_Hun_Sarmatian*[19], *Xianbei_Hun_Berel_300CE*[4], and *Xiongnu*[2] samples, which have similar genomes, each with major ANA ancestry.

As for the Iranian-related sources, we regularly obtained a set of 3$^{rd}$ sources, which qpAdm was unable to distinguish, each containing significant Iranian ancestry. These include Alan, our Saltovo-Mayaki-like Anapa samples from the Caucasus (see below), Tian Shan Huns, Tian Shan Sakas, Karluks and late Xiongnu samples with considerable BMAC ancestries[2] (*lateXiongnu_Sarm_BMAC, lateXiongnu_Sarm_Ulaa_BMAC* and *lateXiongnu_Sarm_Uyuk_BMAC*, see Table S3). We summarized these under the name of Scytho-Iranian in Table S1b.

The rest 17 samples from the cline lacked Conq_Asia_Core ancestry, thus these can be regarded outliers. It is notable that 9 of the outliers were mapped into the Avar-cline, and the others were also shifted towards the Avar-cline on PCA (Extended Data Fig 1a). Five outliers could be modelled from EU_Core and Avar Asia_Core plus a 3$^{rd}$ Iranian-related source, while in 9 other similar models Hun-related samples provided a better fit than Avar_Asia_Core. One elite individual (AGY-92) with the most extreme Asian PCA location was modelled from two different Hun-related sources, plus a 3$^{rd}$ Iranian source, while the distinguished leader from the Karos-2 graveyard (K2-52) was modelled as first-generation EU_Core-Alan admixed progeny (Table S7d).

These results revealed that the Conquerors were assembled from several populations with diverse genetic background, corresponding to our historical knowledge about the blood-contract between uniting tribes. Our data indicate that most of the associated peoples had Hun and Avar genetic background. It seems from our data, that Conqueror elite individuals with Hun-related genomes were clustered in certain cemeteries, for example each sample from Szeged-Öthalom (SEO), Algyő 258-as kútkörzet (AGY), Nagykőrös-Fekete-dűlő (NK), Sándorfalva-Eperjes (SE), Sárrétudvari-Poroshalom (SP) had this ancestry.

Summarizing the genetic makeup of the Conqueror period samples from Table S1b, we found that approximately 25% of the elite carried Conq_Asia_Core genomes, 25% Hun/Avar related genomes, 20% were locals with pure EU_Core makeup, while 30% were admixed between locals and immigrants. The same calculation for the commoners gives 55% local (pure EU_Core), 40% admixed between locals and immigrants, 3% Avar/Hun related, and ~2% Conq_Asia_Core-like genome distribution. These numbers imply that most individuals in the commoner cemeteries represented local individuals as had been inferred from mitogenome data[33] and that the Conquerors rapidly integrated the local population via intermarriage.

Our data allows a rough estimation of the proportion of 10$^{th}$ century immigrants compared to the local population. Assuming that our samples are representative for each cemetery, the proportion of Conq_Asia_Core ancestry in the combined genomes in all cemeteries, extrapolated to grave numbers gives a 17% proportion. This estimation is certainly not exact and can be regarded at best as an upper boundary, because elite samples are certainly overrepresented in our data, and Transdanubia with its much higher local background was not studied.

##### S3.6 The Avar-cline

The Avar-cline visible in brown on Fig. 2 and Extended Data Fig 1b runs between the Conqueror and modern Turkic clines – partially overlapping with both – and it reaches the Eastern end of the “steppe-forest cline”, where we find modern Buryat, Kalmyk, Tuvinian, Khamnegan, Yakut and Mongol groups. This indicates that Avar immigrants had considerably different genome history from that of the conquering Hungarians.

Except for its eastern boundary, significantly less ancient samples fall in the Avar-cline than in the Conqueror cline, these include mainly Xiongnu samples from Mongolia, Medieval Golden Horde, Karluk, Turk and Medieval nomad individuals from Kazakhstan, as well as several Tien Shan Huns, Sakas, Tasmola and Scytho-Siberian outliers. The downwards shift of the Avar samples along PC2 implies that Avar genomes represent a shift from Bronze-Iron Age steppe genomes towards modern Turkic genomes.

ADMIXTURE analysis revealed that the complexity of the Avar genomes is comparable to that of the Conqueror’s, having all 7 admixture components in different proportions (Table S4).

##### S3.7 Defining Avar_Asia_Core

###### S3.7.1 Hierarchical clustering and ADMIXTURE

We classified the Avar-cline the same way as described above for the Conqueror cline. PC50 clustering identified a single genetic cluster at the Asian extreme of the cline with 12 samples, derived from 8 different cemeteries, which we termed Avar_Asia_Core (Table S3 and Fig. 2). According to archeological evaluation, 10 samples of these were assigned to the early Avar period (**CS-465**, **FGD-4**, **KFP-7**, **KV-3369**, KFP-31, KFP-30a, AN-376 FU-215, MM-245 and CSPF-114), the first 4 highlighted in bold belonging to the elite, one sample (CSPF-37) was assigned to the middle Avar period, and one (CSPF-213) to the late Avar Period. Males were overrepresented in this group, as 9/12 of these individuals were males (Table S1a).

Avar_Asia_Core is PC50-clustered together with very ancient samples from the Baikal region, like Shamanka_Eneolithic, Lokomotiv_Eneolithic[23], Fofonovo_EN[2] and Mongolia_N[3], as well as younger samples including Slab Grave, Xiongnu, Xianbei, Tasmola Korgantas and Pazyryk Berel outlier individuals.

ADMIXTURE identified 64-67% Nganasan, 26-33% Han, 5-11% Anat_N, 0-3% EU_N, 0-3% ANE components, while Iran_N and WHG ingredients were entirely missing from this group (Fig. 2). Ancient samples with most similar ADMIXTURE profiles are those which are in the same PC50 cluster; Shamanka_Eneolithic, Lokomotiv_Eneolithic and Slab Grave, indicating that ANA ancestry dominates these genomes. The PCA position and ADMIXTURE profile of Avar_Asia_Core genomes unambiguously indicated East Asian origin, most likely present day Mongolian.

##### S3.7.2 f3-statistics

We run outgroup f3-statistics in the form f3(Mbuti; Avar_Asia_Core, Y) to measure shared drift between Avar_Asia_Core and all available modern and ancient populations with the 1240K marker set. We found that Avar_Asia_Core had highest shared drift with *earlyXiongnu_rest, late_Xiongnu, Ulaanzuukh_SlabGrave*[2] and *Kazakhstan_Nomad_Hun_Sarmatian*[19] genomes (Extended Data Fig 2a nad 2b), which had been shown to carry predominantly ANA ancestry[2].

##### S3.7.3 Two-dimensional f4-statistics:

We run two-dimensional f4-statistics with the 1240K marker set, to detect minor genetic differences between Avar_Asia_Core individuals, obtained via multiple gene flow. Considering the major ANA ancestry and likely Mongolian location, it was reasonable to check the possible traces of Iranian (BMAC), Steppe and ANE influxes. Thus we performed f4 statistics in the form of *f4(Ethiopia_4500BP.SG, Test ; Ulaanzuukh_SlabGrave, Russia_MLBA_Sintashta)* versus *f4(Ethiopia_4500BP.SG, Test ; Ulaanzuukh_SlabGrave, Uzbekistan_BA_Bustan)* or *f4(Ethiopia_4500BP.SG, Test ; Ulaanzuukh_SlabGrave, Kazakhstan_Eneolithic_Botai.SG*).
The results indicated that Avar_Asia_Core individuals were well separated along a BMAC-Steppe_MLBA cline (Extended Data Fig 3a), and 3 individuals had negligible proportion of ancestry related to BMAC and Sintashta. The same result was obtained from the Steppe_MLBA-ANE statistics (Extended Data Fig 3b), indicating that Iranian, Steppe and ANE ancestries covaried in the Avar_Asia_Core group, thus could have arrived together.
As we detected notable differences between the Avar_Asia_Core individuals, and 3 of them with smallest Iranian and Steppe ancestries also visibly separated on PCA, we set apart these 3 individuals (KFP-31, CSPF-114 and KFP-7) under the name of Avar_Asia_Core1, while the other 9 samples (CS-465, FGD-4, KV-3369, KFP-30a, AN-376, FU-215, MM-245, CSPF-37 and CSPF-213) were regrouped as Avar_Asia_Core2 (Extended Data Fig 1a).

##### S3.7.4 distal qpAdm:

In the distal analysis we tested 64 East Eurasian Left populations dated to the pre-Iron Age, mostly from Mongolia, using the following reference (Right) population set (Table S6a):

*Ethiopia_4500BP, China_Tianyuan, ANE (AfontovaGora3+ Russia_MA1, Iran_GanjDareh_N, Anatolia_N, DevilsCave_N, WSHG (Tyumen+Sosnoviy), EEHG (Karelia, Samara, Popovo), modern Koryak, modern Onge*

The initial analysis yielded 107 rank 2 models for Avar_Asia_Core1, and 138 for Avar_Asia_Core2, and after model-competition we obtained 7 passing models for Avar_Asia_Core1:

***Avar_Asia_Core1:***
100% *centralMongolia_preBA* (max P: 0.39, min P: 0.17, average P: 0.31)
100% *Fofonovo_EN* (max P: 0.32, min P: 0.13, average P: 0.26)

In 3 models Avar_Asia_Core1 formed a clade with *centralMongolia_preBA* (ERM003), while in other 3 models with *Fofonovo_EN* (FNO006). Although a few models from the competition runs indicated negative coefficients for a second ancestry, but these can be ignored due to significant nested-P values.

For Avar_Asia_Core2 we obtained 12 identical passing models:

***Avar_Asia_Core2:***
*100% Fofonovo_EN* (max P: 0.74, min P: 0.34, average P: 0.65) .

In each model Avar_Asia_Core2 formed a clade with *Fofonovo_EN* (FNO006).
Both *Fofonovo_EN* and *centralMongolia_preBA* genomes had been modelled as 83%–87% ANA, 12%–17% ANE (Botai)[2], thus all models are equivalent, and indicate that Avar_Asia_Core preserved very ancient Mongolian pre-Bronze Age genomes, with approximately 90% ANA ancestry, as also signified by 2-dimensional f4 statistics, PCA and ADMIXTURE.

**S3.7.5 proximodistal qpAdm:**

In the proximodistal qpAdm we tested 118 potential source populations, including 10 Bronze Age groups, 106 post-Bronze Age populations from the eastern Steppe, as well as modern Han and Nganasan. We used the same Right population set as in the distal analysis, supplemented with *Russia_UstIda_LN.SG* (Table S6b), as this population occurred in none of the 245 plausible distal models, but otherwise belongs to the same *Baikal_EN* group[23] as *Fofonovo_EN* with very similar ancestries, thus it is well suited to measure shared drift along Avar_Asia_Core ancestry components.

The two-source analysis resulted in 560 plausible models for Avar_Asia_Core1, and 955 for Avar_Asia_Core2. After model-competition we obtained 28 passing models for Avar_Asia_Core1, which were of the following types (Table S6b):

***Avar_Asia_Core1***
*95% UstBelaya_N.SG, 5% Steppe_IA* (max P: 0.88, min P: 0.07, average P: 0.77)
*58% Yana_Medieval.SG, 42% Ulaanzuukh/SlabGrave/lateXiongnu/*
(max P: 0.87, min P: 0.23, average P: 0.75)

The first model, which retained a distal source, represented 24/28 of the passing models, with a diverse list of Steppe populations as 2$^{nd}$ source, as 5% ancestry is insufficient for qpAdm to distinguish between similar sources. Just the second type of models can be regarded proximal.

For Avar_Asia_Core2 we obtained 39 comparable passing models which were of the following types (Table S6b):

***Avar_Asia_Core2***
*80-92% UstBelaya_N.SG, 8-20% Steppe_IA* (max P: 0.94, min P: 0.13, average P: 0.85)
*45-69% Xianbei_Hun_Berel_300CE, 31-55% Kazakhstan_Nomad_Hun_Sarmatian*
(max P: 0.97, min P: 0.26, average P: 0.9),

Again, the first model, which retained a distal source, represented 36/39 of the passing models, confirming that the two Avar_Asia_Core groups barely differ from each-other. On the other hand, the slightly higher Steppe_IA fraction in Avar_Asia_Core2 is in agreement with 2-dimensional f4-statistics, which indicated higher Iranian, Steppe and ANE ancestries in Avar_Asia_Core2. Again, just the second type of models can be regarded proximal. *UstBelaya_N*[34] carries predominantly ANA (DevilsCave) ancestry and is very similar to the proximal sources *Yana_Medieval*[34], *Ulaanzuukh/SlabGrave* and the distal source *Fofonovo_EN*[2]. The *Kazakhstan_Nomad_Hun_Sarmatian* also shows East Asian genetic descent[19] as well as *Xianbei_Hun_Berel_300CE*[4].
By all means, the proximate ancestors of Medieval Avars can be found among those late Xiongnus, Xianbeies and Hun_Sarmatians which mostly preserved their *Baikal_EN* ancestry.
Proximal qpAdm models also hint at similar genetic backgrounds of post-Xiongnu Hun-associated individuals and Avars.

The origin of the Avar elite is still debated and their Rouran ancestry is one plausible historical hypothesis[35]. Although the single low coverage Rouran genome[36] provided a poor fit in the qpAm models (Table S6b), our data are compatible with the Rouran origin of the Avar elite, as the Rouran genome is very similar to Avar_Asia_Core.

**S3.8 qpAdm modeling of Avar-cline members**

We analyzed the Conqueror- and Avar-cline individuals together. First, we presumed that as in the Conq-cline, most members of the Avar-cline also represent admixed progenies of immigrants with local people of the 7-9$^{th}$ century Carpathian Basin, which was tested by including just the EU_Core and Asia_Core sources in the Left populations (Table S6c).
From the 76 individuals in the cline 26 could be modelled as simple 2-way admixture between Eu_Core and Avar_Asia_Core (Table S6c, summarized in Table S1b). The rest required either a 3$^{rd}$ source, or entirely different sources, which was also tested in the same qpAdm experiment as shown for the Conqueror cline.

Five individuals could be modelled from EU_Core and Avar_Asia_Core plus a Hun-related 3$^{rd}$ source, two from an Iranian-related 3$^{rd}$ source, while one with Conq_Asia_Core 3$^{rd}$ source.
Two individuals at the Asian side of the cline did not contain EU_Core ancestry, and one (KK2-670) could be modelled from Avar_Asia_Core and Hun-related 2$^{nd}$ plus Iranian-related 3$^{rd}$ sources, while in case of KV-3456 the 3$^{rd}$ source was Scytho-Siberian/Conq_Asia_Core.
The Hun-related and Iranian-related sources had been defined in the Conq-cline section.

In 40 individuals qpAdm did not detect Avar_Asia_Core ancestry, so these samples can be regarded outliers. 28 of these outliers were modelled from EU_Core and Hun-related source with an Iranian-related 3$^{rd}$ source, while for 8 samples the 3$^{rd}$ source was Scytho-Siberian, Conq_Asia_Core, or Sarmatian. Two samples at the Asian extreme (SZOD1-187 and SZRV-266), which also clustered together with Hun_Asia_Core, did not show EU_Core ancestry, and was modeled from Hun_Asia_Core, plus a second Hun-related source and a 3$^{rd}$ Scytho-Iranian source (Table S6d).

Four cemeteries were represented by relatively large number of samples, which formed within-cemetery clines. From the Hortobágy-Árkus (ARK n=17), Szegvár-Oromdűlő (SZOD n=9), Makó-Mikócsa-halom (MM n=7) and Szarvas-Grexa (SZRV n=8) graveyards each individual could be modelled from genomes at the Asian and European extremes of the within-cemetery cline, revealing that immigrants of these communities had common origin (Table S6e). In these cases the genomes of the immigrants could be more accurately determined by unconstrained 3-source modeling of genomes at the Asian side (Table S6d), which detected a major Hun-related ancestry in each case.
In summary, Avar age samples were very heterogeneous, and their genomes could be derived from three major sources; approximately half of the population carried Avar_Asia_Core ancestry, which was most likely derived from the immigrants. The other than half of the Avar-cline individuals were outliers with Hun-related ancestries, and the best explanation for this finding is that the newcomers integrated remnants of the preceding European Hun empire on the Pontic Steppe, as also noted by historians[31]. A significant Iranian-related ancestry was universal in the Avar individuals, a clear sign of a third integrated population, which could derive from the Caucasus or/and the BMAC region. This ancestry could be present already in the Huns as well, as they had been recorded to have integrated the Alans. A few Avar individuals fell into the Conqueror-cline, and accordingly these were modelled from Conq_Asia_Core or Scytho-Siberians, pointing at the presence of other minor other elements among the Avars. We could not detect significant genetic differences between the three Avar period, indicating that the same populations entered the Carpathian Basin in multiple waves.

##### S3.9 Hun period samples

###### S3.9.1 PCA and unsupervised ADMIXTURE

Despite the low number of available Hun period samples, we could identify a Hun cline, which extends from the eastern end of the Avar-cline to the EU-cline (Fig. 2 and Extended Data Fig 1a). The MSG-1 sample maps somewhat below the eastern end of the Avar-cline, VZ-12673 overlaps with the eastern end of the Avar-cline, KMT-2785 maps between the Avar and Conqueror cline, ASZK-1, falls within the Conqueror cline, while the rest of the Hun Period samples (CSB-3, SEI-1, SEI-5, SEI-6, and SZLA-646) are members of the EU_Cline. Due to their diverse PCA positions we analyzed each Hun period samples separately.

The two samples at the Asian extreme of the cline, MSG-1 and VZ-12673 map close to each other on PCA and very likely represent immigrant genomes without European admixture. We must note that our VZ-12673 sample is identical to the Hungary_Hun_350CE (HUN001.merged) sample published in[4], which we resequenced with a higher coverage. The Admixture profile of MSG-1 shows 43% Nganasan, 40% Han, 12% Anat_N, 4% ANE and 1% EU_N components, while WHG and Iranian components are missing. The very similar VZ-12673 contains 40% Nganasan, 37% Han, 10% ANE, 9% Anat_N, 2% EU_N and 2% Iran_N components and no WHG (Fig. 2b and Table S4). On PC50 clustering MSG-1 and VZ-12673 fall in the same cluster, as they harbor the same ancestries (see below), therefore we termed them Hun_Asia_Core, but analyzed them separately.

###### S3.9.2 f3-statistics

Outgroup f3 statistics in the form f3(Mbuti; MSG-1, Y) and f3(Mbuti; VZ-12673, Y) gave comparable results, as out of the populations with top 50 f3 values, 41 were shared in the two statistics (Extended Data Fig 2c and 2d), indicating that MSG-1 and VZ-12673 had common ancestry. Both individuals shared highest drift with Neolithic farmers from the Wuzhuangguoliang site in northern China[3], earlyXiongnu_rest, Ulaanzukh and Slab Grave individuals[2], pointing to a likely Mongolian origin and Xiongnu affinity of these samples. It is notable that our Avar_Asia_Core1 group was the 16$^{th}$ in the top list of VZ-12673 and 35$^{th}$ in that of HUN-1, signifying common ancestry of European Huns and Avars. This relation is even more apparent if we compare the outgroup f3-statistics of Avars and Huns: out of the populations with top 50 f3 values of Avar_Asia_Core1, 41 are shared with VZ-12673 and 40 with MSG-1.

##### S3.9.3 distal qpAdm modeling of Hun_Asia_Core

In the distal modeling we tested 43 potential pre-Iron Age source populations from Mongolia and the surrounding regions (Table S5a) using the following Right populations:

*Ethiopia_4500BP.SG, China_Tianyuan, Iran_GanjDareh_N, Anatolia_N, Baikal_EN (Lokomotiv+Shamanka), DevilsCave_N.SG, WSHG (Tyumen+Sosnoviy), modern Onge and modern Koryak*

We obtained 32 plausible 2 source models for MSG-1 and 39 for VZ-12673, of which two models passed after model competition for MSG-1 and five for VZ-12673.

***MSG-1:***
*50% China_Wuzhuangguoliang_LN, 50% Mongolia_LBA_CenterWest_4D (I6351)*
(max P: 0,66, min P: 0,095, average P: 0,54)
***70% China_Wuzhuangguoliang_LN, 30% Mongolia_Khovsgol_LBA (ARS009.A0101)***
(max P: 0,32, min P: 0,056, average P: 0,23)

***VZ-12673:***
*88% Khovsgol_LBA (ARS009.A0101), 12% Russia_MN_Boisman (I3355, I1190)*
(max P: 0,67, min P: 0,068, average P: 0,51)
*93% China_Wuzhuangguoliang_LN, 7% Chemurchek_southAltai*
(max P: 0,61, min P: 0,17, average P: 0,45)
***88% China_Wuzhuangguoliang_LN, 12% Mongolia_Khovsgol_LBA (ARS026.A0101)***
(max P: 0,35, min P: 0,06, average P: 0,26)
*84% China_Wuzhuangguoliang_LN, 16% Altai_MLBA_o (I13173, KHI001.A0101)*
(max P: 0,35, min P: 0,06, average P: 0,26)
*94% China_Wuzhuangguoliang_LN, 6% Afanasievo_Mongolia*
(max P: 0,3, min P: 0,05, average P: 0,21)

The bolded models predict the same sources for the two individuals, thus demonstrate the same ancestry with slight differences between MSG-1 and VZ-12673, with somewhat higher Wuzhuangguoliang ancestry in the latter. Wuzhuangguoliang_LN was derived from a contact zone between the Central Plains and the North cultural regions, and have most similar genomes to present-day northern Han Chinese[3] predicting Han Chinese admixture in these individuals, which was typical in late Xiongnus[2]

##### S3.9.4 proximodistal qpAdm modeling of Hun_Asia_Core

In the proximodistal modeling we tested 93 sources, 7 from the Bronze Age, 86 from the post-Bronze Age, including Avar_Asia_Core and Conq_Asia_Core, as well as modern Han and Oroqen. We used the same Right population as in the distal analysis, except for *China_Tianyuan,* which was replaced with *China_Wuzhuangguoliang_LN* (Table S5b).

We obtained 829 plausible models for MSG-1, of which 88 passed after model-competition, which can be grouped into the following types:

***MSG-1:***
*72-88% lateXiongnu_han, 12-28% Scythian/Saka/Xiongnu*
(max P: 0.99, min P: 0.45, average P: 0.97)
*78-87% Kazakhstan_OutTianShanHun.SG, 13-22% Xiongnu/Han*
(max P: 0.98, min P: 0.39, average P: 0.93)
*50-76% Kazakhstan_Nomad_Hun_Sarmatian (DA27), 24-50% Scythian/Xianbei/Xiongnu*
(max P: 0.95, min P: 0.1, average P: 0.84)

56/88 of the top P-value models belonged to the first type, predicting a majority source of *lateXiongnu_han* (represented by samples KHO006.A0101, SON001.A0101, ATS001.A0101, BAM001.A0101) which had been modelled as 52% Ulaanzuukh_SlabGrave (ANA), 37% Han, 12% Chandman_IA[2].
The *Kazakhstan_OutTianShanHun* (DA127) sample is a genetic outlier, with enhanced East Asian features compared to its group members[19] and accordingly it clustered together with MSG-1, Xiongnu and Medieval Mongol samples on our PC50 Hierarchical Clustering (Table S3). The *Kazakhstan_Nomad_Hun_Sarmatian* (DA27) sample has similar enhanced East Asian features according to outgroup-f3 and PCA data[19], and it is mapped in the vicinity of MSG-1 on our PCA projection, yet latter models still require significant contribution from Scythian outlier (*Tasmola_Birlik_640BCE*[4]) or Xiongnu/Xianbei-like ancestries. The few other models with smallest P-values typically also contain late Xiongnu genomes.

For VZ-12673 we obtained 966 plausible models, of which 48 passed after model competition. The passing proximal models (with no Bronze Age sources) were mainly of the following types:

***VZ-12673:***
*63% lateXiongnu_han, 37% Xianbei (BRE011)*
(max P: 0.98, min P: 0.07, average P: 0.8)
*100% Kazakhstan_OutTianShanHun* (DA127)
(max P: 0.96, min P: 0.42, average P: 0.88)
*75-100% Kurayly_Hun_380CE, 11-25% Xiongnu/Avar_Asia_Core/Scythian outlier/Slabgrave/ Kazakhstan_Nomad_Hun_Sarmatian/Han*
(max P: 0.89, min P: 0.23, average P: 0.76)

In 5 models with top P-values VZ-12673 formed a clade with the same *Kazakhstan_OutTianShanHun* (DA127) sample which also occurred in the MSG-1 models, while in 19/48 of the next best models the majority ancestry was derived from the *Kurayly_Hun_380CE* (KRY001.A0101) sample, discovered in the Southern Ural. In around 1/3 of latter models *Kurayly_Hun_380CE* also formed a clade with VZ-12673. *Kurayly_Hun_380CE* had been shown to cluster together with *Hun_elite_350_CE* (=VZ-12673) and carry very similar ancestries[4]. On PCA *Kurayly_Hun_380CE* and *Kazakhstan_OutTianShanHun* map very close to VZ-12673 (Extended Data Fig 1a), and these 3 samples plus MSG-1 also tightly cluster together on our PC50 clustering (Table S3), thus these Hun individuals carry very similar genomes, despite of their significant geographical distances. For this reason, we added *Kurayly_Hun_380CE* and *Kazakhstan_OutTianShanHun* to the Hun_Asia_Core group (Fig. 2), without merging their sequences in the analysis.

In summary, our results indicate that ancestors of MSG-1 and VZ-12673 lived around present day Mongolia, and can be traced back to late Xiongnu ancestors which had significant Han admixture. Their high genetic affinity to other post-Xiongnu Hun genomes, *Kurayly_Hun_380CE*, *Kazakhstan_OutTianShanHun* and *Kazakhstan_Nomad_Hun_Sarmatian*[19] designates shared genetic ancestry of post-Xiongnu Huns. Besides, this genetic ancestry was also shared with Avar_Asia_Core.

##### S3.9.5 qpAdm modeling of Hun cline members

As samples KMT-2785 and ASZK-1 were in the middle of the genetic cline, we presumed that these could be admixed progenies of European and Asian ancestors. Therefore, in the Left populations of the proximal qpAdm analysis we included all EU_Core and Asia_Core groups, together with 98 additional post-Bronze Age populations. The following Right population set was used in the analysis (Table S5c):

*Ethiopia_4500BP, Iran_GanjDareh_N, Anatolia_N, EEHG (Karelia, Samara, Popovo), Latvia_HG, Baikal_EN (Lokomotiv+Shamanka), Russia_MLBA_Sintashta, modern Finnish, modern Han, modern Ket*

Out of the 63 plausible two-source models for KMT-2785, 15 passed after model-competition. Most passing models were of the following two types:

***KMT-2785:***
*67-78% LateXiongnu/Xianbei_Hun_Berel/Tasmola/Karakaba_830CE, 22-46% EU_Core*
(max P: 0.96, min P: 0.37, average P: 0.89)
*86% Russia_Sarmatian, 14% Mongolia_XiongNu*
(max P: 0.90, min P: 0.06, average P: 0.67)
*74% Russia_Sarmatian, 26% Kazakhstan_OutTianShanHun*
(max P: 0.69, min P: 0.13, average P: 0.56)

The LateXiongnu samples (UGU004.A0101, UGU011.A0101) occurred in 4/15 models, and these had the highest P-values. These LateXiongnu genomes had been modelled as 46-52% *Sarmatian* and 48-54% *Ulaanzuukh_SlabGrave*[2]. In the other models Sarmatians were the majority source, thus each model predicted a significant Sarmatian ancestry in KMT-2785, supplemented with Xiongnu/ Kazakhstan_OutTianShanHun. Latter sample formed a clade with VZ-12673 and was predicted to be a majority component of MSG-1 as well. In other words, ancestors of KMT-2785 may have derived from a Sarmatian-Xiongnu/Hun admixture, followed by a possible later admixture with local Europeans.

Out of the 673 plausible two-source models for ASZK-1, 83 passed after model-competition. The passing models were of two basic types:

***ASZK-1:***
*100% Russia_Sarmatian*
(max P: 0.88, min P: 0.11, average P: 0.76)
*52-74% Russia_IA, 26-48% LateXiongnu_sarmatian/ Sarmatian_450BCE/ /Kazakhstan_OutCentralSaka/ Kyrgyzstan_TianShanSaka*
(max P: 0.74, min P: 0.10, average P: 0.60)
*62-87% Russia_Late_Sarmatian, 13-38% LateXiongnu/ Korgantas_300BCE/ Sargat_300BCE*
(max P: 0.34, min P: 0.06, average P: 0.26)

In 75/83 of the top P-value models ASZK-1 formed a clade with the same *Russia_Sarmatian* group (DA145.SG, Pr10.SG), which appeared in the KMT-2785 passing models, indicating that major ancestry components of KMT-2785 and ASZK-1 could be shared in a large extent.
In the alternative models with lower P-values we also find significant Sarmatian ancestries complemented with late Xiongnu or comparable eastern Scythian genomes, derived from regions close to Mongolia (Tuva, Eastern Kazakhstan, Tian Shan).

In summary, KMT-2785 and ASZK-1 genomes indicate a majority Sarmatian ancestry, with a lower Xiongnu/Hun contribution. Similar genomes with a wide range of Sarmatian ancestry, including 100% in some of the the LateXiongnu_sarmatian individuals, had been reported from the Mongolian late Xiongnu period[2].

##### S3.9.6 qpAdm modeling Hun period EU-cline members

The rest of the Hun Period samples mapped to the northern part of the EU-cline, next to EU_Core5 samples and modern Hungarians. We performed proximal qpAdm modelling of these samples using 106 source populations, including our EU_Core groups, plus available post-Bronze Age groups from Europe and Asia, using the following Right populations (Table S5d):

*Ethiopia_4500BP, Iran_GanjDareh_N, Anatolia_N, Latvia_HG, Baikal_EN (Lokomotiv+Shamanka), Russia_MLBA_Sintashta, modern Finnish, and modern Ket*

For CSB-3 we obtained 463 plausible two-source models, of which 74 passed after model-competition. These were of the following 4 types:

***CSB-3:***
*57-93% EU_Core4/3, 7-43% Scytho-Siberian/Xiongnu*
(max P: 0.78, min P: 0.2, average P: 0.65)
*80-83% Ukraine_Chernyakhiv, 17-20% Russia_Alan/Anapa (own samples)*
(max P: 0.76, min P: 0.08, average P: 0.63)
*68-77% Czech_HallstattBylany, 23-32% Xiongnu/Kangju*
(max P: 0.69, min P: 0.06, average P: 0.54)
*71-85% Hungary_Scythian, 15-29% Xiongnu/Saka/Kangju*
(max P: 0.63, min P: 0.08, average P: 0.52)

64/74 of the models were of the first type, two of the second type, 6 of the third type, while two of the fourth type. Except for the *Chernyakhiv-Alan* models all other models can be regarded equivalent, as they indicate the admixture of a majority local population in the Carpathian Basin with immigrants from Eastern-Central Asia. The *Chernyakhiv-Alan* models, which had the second and fourth best P-values; however, predict a different story, with Goth and Alan ancestors from the Pontic Steppes. This individual obviously harbored a major European and a minor steppe ancestry, but qpAdm can poorly distinguish alternative models with similar minor components. Both alternatives are compatible with the known historical background, as Goths and Alans were part of the European Hun confederation.

For SEI-1 we obtained 417 plausible two-source models, of which 97 passed after model-competition.

***SEI-1:***
*70-76% EU_Core4, 24-30% Sarmatian*
(max P: 0.99, min P: 0.18, average P: 0.95)
*100% Ukraine_Chernyakhiv.SG*
(max P: 0.98, min P: 0.19, average P: 0.88)
*71-91% EU_Core/ Hungary_Scythian/ Czech_HallstattBylany/ Germany_EMedieval,*
*9-29% Xiongnu/Scytho Siberian*
(max P: 0.93, min P: 0.15, average P: 0.82)

The top 9 models with best P-values belonged to the first type, the next 6 top models to the second type, while the rest to the third type. The alternative models again predict two different scenarios, either a local ancestry admixed with Sarmatian/Xiongnu-like genomes, or this individual was 100% Goth. Latter case is supported by 12 alternative models in which 85-90% Early Medieval German ancestry was predicted, as Goth were a Germanic group.

For SEI-5 we obtained 476 plausible two-source models, of which 83 passed after model-competition.

***SEI-5:***
*66-77% EU_Core4/3, 23-34% Sarmatian*
(max P: 0.99, min P: 0.22, average P: 0.90)
*74-94% EU_Core/ Germany_EMedieval/ Hungary_Scythian/ Czech_HallstattBylany,*
*6-26% Xiongnu/Scytho Siberian*
(max P: 0.78, min P: 0.25, average P: 0.67)
*100% Ukraine_Chernyakhiv*
(max P: 0.79, min P: 0.05, average P: 0.66)

The top 16 models were of the first type, the rest of the second type, while the weakest 4 models the third type, often with negative coefficients. SEI-1 and SEI-5 are in close proximity on PCA, thus are expected to give similar models, and their best models fulfill this criteria.

For SEI-6 we obtained 212 plausible two-source models, of which 6 passed after model-competition.

***SEI-6:***
*100% Ukraine_Chernyakhiv*
(max P: 0.91, min P: 0.30, average P: 0.80)
*76-81% Czech_HallstattBylany, 19-24% Russia_Alan/Anapa*
(max P: 0.69, min P: 0.18, average P: 0.58)
*80-84% Germany_EMedieval, 16-20% EU_Core1/Italy_Imperial*
(max P: 0.5, min P: 0.09, average P: 0.4)

The two top P-value models were of the first type, the weakest two of the last type. The dominant Germanic components in four of these models indicate a likely Gothic ancestry of this individual.

We did not obtain valid models for SZLA-646, indicating that sources of this individual were not present in the Left populations. As outgroup f3-statistics indicated Baltic and Germanic ancestors of SZLA-646 (data not shown), we rerun qpAdm with a different Left and Right population set (Table S5e). Right population:

*Ethiopia_4500BP.SG, Anatolia_N, Latvia_HG, WSHG (Tyumen+Sosnoviy), Russia_MLBA_Sintashta, Sweden_HG_Motala, Czech_CordedWare and modern Finnish*

From the 103 plausible models 42 passed, and in the top 21 models SZLA-646 formed a clade with *Lithuania_Late_Antiquity*[19] or *England_Saxon*[37] samples (Table S5e).

***SZLA-646:***
*100% Lithuania_Late_Antiquity*
(max P: 0.99, min P: 0.8, average P: 0.95)
*100% England_Saxon*
(max P: 0.97, min P: 0.44, average P: 0.9)

This result also shows northern European affinity, so the SZLA-646 woman could be also part of a Germanic group which joined the Huns.

In summary, our Hun period samples with majority European ancestries were either Goths/Germans, or local individuals with Xiongnu-like or Sarmatian-like minor ancestries.
Overall our results from the Hun period samples are compatible with the Xiongnu origin of the core population of European Huns, which was extensively admixed with Sarmatians and Germanic groups. These results are fully in line with the historical sources as well.

##### S3.10 samples from the Caucasus

##### S3.10.1 PCA and ADMIXTURE

We have sequenced 3 samples derived from the Caucasus region. Kinship analysis indicated that Anapa-10 and Anapa-11 were first degree relatives, probably brothers, as they had identical maternal (J1c) and paternal (R1a1a1b1a2b3a~) haplogroups, while the Anapa-9 female was a more distant relative of both, with a different (U5a2a1) maternal lineage (Table S1a).

The ADMIXTURE profile of the Anapa-11 individual was 33% Iranian, 32% EU_N, 19% ANE, 8% Anat_N, 6% Han 3% Nganasan, and no WHG (Table S4).
On PCA Anapa individuals map among modern samples from the Caucasus (Fig. 2), and they cluster together with ancient samples from the Caucasus and BMAC regions (Table S3). According to outgroup f3 statistics in the form f3(Mbuti; Anapa, Y) Anapa samples had highest shared drift with European Neolithic and Bronze Age, Russia_Early_Sarmatian, Russia_Caucasus_LBA_Dolmen, and Russia_Srubnaya samples (data not shown).
These data indicated a deep-rooted European Neolithic ancestry, with subsequent Iranian and Steppe admixture, and latter also included some east Eurasian (ANA) ancestry according to ADMIXTURE.

##### S3.10.2 qpAdm analysis

We merged the 3 relatives for this analysis. To clarify the possible origin of the high Iranian element in Anapa we assembled 60 potential source populations with significant Iranian elements, also including several Steppe populations and populations with Steppe-Iranian admixture (Table S5f). We used the following Right population set:

*Ethiopia_4500BP, ANE (AfontovaGora3+ Russia_MA1), Iran_GanjDareh_N, Georgia_Kotias, Anatolia_N, Latvia_HG, Romania_Mesolithic_IronGates, Baikal_EN (Lokomotiv+Shamanka), Russia_Steppe_Catacomb, modern Armenian*

We obtained 27 plausible two source models, and all of these indicated a 74-97% majority component derived from the Caucasus region *Armenia_MBA*[38], *Russia_Alan*[19], *Armenia_Caucasus_EBA_KuraAraxes*[39], or *Armenia_LchashenMetsamor*[19]) supplemented with eastern steppe genomes derived from Kazakhstan, Mongolia, Kyrgyzstan or Turkmenistan. Model competition resulted in 10 passing models, exactly the 10 highest P-value models from the initial analysis:

***Anapa:***
*76-96% Armenia_MBA.SG, 4-24% Hun_Sarmatian/TianShanHun/TianShanSaka/ CentralSteppe_EMBA/Xiongnu/Wusun/Turk*
(max P: 0.84, min P: 0.63, average P: 0.78)

Armenia_MBA was the majority source in each model, and these results indicated that Anapa individuals carried ancient genomes derived from the Caucasus region, admixed with Steppe genomes containing east Eurasian elements.

To clarify the Steppe source we run the next qpAdm analysis only with the best sources from the Caucasus, supplemented with 55 Steppe source populations (Table S5g). We used the same Right populations as before, except for *ANE* and *Romania_Mesolithic_IronGates* which were removed, because they poorly rejected models according to the log files of the first qpAdm experiment.

We obtained 216 plausible two source models, and 58 valid models after model-competition. The valid models were of the following two types:

***Anapa:***
91-96% *Armenia_MBA.SG, 4-9% late Xiongnu/Scytho-Siberian/Hun_Sarmatian/ Avar_Asia_Core/Conq_Asia_Core/Karluk/Turk/*
(max P: 0.88, min P: 0.25, average P: 0.77)
*100% Russia_SaltovoMayaki.SG*
(max P: 0.70, min P: 0.06, average P: 0.58)

In the first 26 models with top P-values *Armenia_MBA* was the majority source, indicating that the best proximal sources are missing from the available dataset. According to the second type of models, with lower P-values, *Russia_SaltovoMayaki*[19] formed a clade with our Anapa samples. Latter model needs to be interpreted with caution, as the three available *Russia_SaltovoMayaki* genomes have low coverage (0.029, 0.04 and 0.072), furthermore, despite the presence of *Russia_SaltovoMayaki* in the Left populations, it did not appear among the plausible models of the first qpAdm run. At the same time the close proximity of Saltovo-Mayaki samples on PCA as well as their PC50 clustering together suggest that proximal sources of the Anapa samples could be similar to Saltovo-Mayaki, which was contemporary to Anapa, and both could be part of the Khazar Khaganate.

##### S3.11 Neolithic samples

We have sequenced three new Neolithic genomes from Hungary from three different cemeteries. Two individuals represented the Hungary_Tisza Neolithic culture, from the Vésztő-Mágori-halom and Hódmezővásárhely-Gorzsa sites, and one individual the Hungary_Starcevo early Neolithic culture, from the Alsónyék-Bátaszék, Mérnöki telep site. Other genomes had been published previously from each site[40], and our genomes do not show any difference from these. They have the same ADMIXTURE profile (Table S4), PCA location (Fig. 2), and cluster together with Anatolian and European farmers on our PC50 clustering (Table S3). The G2a2 Y-chromosome haplogroups as well as J1c2 and H maternal lineages (Table S1a) also fit this conclusion as these were frequent in Neolithic Europe[40].

**S3.12 Y-chromosome and mtDNA results**

The distribution of uniparental markers along the PCA genetic clines shows a general rule: at the Asian side of the cline we find individuals with Asian haplogroups (Hgs), while at the European side individuals carry European Hgs. Along the cline from the Asian side towards the European side the same trend prevails, decreasing frequency of Asian and increasing of European Hgs. The few exceptions from this rule are nearly always detected in admixed individuals between EU_Core and Asia_Core (Table S1a), nevertheless, several individuals in the EU-cline carry Asian Hgs, testifying distant Asian forefathers. This is especially prominent in the Jánoshida-Tótkérpuszta (JHT) Late Avar graveyard, where all three males carried R1a1a1b2a (R1a-Z94) Asian Y-Hg, in spite of their European genomes. The Middle Avar MT-17 individual from Madaras-Téglavető, who also mapped in the EU-cline, also carried R1a-Z94, though in this cemetery all three other males carried N1a1a1a1a3a and had Asian genomes.

Both Hun_Asia_Core individuals (VZ-12673 and MSG-1) carry R1a-Z94, as well as ASZK-1 from the Hun-cline. The other two published genomes united in Hun_Asia_Core, *Kurayly_Hun_380CE* (KRY001.A0101)[4] and *Kazakhstan_OutTianShanHun* (DA127)[19] carry Hgs R1a-Z94 and Q respectively, indicating that these Hgs could be common among the Huns, very likely inherited from Xiongnus[2]. Considering all published post-Xiongnu Hun era genomes (Hun period nomad, Hun-Sarmatian, Tian Shan Hun[19] and Xianbei-Hun Berel[4], we counted 10/23 R1a-Z93 and 9/23 Q Hgs, affirming above observation. These Y-Hgs were most likely inherited from Xiongnus, among whom high frequency of these Hgs had been found[41]. The rest of our Hun period samples with European genomes carried derivatives of R1a1a1b1, a Hg typical in North-Western Europe, in line with the Germanic affinity of many of these samples, detected with qpAdm.

From the 9 Avar_Asia_Core males 7 carried the N1a1a1a1a3a (N1a-F4205) Y-Hg, one C2a1a1b1b and one R1a1a1b~ (very likely R1a-Z94) confirming, that N1a-F4205, most prevalent in modern Chukchis and Buryats[42], was also prevailing among the Avar elite as shown before[22,43]. This Hg was indeed common in members of the Avar-cline, and also seems to cluster in certain cemeteries. In the Ároktő (ACG), Felgyő (FU), Szegvár-Oromdűlő (SZOD), Csepel-Kavicsbánya (CS), Kiskőrös-Vágóhídi dűlő (KV), Kunpeszér-Felsőpeszér (KFP), Csólyospálos-Felsőpálos (CSPF), Kiskundorozsma-Kettőshatár II (KK2), Tatárszentgyörgy (TTSZ), Madaras-Téglavető (MT) Avar graveyards all, or the majority of males carried the N1a-F4205 Hg, mostly accompanied with Asian maternal lineages. These cemeteries must have belonged to the immigrant Avar population, while the local population seems to have separated, as many Avar period cemeteries show no sign of Asian ancestry. Latter include Mélykút-Sándordűlő (MS), Szeged-Fehértó A (SZF), Szeged-Kundomb (SZK), Szeged-Makkoserdő (SZM), Kiskundorozsma-Kettőshatár I (KK1), Kiskundorozsma-Daruhalom (KDA), Orosháza-Bónum Téglagyár (OBT), Székkutas-Kápolnadűlő (SZKT), Homokmégy-Halom (HH), Alattyán-Tulát (ALT), Kiskőrös-Pohibuj Mackó dűlő (KPM), and Sükösd-Ságod (SSD), in which Asian lineages barely occur. In the SZK, ALT, KK1, OBT, SZKT, HH and SZM cemeteries most males belonged to the E1b1b1a1b1 (E-V13) Hg, which is most prevalent in the Balkan[44], and accordingly many of the samples from these cemeteries fell in EU_Core1, or its vicinity, with typical Southern European genomes.

There is a third group of Avar period cemeteries representing immigrants from Asia, but with a different genetic background. In males from Makó-Mikócsa-Halom (MM), Dunavecse-Kovacsos dűlő (DK), Árkus Homokbánya (ARK), and Szarvas-Grexa (SZRV) Y-Hgs R1a-Z94 and Q1a2a1 dominated, which were typical in European Huns, and were accompanied with mostly Asian maternal lineages. These Avar period peoples could have represented Hun remnants, which joined the Avars but isolated in separate communities. These inferences are also supported by genomic data, as most qpAdm models from these cemeteries indicated the presence of Hun_Asia_Core or Xiongnu ancestries (Table S1b). As indicated before, Hun ancestry was also present in several other cemeteries, like Hg R1a-Z94, but in those cemeteries the population was less uniform.

The Conqueror population had a more heterogeneous Hg composition than the Huns and Avars. In the 6 males of Conq_Asia_Core we detected three N1a1a1a1a2~, one D1a1a1a1b, one C2a1a1b1b and one Q1a1a1 Y-Hg, generally accompanied by Asian maternal lineages. Two other N1a1a1a1a2a1c~ Y-Hgs were detected in the SO-5 Conqueror elite and the PLE-95 commoner individuals, thus this Hg was specific for the Conqueror group. Obviously this Hg links Conquerors with Mansis, as had been shown before[45]. Another related Y-Hg, N1a1a1a1a4 (M2128) was detected in two Conqueror elite samples from present study, as well as from other two Conqueror elite samples in our previous study[22]. This Hg is typical for modern Yakuts, and occurs with lower frequency among Khantys, Mansis and Kazakhs[42], thus may also link Conquerors and Mansis, though it was also present in one Middle Avar individual. It is notable, that the European Y-Hg I2a1a2b1a1a was also specific for the Conqueror group, especially for the elite, as also shown before[22], very often along with Asian maternal lineages, indicating that this Hg could be more typical for the immigrants than to the local population. Additionally, two other Y-Hgs appeared with notable frequencies among the Conquerors, R1a-Z94 was present in 3 elite and 2 commoner individuals, while Hg Q was carried by 3 elite individuals, which may be sign of Hun relations, also detected at the genome level.

#### S4 Genetic References:

11. Amorim, C. E. G. *et al.* Understanding 6th-century barbarian social organization and migration through paleogenomics. *Nat. Commun. 2018 91* **9**, 1–11 (2018).
12. Antonio, M. L. *et al.* Ancient Rome: A genetic crossroads of Europe and the Mediterranean. *Science* **366**, 708–714 (2019).
13. Lazaridis, I. *et al.* Genetic origins of the Minoans and Mycenaeans. *Nature* **548**, 214–218 (2017).
14. Allentoft, M. E. *et al.* Population genomics of Bronze Age Eurasia. *Nature* **522**, 167–172 (2015).
15. Mathieson, I. *et al.* Genome-wide patterns of selection in 230 ancient Eurasians. *Nature* **528**, 499–503 (2015).
16. Olalde, I. *et al.* The Beaker phenomenon and the genomic transformation of northwest Europe. *Nat. 2018 5557695* **555**, 190–196 (2018).
17. Gamba, C. *et al.* Genome flux and stasis in a five millennium transect of European prehistory. *Nat. Commun.* **5**, 5257 (2014).
18. Veeramah, K. R. *et al.* Population genomic analysis of elongated skulls reveals extensive female-biased immigration in Early Medieval Bavaria. *Proc. Natl. Acad. Sci. U. S. A.* **115**, 3494–3499 (2018).
19. De Barros Damgaard, P. *et al.* 137 ancient human genomes from across the Eurasian steppes. *Nature* **557**, 369–374 (2018).
20. Jeong, C. *et al.* The genetic history of admixture across inner Eurasia. *Nat. Ecol. Evol.* **3**, 966–976 (2019).
21. Neparáczki, E. *et al.* Mitogenomic data indicate admixture components of Central-Inner Asian and Srubnaya origin in the conquering Hungarians. *PLoS One* (2018) doi:10.1371/journal.pone.0205920.
22. Neparáczki, E. *et al.* Y-chromosome haplogroups from Hun, Avar and conquering Hungarian period nomadic people of the Carpathian Basin. *Sci. Rep.* **9**, (2019).
23. De Barros Damgaard, P. *et al.* The first horse herders and the impact of early Bronze Age steppe expansions into Asia. *Science (80-. ).* (2018) doi:10.1126/science.aar7711.
24. Róna-Tas, A. *Hungarians and Europe in the Early Middle Ages: An Introduction to Early Hungarian History*. (Central European University Press, 1999).
25. Abondolo, D. M. *The Uralic languages*. (Routledge, 1998).
26. Järve, M. *et al.* Shifts in the Genetic Landscape of the Western Eurasian Steppe Associated with the Beginning and End of the Scythian Dominance. *Curr. Biol.* **29**, 2430-2441.e10 (2019).
27. Krzewińska, M. *et al.* Ancient genomes suggest the eastern Pontic-Caspian steppe as the source of western Iron Age nomads. *Sci. Adv.* **4**, (2018).
28. Lazaridis, I. *et al.* Ancient human genomes suggest three ancestral populations for present-day Europeans. *Nature* **513**, 409–413 (2014).
29. Mallick, S. *et al.* The Simons Genome Diversity Project: 300 genomes from 142 diverse populations. *Nat. 2016 5387624* **538**, 201–206 (2016).
30. Moorjani, P. *et al.* A genetic method for dating ancient genomes provides a direct estimate of human generation interval in the last 45,000 years. *Proc. Natl. Acad. Sci.* **113**, 5652–5657 (2016).
31. Kim, H. J. *The Huns, Rome and the Birth of Europe*. *The Huns, Rome and the Birth of Europe* (Cambridge University Press, 2013). doi:https://doi.org/10.1017/CBO9780511920493.
32. Csáky, V. *et al.* Early medieval genetic data from Ural region evaluated in the light of archaeological evidence of ancient Hungarians. *Sci. Reports 2020 101* **10**, 1–14 (2020).
33. Maár, K. *et al.* Maternal Lineages from 10–11th Century Commoner Cemeteries of the Carpathian Basin. *Genes 2021, Vol. 12, Page 460* **12**, 460 (2021).
34. Sikora, M. *et al.* The population history of northeastern Siberia since the Pleistocene. *Nat.*

*2019 5707760* **570**, 182–188 (2019).
35. Golden, P. B. Some Notes on the Avars and Rouran. in *The Steppe Lands and the World Beyond Them: Studies in Honor of Victor Spinei on His 70th Birthday* (eds. Curta, F. & Maleon, B.-P.) 43–66 (Editura Universitatii 'Alexandru Ioan Cuza', 2013).
36. Li, J. *et al.* The genome of an ancient Rouran individual reveals an important paternal lineage in the Donghu population. *Am. J. Phys. Anthropol.* **166**, 895–905 (2018).
37. Schiffels, S. *et al.* Iron Age and Anglo-Saxon genomes from East England reveal British migration history. *Nat. Commun. 2016 71* **7**, 1–9 (2016).
38. Lazaridis, I. *et al.* Genomic insights into the origin of farming in the ancient Near East. *Nature* **536**, (2016).
39. Wang, C. C. *et al.* Ancient human genome-wide data from a 3000-year interval in the Caucasus corresponds with eco-geographic regions. *Nat. Commun. 2019 101* **10**, 1–13 (2019).
40. Lipson, M. *et al.* Parallel palaeogenomic transects reveal complex genetic history of early European farmers. *Nature* (2017) doi:10.1038/nature24476.
41. Keyser, C. *et al.* Genetic evidence suggests a sense of family, parity and conquest in the Xiongnu Iron Age nomads of Mongolia. *Hum. Genet.* **140**, 349–359 (2021).
42. Ilumäe, A. M. *et al.* Human Y Chromosome Haplogroup N: A Non-trivial Time-Resolved Phylogeography that Cuts across Language Families. *Am. J. Hum. Genet.* **99**, 163–173 (2016).
43. Csáky, V. *et al.* Genetic insights into the social organisation of the Avar period elite in the 7th century AD Carpathian Basin. *Sci. Reports 2020 101* **10**, 1–14 (2020).
44. Cruciani, F. *et al.* Tracing past human male movements in northern/eastern Africa and western Eurasia: New clues from Y-chromosomal haplogroups E-M78 and J-M12. *Mol. Biol. Evol.* **24**, 1300–1311 (2007).
45. Post, H. *et al.* Y-chromosomal connection between Hungarians and geographically distant populations of the Ural Mountain region and West Siberia. *Sci. Reports 2019 91* **9**, 1–10 (2019).
